# ATP Regeneration from Pyruvate in the PURE System

**DOI:** 10.1101/2024.09.06.611674

**Authors:** Surendra Yadav, Alexander J. P. Perkins, Sahan B. W. Liyanagedera, Anthony Bougas, Nadanai Laohakunakorn

## Abstract

The ‘Protein synthesis Using Recombinant Elements’ (‘PURE’) system is a minimal biochemical system capable of carrying out cell-free protein synthesis using defined enzymatic components. This study extends PURE by integrating an ATP regeneration system based on pyruvate oxidase, acetate kinase, and catalase. The new pathway generates acetyl phosphate from pyruvate, phosphate, and oxygen, which is used to rephosphorylate ATP in situ. Successful ATP regeneration requires a high initial concentration of *∼* 10 mM phosphate buffer, which surprisingly does not affect the protein synthesis activity of PURE. The pathway can function independently or in combination with the existing creatine-based system in PURE; the combined system produces up to 233 *µ*g/ml of mCherry, an enhancement of 78% compared to using the creatine system alone. The results are reproducible across multiple batches of homemade PURE, and importantly also generalise to commercial systems such as PURExpress^®^ from New England Biolabs. These results demonstrate a rational bottom-up approach to engineering PURE, paving the way for applications in cell-free synthetic biology and synthetic cell construction.

## I. INTRODUCTION

Cell-free protein synthesis (CFPS) harnesses the core biological processes of transcription and translation to generate mRNA and protein from a DNA template, in controlled biochemical reactions outside of living cells. These systems were originally used to elucidate fundamental mechanisms in molecular biology, but recently have been deployed for a wide range of applications within synthetic biology and biotechnology [1]. These include practical applications such as biomanufacturing, biosensing, and diagnostics [2], as well as more fundamental research involving bottom-up construction of biomolecular systems which aim to mimic aspects of living cells [1].

CFPS reactions are composed of three categories of components: 1) a DNA template encoding for a gene of interest; 2) small-molecule substrates and cofactors such as nucleotide triphosphates (NTPs), amino acids, and tRNA; and 3) the molecular machinery needed to carry out in vitro protein synthesis, which includes enzymes, translation factors, and ribosomes.

This molecular machinery is supplied either in the form of clarified cell lysates, which can be obtained from a number of different organisms [3–6]; or they can be recombinantly produced and individually purified. Such a defined system, known as PURE (Protein synthesis Using Recombinant Elements) was originally developed by Shimizu and coworkers in 2001 [7], and consists of purified *Escherichia coli* ribosomes and 36 protein factors, which together carry out transcription, translation, tRNA aminoacylation, and biochemical energy regeneration, the four processes needed to sustain cell-free protein synthesis.

Since the PURE system is defined, it has been used in a number of studies which can take advantage of this, such as unnatural amino acid incorporation and in vitro directed evolution [8]. It is also an ideal starting point for the construction of more complex subsystems which may eventually be combined into a replicating, autonomous synthetic cell [9–11]. However, these ambitious applications are challenged by the fact that the system in its current form is limited in both protein synthesis yield (for example, *∼* 200 *µ*g/ml green fluorescent protein, GFP) and reaction lifetime (*∼* 1 hour) [12]. The limitations of PURE include low processivity and speed of translation leading to truncation, inactive products, and stalled ribosomes [13, 14]; protein misfolding [15]; tRNA and translation factor depletion; and accumulation of inhibitory misfolded mRNA [16] and inorganic phosphate [14].

To an extent these limitations are shared with lysate-based systems, which have been more intensively studied. Early work in the 2000s focused on engineering *E. coli* lysates to improve the performance of batch-mode CFPS reactions, and a number of limitations were discovered and addressed [17]. It was recognised that a major limitation was the buildup of inorganic phosphate (P_i_) as the reaction proceeds, resulting from ATP hydrolysis; this phosphate chelates and reduces the concentration of free Mg^2+^ ions in the system. Magnesium is a cofactor for many enzymes involved in protein synthesis, and additionally stabilises the structure of the ribosome [18], and so a reduction in free Mg^2+^ due to phosphate accumulation was proposed as a cause of batch reaction termination after only *∼* 1 hour [19].

Since CFPS is a very energy intensive process, consuming *∼* 4 *−* 5 ATP equivalents per peptide bond synthesized (BioNumbers BNID 107782, [20]), its biochemical energy must be replenished to allow the reaction to proceed beyond a few minutes in vitro. Simple ATP regeneration schemes involving substrate-level phosphorylation, using enzyme/substrate pairs such as pyruvate kinase/phosphenolpyruvate [21], acetate kinase/acetyl phosphate [22], or creatine kinase/creatine phosphate [23], result in a unidirectional transfer of phosphate to ATP, which then accumulates following ATP hydrolysis.

A solution to this problem is to either remove the accumulated phosphate through physical means (e.g. continuous exchange [24, 25] or flow reactors [26, 27]), or alternatively to engineer a biochemical regeneration scheme which recycles phosphate. The first such demonstration was by Kim and Swartz who supplemented *E. coli* lysates with pyruvate oxidase from *Pediococcus* sp. [28]. Pyruvate oxidase catalyzes the condensation of pyruvate and inorganic phosphate to produce acetyl phosphate, which regenerates ATP through its conversion to acetate; the second reaction is catalyzed using endogenous acetate kinase already present in the extract. This system produces the phosphorylated substrate acetyl phosphate in situ, while avoiding phosphate accumulation, and the authors showed that both the reaction yield and lifetime were increased. Current state-of-the-art systems also benefit from phosphate recycling, such as the maltose/maltodextrin-based ‘TXTL’ systems developed by Noireaux and coworkers which condense maltose with phosphate to form glucose-1-phosphate, which is subsequently fed into the glycolytic pathway to regenerate ATP. Such systems are able to achieve up to 4 mg/ml of GFP, with reaction lifetimes up to 20 h [29, 30].

Unlike lysates, PURE systems typically use a creatine phosphate/creatine kinase (CP/CK) energy regeneration scheme, and hence are proposed to suffer from phosphate accumulation [14] (Figure 1a). We hypothesized that implementing a similar phosphate recycling scheme to that developed by Kim and Swartz should directly address the limitations of reaction yield and lifetime in PURE (Figure 1b).

**FIG. 1.**
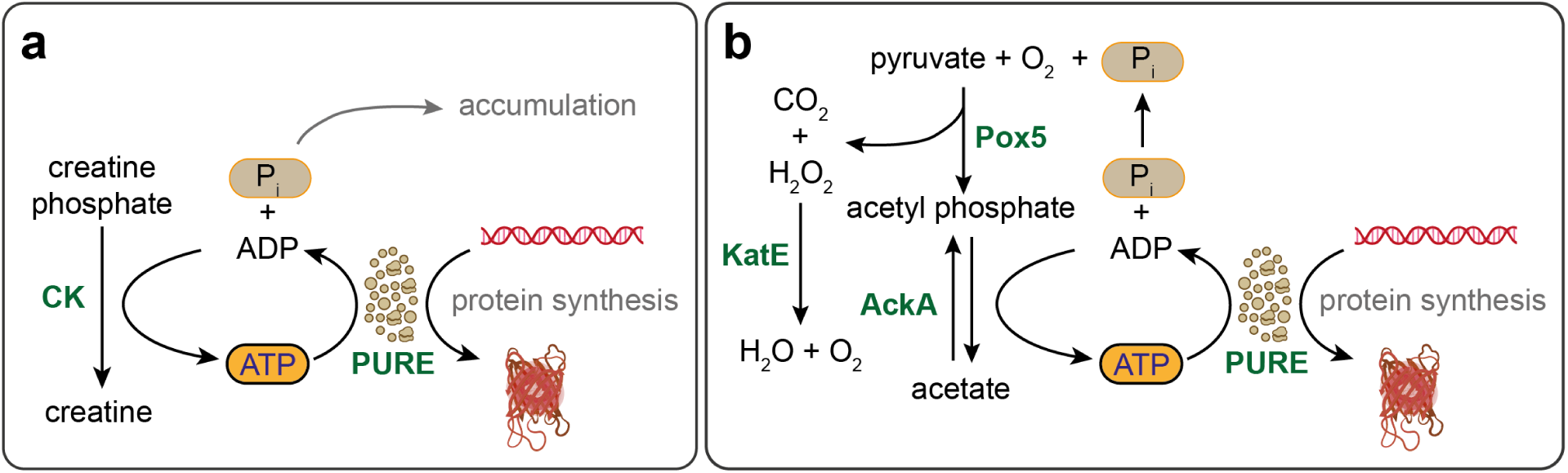
Schematic of CP/CK **(a)**, and PAP **(b)** as ATP regeneration pathways in the PURE system. **(a)** CP/CK employs creatine phosphate as a substrate to regenerate ATP from ADP, utilizing the enzyme creatine kinase (CK). This process results in the accumulation of inorganic phosphate (P*_i_*) as a byproduct. **(b)** PAP utilizes pyruvate, inorganic phosphate, and molecular oxygen as substrates to regenerate ATP, mediated by the enzymes pyruvate oxidase (Pox5) and acetate kinase (AckA). Additionally, the enzyme catalase (KatE) is involved in the breakdown of hydrogen peroxide, a byproduct of the Pox5 reaction, into water and molecular oxygen.

In this work we demonstrate the production and purification of three enzymes: *Lactobacillus plantarum* pyruvate oxidase (Pox5), *E. coli* acetate kinase (AckA), and *E. coli* monofunctional catalase (KatE). We supplement the enzymes into a PURE system produced in-house, and demonstrate that the new ‘pyruvate-acetate pathway’ (‘PAP’) can energize the PURE system using pyruvate. We apply a design of experiments approach to rationally improve the performance of the pathway, and find that the optimised pathway can power the synthesis of 72 *µ*g/ml of fluorescent protein mCherry over 4 h. Compared to the existing creatine phosphate/creatine kinase system, which produces *∼* 130 *µ*g/ml, this pathway is not an effective replacement. However the two pathways in combination can produce up to 230 *µ*g/ml of mCherry after 4 h. Importantly, the results generalize when the pathway is supplemented into commercial PURExpress^®^ systems (New England Biolabs).

This work demonstrates that alternative energy regeneration schemes can be straightforwardly implemented in combination with the PURE system. These results as well as other similar studies combining PURE with synthetic metabolic systems [31–33] pave the way for rationally improving the PURE system’s performance through bottom-up construction.

## II. RESULTS

### A. *L. plantarum* pyruvate oxidase can be recombinantly expressed in *E. coli*

Genes (pox5 from *L. plantarum*, codon optimized for *E. coli* using the IDT Codon Optimization Tool; ackA and katE from *E. coli*) encoding the three pathway enzymes were obtained as linear DNA fragments (gBlocks, IDT) and individually cloned into the pET21a expression vector. A 6x-His tag was incorporated at the C-terminus of each enzyme to facilitate affinity purification [34]. The enzymes AckA and KatE were then overexpressed in BL21(DE3) cells using conventional IPTG induction at 37*^◦^*C for 3 hours [35]. However, during the heterologous expression of Pox5 in *E. coli* under similar conditions, the protein was found exclusively in the insoluble fraction of the cellular lysate (Figure S1, Supporting Information). To address this, we implemented a modified overexpression protocol by lowering the expression temperature to 15°C and extending the induction period to 15 hours [36]. This adjustment successfully facilitated the extraction of a fraction of the enzyme within the soluble lysate (Figure S1, Supporting Information). After overexpression, the enzymes were purified using Ni-NTA affinity chromatography [34], and their activities were verified through individual enzymatic assays (Figure S2–S4, Supporting Information).

### B. Phosphate buffer activates PAP without inhibiting protein synthesis in PURE

Initially, PAP was tested by adding the following components into a CFPS reaction: the pathway enzymes (Pox5, AckA, and KatE); two co-factors for Pox5, thiamine pyrophosphate (TPP) and flavin adenine dinucleotide (FAD); the PURE system; pyruvate; and a modified energy solution. In order to ensure energy regeneration occurs solely via PAP and not the existing CP/CK system, we omitted creatine phosphate from the energy solution formulation (we verified that the alternative approach, of removing the creatine kinase enzyme from the PURE system, also gave the same results, Figure S5, Supporting Information). Phosphate was not added as a substrate initially as we expected the pathway to utilize the inorganic phosphate generated from cell-free protein synthesis through the consumption of ATP.

This approach initially led to markedly low mCherry protein yields (*<* 25 *µ*g/ml) (Figure 2a). We hypothesized that the diminished protein yield stemmed from low levels of inorganic phosphate within the reaction, which limits the substrate availability for the Pox5 enzyme [37]. To address this, we tested the addition of two different phosphate sources in the reaction, monobasic potassium phosphate (KH_2_PO_4_, pH 4.5) and potassium phosphate buffer (a mixture of monobasic and dibasic potassium phosphate K_2_HPO_4_, pH 7). Remarkably, supplementing with 10 mM potassium phosphate buffer significantly increased CFPS output, whereas supplementation with monobasic potassium phosphate inhibited protein synthesis (Figure 2a).

**FIG. 2.**
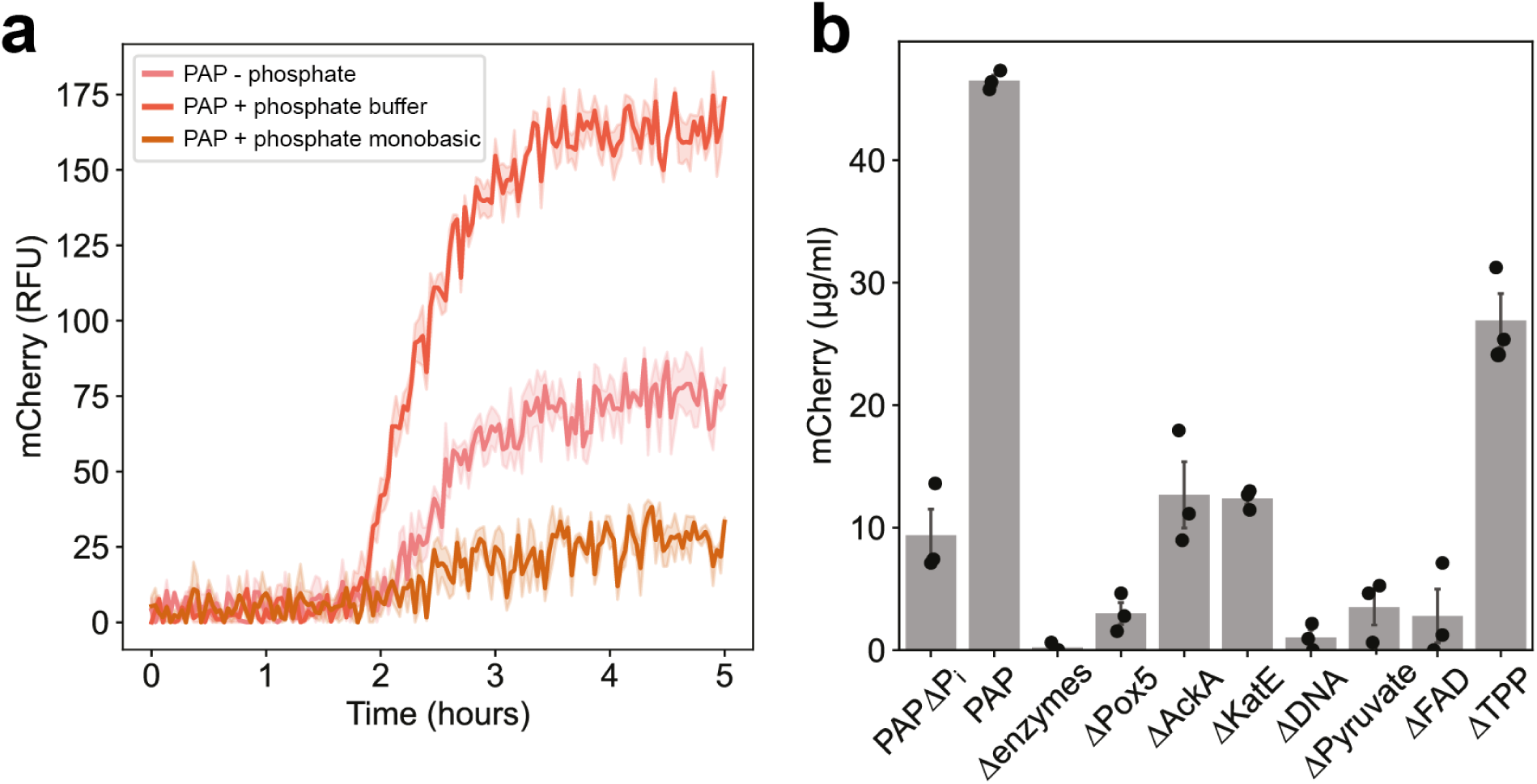
(**a**) PAP functioned effectively as an ATP regeneration pathway in the PURE system when it was supplemented with 10 mM exogenous phosphate in the form of a potassium phosphate buffer at pH 7. In contrast, 10 mM monobasic potassium phosphate inhibited the reactions. **(b)** Final protein synthesis yields from PAP-powered reactions after 5 hours are reported. Protein synthesis did not occur in the absence of the mCherry DNA template (ΔDNA) or the pathway enzymes (Δenzymes). Pyruvate, Pox5 and FAD are critically important for the pathway’s function, while the pathway retained some activity with the exclusion of phosphate, AckA, and KatE. The Pox5 cofactor TPP is the least essential component, indicating that some TPP might remain bound to the enzyme after purification. All experiments were performed in triplicate. Data are shown as mean *±* s.e. (n = 3).

To verify this effect and further characterize these observations, we carried out a titration of the two phosphate sources in PURE reactions using the original CP/CK energy regeneration system. We observed that the addition of monobasic potassium phosphate at concentrations *≥*15 mM inhibited protein synthesis, whereas the supplementation of phosphate buffer up to 20 mM had no inhibitory impact on the reactions (Figure S6–S7, Supporting Information).

We next carried out experiments using PAP+PURE with the exclusion of various components (Figure 2b; Figure S8, Supporting Information). We observed that the exclusion of pathway enzymes and DNA effectively abolished protein synthesis activity. Similarly, the exclusion of pyruvate, Pox5 or FAD led to low levels of protein synthesis. However, we observed that the exclusion of TPP from PAP still yielded 26.9 *±* 2.2 *µ*g/ml of mCherry protein. This finding is consistent with the hypothesis that TPP binds tightly to the Pox5 enzyme, and a fraction of TPP remains bound to the enzyme even after the protein purification process is completed [37], resulting in the enzyme being partially active in the absence of exogenously added TPP.

The KatE enzyme was found to be crucial for the efficient functioning of the pathway, as its exclusion resulted in a significant decrease in protein yields (Figure 2b). The absence of KatE may lead to the accumulation of hydrogen peroxide within the reaction, which can directly oxidize protein thiol groups, particularly cysteine residues, and initiate further radical-mediated damage. This oxidation process can cause protein dysfunction, misfolding, and aggregation, ultimately resulting in a loss of protein synthesis activity [38, 39]. Furthermore, we observed that protein synthesis activity persisted at a low level even when the AckA enzyme, the sole enzyme responsible for ATP regeneration in the system, was excluded (Figure 2b). We hypothesize that this residual activity may be attributed to trace amounts of co-purified AckA present within the PURE protein mixture, as AckA is an endogenous enzyme of *E. coli* [28].

With the supplementation of 10 mM potassium phosphate buffer, PURE reactions using the PAP energy regeneration scheme achieved a final protein yield of 46.5 *±* 0.5 *µ*g/ml, a five-fold increase compared to phosphate-free reactions (Figure 2b). Encouraged by these enhancements, we decided to adopt a design of experiments (DOE) approach for pathway optimization, with the aim of further increasing protein yields using PAP.

### C. PAP can be optimised using a design of experiments (DOE) approach

To enhance the performance of the pathway, we first adjusted the initial concentrations of key components pyruvate, phosphate buffer, and magnesium glutamate, using a rational design of experiments (DOE) approach. We utilized a circumscribed central-composite design (cCCD) to efficiently explore the design space and examine the interactions between these factors (Figure 3a) [40]. This choice was informed by previous findings that demonstrated a nonlinear relationship between Mg^2+^ and protein yield in CFPS reactions, as well as the known interaction between Mg^2+^ and phosphate [41]. The experimental design space was defined by setting maximum and minimum levels for each factor based on plausible ranges encountered in the literature (Figure 3b) [28, 42–44].

**FIG. 3.**
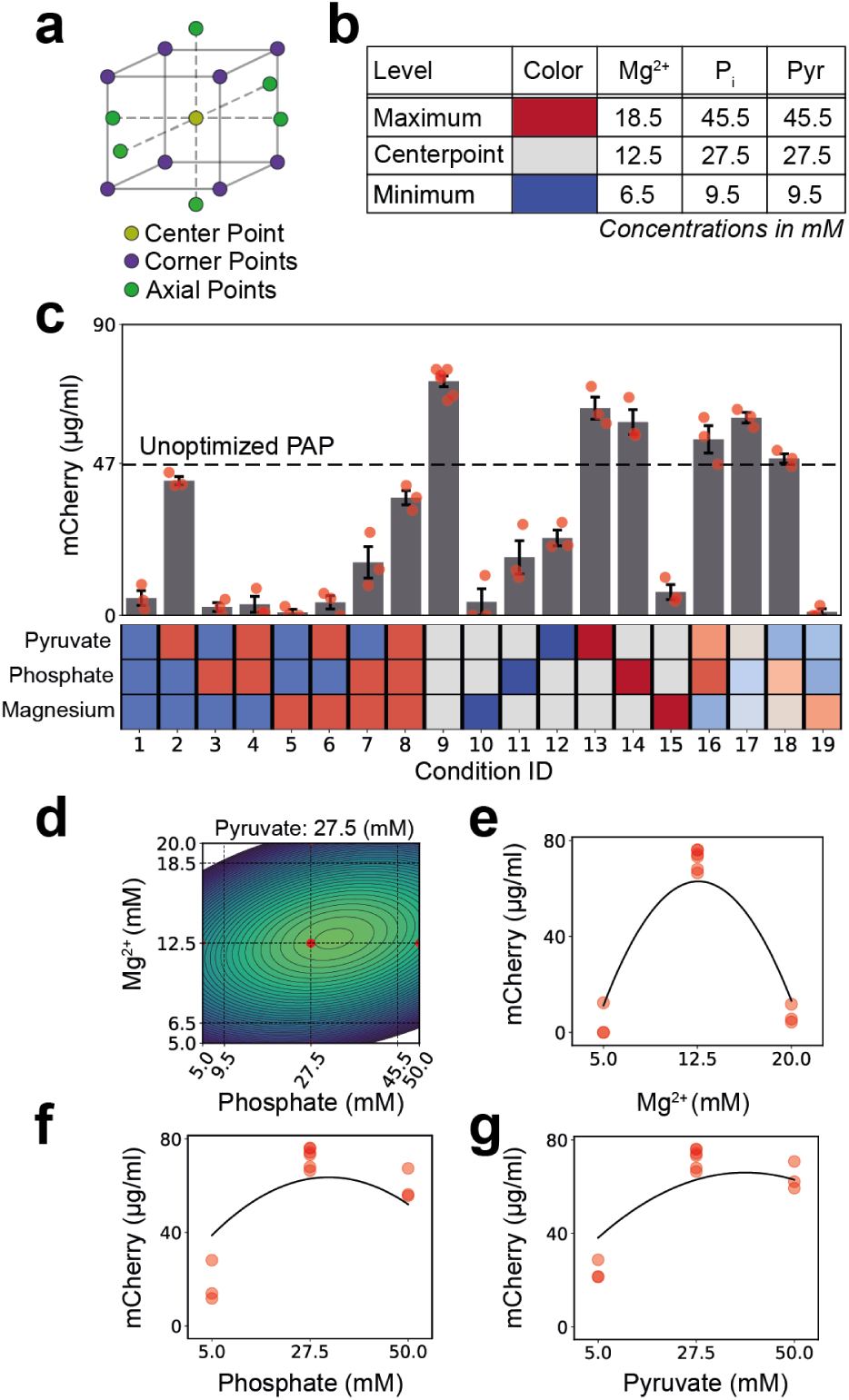
Optimization of PAP using a design of experiments approach. **(a)** Illustration of the geometric relationship between the different elements within the circumscribed central composite design (cCCD). **(b)** Table of the design space boundaries with the maximum, minimum and centerpoint concentrations for each factor. **(c)** Mean and standard error of the mCherry yield after 5 hours, for each condition tested. Individual data points are plotted as red markers, and the assigned levels for each condition are visualised in the heat map below. The mCherry yield of the unoptimized system is plotted as a dashed line at 46.5 *µ*g/ml. **(d)** Contour plot of the model predicted yield between magnesium and phosphate, where pyruvate is fixed at 27.5 mM. Experimental data points are plotted as red markers with size scaled by yield. **(e, f, g)** Plots of the mCherry protein yield as a function of each factor, with the others fixed at their centerpoint levels: experimental data are indicated with red markers, and the model predictions with the solid line.

Initial examination of the data (Figure 3c, Table S1, and Figure S9, Supporting Information) showed that six conditions had an enhanced performance compared to the unoptimized system, with the best performing condition corresponding to the center point (Mg^2+^ = 12.5 mM, phosphate = 27.5 mM, and pyruvate = 27.5 mM). This condition yielded 74.5 *±* 0.8 *µ*g/ml of mCherry, an increase over the baseline of 60.2%. Notably, the worst performing conditions corresponded to high or low Mg^2+^ values, highlighting the sensitivity of the reaction to Mg^2+^ concentration.

In order to probe the behaviour of the system more quantitatively, we carried out a response surface analysis by fitting an unsaturated linear regression model to the data, which included linear and quadratic terms for each factor, and a single interaction term between Mg^2+^ and phosphate,

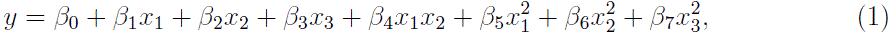

where the variables *x*_1_, *x*_2_, and *x*_3_ correspond to magnesium, phosphate, and pyruvate concentrations respectively, and *y* the yield of mCherry. The final fitted model parameters are given in Table S2, Supporting Information; this yielded *R*^2^ = 0.83 with respect to the cCCD training data, and *R*^2^ = 0.42 on held-out validation data outside the original dataset, which suggests a moderate degree of overfitting, although the model predictions are more accurate at higher expression levels (Figure S10, Supporting Information). The contributions of model terms to predicting the response can be quantified using a t-statistic (Figure S11, Supporting Information)[45], which shows that the Mg^2+^ concentration is the most sensitive factor, in agreement with the initial qualitative observations. The Mg^2+^-phosphate interaction is also significant, while the quadratic terms are negligible in importance.

The overall response can be visualised in Figure 3d–g, which shows a sharp optimum dominated by Mg^2+^ concentration, with weaker dependencies on pyruvate and phosphate. The interaction between Mg^2+^ and phosphate is illustrated in the contour plot in Figure 3d, which shows that the Mg^2+^ optimum varies weakly as a function of phosphate concentration.

### D. Combining ATP regeneration from PAP and CP/CK improves protein yield in PURE

The DOE-optimized pathway resulted in a yield of 74.5 *±* 0.8 *µ*g/ml of mCherry. Compared to the CP/CK system alone, which produces 130.9 *±* 8.0 *µ*g/ml, the PAP system does not serve as an effective replacement (Figure 4a, orange and blue curves); however, we hypothesised that the two pathways in combination might achieve a higher total yield.

**FIG. 4.**
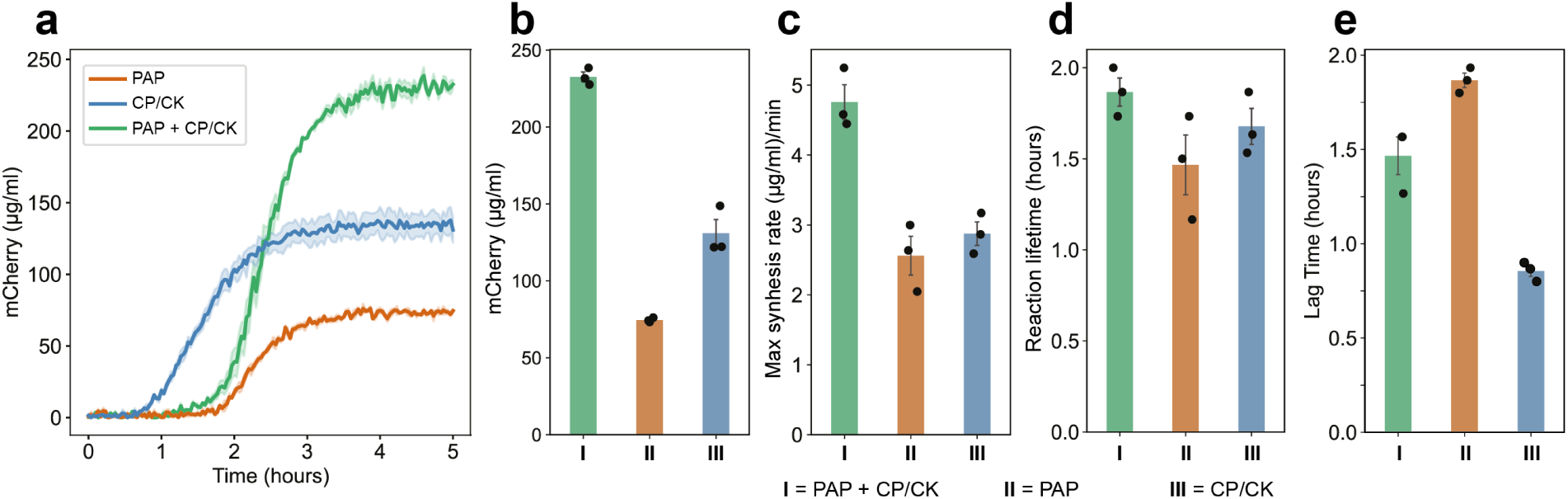
The combination of the PAP and CP/CK pathways significantly increased both the rate of protein synthesis and the final protein yield. **(a)** Protein expression kinetics of PURE reactions utilizing different ATP regeneration pathways are shown. **(b)** Final protein synthesis yields after 5 hours. PURE reactions utilizing the CP/CK system gave a higher protein yield compared to those using optimized PAP; however the combined PAP+CP/CK system exceeds either system alone. **(c)** Maximal protein synthesis rates are shown for each system. Similar to the yield, the combined PAP+CP/CK system exhibited the highest maximal protein synthesis rates. **(d)** The reaction lifetime remained approximately constant in all three cases. **(e)** Reactions containing PAP exhibit increased lag time compared to the CP/CK reaction. All experiments were performed in triplicates. Data are shown as mean ±s.e. (n = 3).

To accomplish this, the final reaction mixture was supplemented with both creatine phosphate and the PAP components (Table S3, Supporting Information). This setup endowed the PURE system with two ATP regeneration pathways, CP/CK and PAP, each utilizing different substrates. The combination of both systems led to a significant increase in protein yield up to 232.6 *±* 3.2 *µ*g/ml of mCherry, marking a 77.6% enhancement compared to using only the CP/CK system, and 212% increment compared to using only PAP (Figure 4a and 4b). Concurrently with the increased yield, the combined system also shows an increase in the maximal rate of protein synthesis (Figure 4c).

### E. PAP-powered reactions exhibit increased lag time for fluorescent protein production

We additionally assessed the kinetics of protein expression by analysing two timescales associated with the reaction. The first is the lag time, defined as the time interval from the start of the reaction until the first appearance of a fluorescence signal: this is the minimal time for synthesis and maturation of the first fluorescent proteins. The second measure is the reaction lifetime, which we defined as the time interval between the first appearance of signal until the saturation of the fluorescence at its maximal value. This is equal to the time interval over which protein synthesis remains active. Details of this analysis are given in Figure S12, Supporting Information.

Our original hypothesis that the reaction lifetime would be increased under the PAP system was not proven, as this interval remained roughly constant between the CP/CK and PAP reactions (Figure 4d). However, we observe that PAP-powered reactions exhibit an increased lag time before the first appearance of mCherry fluorescence (*∼* 1.5h), compared to CP/CK-powered reactions (*<* 1h) (Figure 4e). This could be due to a number of reasons, including a delayed start to transcription and/or translation, as well as a delay in the maturation of the fluorescent protein.

Further analysis of the DOE dataset revealed that pyruvate concentration was correlated with lag time (Figure S13, Supporting Information): the greater the initial concentration of pyruvate, the longer the delay. Our first hypothesis was that an increase in pyruvate would result in increased H_2_O_2_ production, which would be inhibitory to CFPS. This should be compensated by the addition of additional catalase; however a titration of catalase did not reveal any effect on lag time (Figure S14, Supporting Information).

Limited availability of acetyl phosphate, and hence ATP, can also be ruled out as an explanation for the increased lag time, as under that hypothesis the lag should decrease with increasing pyruvate. Additionally, the increased lag is present in the combined CP/CK+PAP system, where ATP availability is assured from the CP/CK pathway. Thus it is more likely that it is the fluorescent protein maturation, rather than CFPS, which is responsible for the increased lag.

Since the pyruvate oxidase reaction is oxygen-dependent, our working hypothesis therefore is that a lowered concentration of dissolved oxygen brought about by Pox5 activity could delay mCherry maturation [46]. This anaerobic state would be maintained as long as pyruvate is available, but would cease as soon as pyruvate runs out. Hence increasing initial pyruvate concentrations would increase the lag time. A rough quantitative estimate using average parameters obtained from BRENDA (pyruvate oxidase EC 1.2.3.3, *k_cat_ ∼* 20/s, *K_M,pyr_* = 0.4 mM, *K_M,P i_* = 2.3 mM) predicts that the total conversion of pyruvate under our initial conditions should take of the order of 20 minutes, consistent with the observed increase in lag time.

### F. PAP can power the commercial PURExpress^®^ system

Finally, we assessed the reproducibility of the PAP across different batches of PURE produced in-house, as well as the commercially available PURExpress^®^ system. The pathway exhibited consistent performance across different batches of homemade PURE, yielding comparable final protein yields (Figure S15, Supporting Information). To test PAP as an ATP regeneration component in a commercial PURE system, the PURExpress^®^ ΔRibosome Kit from New England Biolabs was used. The experiments were conducted using only the ‘Factor Mix’ from the kit as it contained all the PURE proteins. All other reaction components were homemade, including ribosomes, energy solution, mCherry DNA template, and other required additives. Upon addition to PURExpress^®^ without creatine phosphate, PAP by itself resulted in increased protein yield (159.9 *±* 4.5 *µ*g/ml), surpassing the protein yield under the CP/CK system (113.7 *±* 3.8 *µ*g/ml). Furthermore, combining PAP with the CP/CK system in PURExpress^®^ resulted in protein yields exceeding 350.0 *µ*g/ml, showing 207.8% enhancement compared to PURExpress^®^ utilizing only the CP/CK component, and 118.9% increment compared to only using the PAP (Figure 5a and 5b). Broadly similar trends to the in-house PURE data were observed with synthesis rates and lag times, although the reaction lifetime appeared marginally increased under the dual energy system (Figure 5c–e).

**FIG. 5.**
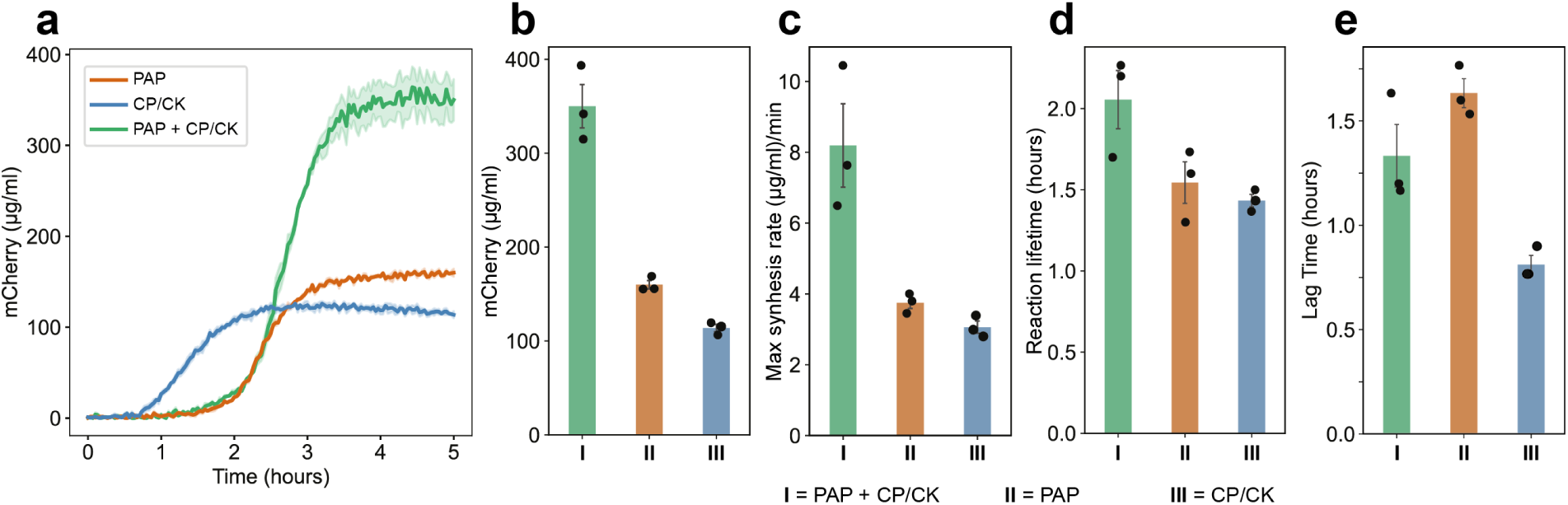
PAP functions as an ATP regeneration component in the commercial PURExpress^®^ system. Only the ‘Factor Mix’ from the PURExpress^®^Δ Ribosome Kit was used in the experiments as it contained all the PURE proteins. All other reaction components were homemade, including ribosomes, energy solution, mCherry DNA template, and other required additives. **(a)** Protein expression kinetics of ‘Factor Mix’ reactions utilizing different ATP regeneration pathways are shown. **(b)** Final protein synthesis yields after 5 hours are shown. ‘Factor Mix’ reactions utilizing optimized PAP provided a higher protein yield compared to those using CP/CK. Similar to previous experiments, the combined system gave the highest yield. **(c)** ‘Factor Mix’ reactions utilizing both optimized PAP and CP/CK achieved the highest protein synthesis rates, compared to each pathway alone. **(d)** The reaction lifetime of the combined system is marginally greater than for the individual systems. **(e)** Reactions containing PAP exhibit increased lag time compared to the CP/CK reaction. All experiments were performed in triplicates. Data are shown as mean ±s.e. (n = 3).

## III. DISCUSSION

In this work, we demonstrated that parallel energy regeneration systems can be straightforwardly integrated with PURE. While the original CP/CK system regenerates sufficient ATP to power protein synthesis, it has been hypothesised to be self-limiting due to inorganic phosphate accumulation [14]. We tested a pathway which regenerates ATP concurrently with phosphate recycling, as originally proposed by Kim and Swartz [28]. Since the pathway now relies on phosphate as a substrate, our initial observation that the pathway resulted in low synthesis yields was unsurprising, as the only source of inorganic phosphate came from ATP hydrolysis.

What was surprising however was the finding that 20 mM of phosphate buffer can activate the pathway, as such phosphate concentrations were originally suggested to be inhibitory to the PURE system in particular. Li et al. measured up to 20 mM of phosphate accumulation in PURE, and additionally showed that replenishing the reaction at later times with magnesium acetate increased CFPS activity, which they attributed to the mitigation of magnesium sequestration [14]. However, our finding that 20 mM of phosphate buffer does not inhibit PURE reactions is at odds with the earlier results.

As is well known, the protonation state of the phosphate anion is pH dependent: at pH 7, phosphate mainly exists in the monobasic 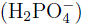 and dibasic forms 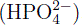, while at pH 4.5, it is only the monobasic form that is predominant. The dissociation constant of Mg^2+^ with dibasic phosphate is estimated to be *K_d_ ∼* 10–60 mM [47–49]. Thus it is possible that magnesium sequestration would only be significant at higher P*_i_* concentrations than the 20 mM or so which accumulates in PURE.

These observations are mirrored in studies using lysate systems: supplementation of CFPS reactions with up to 10 mM phosphate is necessary to activate glucose metabolism in lysates, as demonstrated by Calhoun et al. [50]. Importantly, they find inhibitory concentrations of dibasic phosphate are high—up to 50 mM.

The combination of multiple ATP regeneration pathways has been demonstrated before in lysates, for example with combined CP/CK and glucose metabolism [19]. In our work we observe that the combination of PAP and CP/CK leads to roughly an additive increase in both final protein yield and maximum synthesis rate. This suggests that the pathways are independently contributing to the ATP regeneration, possibly by maintaining a higher steady-state ATP level throughout the CFPS process. Time-resolved measurements of ATP level throughout the reactions could verify this hypothesis.

In light of these findings, we conclude that while PAP functions effectively as an ATP regeneration pathway, the contribution of its phosphate recycling mechanism towards increasing reaction yield is unlikely to be significant under the conditions tested. However, PAP may enable the operation of multiple pathways whose combined effects would otherwise result in high (*>* 50 mM) levels of phosphate accumulation, and we plan to test these conditions in future work.

We propose that the inhibition observed upon addition of 20 mM monobasic phosphate is likely due to lowered pH rather than magnesium sequestration: it is well known that lowered pH inhibits CFPS [51]. To support this, we measured the initial pH of reactions supplemented with either monobasic phosphate or phosphate buffer. We found that increasing concentrations of monobasic phosphate lowered the pH of the reactions from 6.95 to 6.54. In contrast, increasing concentrations of phosphate buffer had negligible effects on the pH of the reactions (Figure S16, Supporting Information).

Finally, while PAP demonstrates the feasibility of introducing a parallel, independent ATP regeneration system in PURE, there are a few weaknesses associated with the design. The oxygen requirement of pyruvate oxidase imposes limitations on scaling up the reactions beyond *∼* 50-100 *µ*l, while the enzyme’s oxygen-consuming activity delays fluorescent protein maturation and hence limits kinetic monitoring. The buildup of acetate eventually lowers the pH of the reactions, and can be a limiting factor. A potential alternative pathway to explore is based on pyruvate dehydrogenase and phosphate acetyltransferase, which could generate acetyl phosphate from pyruvate, using coenzyme-A and NAD^+^ as cofactors.

## IV. CONCLUSION

In conclusion, we have demonstrated the construction of an ATP regeneration system based on pyruvate oxidase, acetate kinase, and catalase, and its integration into the PURE cell-free protein synthesis system. This PAP pathway utilises pyruvate and phosphate as substrates, generating an acetyl phosphate intermediate that rephosphorylates ATP in situ. The pathway itself can function independently, producing up to 74.5 *±* 0.8 *µ*g/ml of mCherry, or it could be combined with the existing creatine phosphate/creatine kinase system in PURE to produce 232.6 *±* 3.2 *µ*g/ml of mCherry. This behaviour is reproducible across multiple batches of homemade PURE, and generalizable to the commerical PURExpress^®^ system. This demonstration indicates the relative ease and flexibility of constructing parallel metabolic pathways in PURE, which should enable future applications in cell-free protein synthesis and the construction of synthetic cells.

## V. MATERIALS AND METHODS

Brief descriptions of methods are given here; for fully detailed procedures please see the ‘Experimental Details’ section in the Supporting Information.

### Materials

All materials, including chemicals and reagents used in this study, are listed in the Supporting Information (Table S4), complete with their respective supplier catalog numbers.

### *E. coli* strains and plasmids

*E. coli* TOP10 was used for plasmid maxiprep, *E. coli* DH5*α* was used for plasmid maintenance, and *E. coli* BL21(DE3) was used for protein expression and ribosome purification (Table S5, Supporting Information). Plasmids encoding PURE proteins used in this study were gifts from Sebastian Maerkl and Takuya Ueda (Addgene plasmids #124103-124138), except pET21a-MTF-6xHis. Genes coding for MTF-6xHis, Pox5-6xHis, KatE-6xHis, and AckA-6xHis were obtained as linear DNA fragments (gBlocks, IDT) (Table S6, Supporting Information), cloned into a pET21a vector using the Gibson Assembly^®^ Cloning Kit (NEB) according to the manufacturer’s protocol [52], and then transformed into *E. coli* BL21(DE3) and DH5*α* cells. The gene coding for mCherry-6xHis was cloned into a T7p14 vector, and then transformed into *E. coli* TOP10 and BL21(DE3) cells. A list of PURE proteins along with their corresponding vector, gene, and strain are provided in Table S7, Supporting Information. The primers used in this study are provided in Table S8, Supporting Information. Plasmid maps and amino acid sequences of all expressed proteins are given in Table S9–S10, Supporting Information.

### PURE proteins preparation

The PURE proteins were prepared using a modified ‘OnePot’ protocol based on established procedures [12]. Briefly, strains for the PURE system were grown from standardised liquid glycerol stocks to an optical density of OD600=2-3, followed by co-inoculation of all 36 strains into 500 ml of LB media with ampicillin. Cultures were incubated at 37°C, with shaking at 220 RPM until OD600=0.2-0.3, at which point protein expression was induced using a final concentration of 0.1 mM IPTG. Following a 3-hour expression period, cells were harvested and washed with PBS and flash frozen in liquid nitrogen. The cell pellets were thawed and re-suspended in a lysis buffer, and lysed by sonication. Lysates were clarified at 15,923g and incubated with Ni-NTA resin for 3 hours at 4°C. Protein purification involved a single 5 mM immidazole wash step followed by elution with 450 mM immidazole. Purified proteins were dialyzed, concentrated, and adjusted to a final concentration of 12.5 mg/ml in 30% glycerol buffer. The protein solution was aliquoted and stored at –80°C until use. PUREΔCK systems were created in the same way but with the omission of the creatine kinase strain in the main culture.

### Crude ribosome preparation

Crude 70S ribosomes used in the CFPS reactions were purified using high-speed zonal centrifugation as described previously [53]. Ribosomes were purified from *E. coli* BL21(DE3) cell cultures harvested at an optical density of OD600=0.6. Briefly, a starter culture was inoculated from a glycerol stock and incubated, with a parallel sample to test for media contamination. The culture was then scaled up to 4 *×* 750 ml through successive stages, with optical density monitoring to guide the growth to the desired cell density. Following growth, cells were cooled, harvested by centrifugation, and washed in PBS to remove media. Cell lysis was achieved via sonication, and the lysates were clarified through an initial centrifugation at 30,000g for 1 hour at 4°C. Clarified lysate was subsequently ultra-centrifuged at 100,000g for 4 hours at 4°C to obtain crude ribosome pellets. These pellets were then processed through several re-suspension and centrifugation steps using high salt buffer to further purify the ribosomes. Finally, the ribosome concentration was determined using Nanodrop spectrophotometry, and the ribosome solutions were aliquoted and stored at –80°C until use.

### Buffers used for protein and ribosome purification

Buffers used for protein and ribosome purification are listed in Table S11–S12, Supporting Information. All buffers were filter-sterilized using bottle-top filters with a 0.2 µm PES membrane, and stored at 4°C until use. Reducing agent TCEP (for proteins) or DTT (for ribosomes) was added immediately before use.

### Plasmid maxiprep of T7p14-mCherry-6xHis

Plasmid DNA was purified using a modified ZymoPURE II Plasmid Maxiprep kit protocol, adjusted to process the equivalent of three standard reactions to accommodate higher plasmid quantities. The protocol commenced with inoculation of a single colony or glycerol stock into LB media with ampicillin, followed by 500 ml overnight culture. Post-culture, cells were harvested and subjected to a lysis-clearing-neutralization sequence using ZymoPURE^™^ reagents P1, P2, and P3. Following lysis, cell debris was removed using ZymoPURE^™^ Syringe Filter-X units, ensuring critical attention to avoid sample loss and ensure the recovery of 70-100 ml of cleared lysate. The lysate was then mixed with ZymoPURE^™^ Binding Buffer, and the DNA-bound mixture was processed through Zymo-Spin^™^ V-PX Column assemblies with subsequent wash steps to remove contaminants. DNA was eluted in 500 *µ*l of Nuclease-Free Water, pre-heated to 55°C. This elution was further purified for endotoxins using EndoZero^™^ Spin-Columns, with elutions combined for final DNA concentration and purity assessment via Nanodrop spectrophometry, targeting a yield of around 900 ng/*µ*l.

### Energy solution preparation

The energy solution for PURE reactions was prepared as previously described [12], with several modifications. A 4x energy solution, excluding creatine phosphate, magnesium glutamate, and potassium glutamate, was prepared. This solution contained 200 mM HEPES, 8 mM ATP, 8 mM GTP, 4 mM CTP, 4 mM UTP, 14 mg/ml tRNA, 4 mM TCEP, 0.08 mM folinic acid, 8 mM spermidine, and 1.2 mM of each amino acid (with the exception of leucine, which was at 1 mM) [12]. For more information, refer to Table S13, Supporting Information.

### Reaction setup for cell-free protein synthesis

PURE reactions utilizing only the CP/CK ATP regeneration system were prepared in a 50 *µ*l master mix containing 12.5 *µ*l of 4x Energy Solution, 20 mM creatine phosphate, 2.4 mg/ml PURE Protein solution, 2.275 *µ*M ribosome solution, 11.8 mM Mg-glutamate, 100 mM K-glutamate, 2% PEG 8K, and 10 nM mCherry plasmid DNA template. Other reaction components such as phosphate (if added) varied according to the reaction requirement. PURE reactions utilizing both CP/CK and PAP systems were prepared in a 50 *µ*l master mix containing 12.5 *µ*l of 4x Energy Solution, 20 mM creatine phosphate, 2.4 mg/ml PURE Protein solution, 2.275 *µ*M ribosome solution, 12.5 mM Mg-glutamate, 100 mM K-glutamate, 2% PEG 8K, 27.5 mM K-phosphate buffer (pH 7), 27.5 mM pyruvate, 3.03 *µ*M Pox5, 13.89 *µ*M AckA, 2.38 *µ*M KatE, 2 mM TPP, 0.2 mM FAD and 10 nM mCherry plasmid DNA template. PURE reactions utilizing only the PAP system were prepared in a 50 *µ*l master mix containing 12.5 *µ*l of 4x Energy Solution, 2.4 mg/ml PURE Protein solution, 2.275 *µ*M ribosome solution, 100 mM K-glutamate, 2% PEG 8K, 13.89 *µ*M AckA, 2 mM TPP, 0.2 mM FAD, and 10 nM mCherry DNA template. All other reaction components, including enzymes, substrates, etc. varied according to the reaction setup. Detailed reaction setups are provided in Table S3, Supporting Information. PURExpress^®^ reactions (50 *µ*l master mix) utilizing CP/CK, or PAP, or both components were set up in the same way as mentioned above, except the PURE protein solution was replaced with 6 *µ*l PURExpress^®^ Factor Mix. PURExpress^®^ control reaction (50 *µ*l master mix) contained 20 *µ*l Solution A, 6 *µ*l Factor Mix, 9 *µ*l NEB ribosomes, and 10 nM mCherry DNA template (Figure S17, Supporting Information). All the reaction master mixes were prepared on ice, mixed using pipetting and split into three 15 *µ*l reactions in a 384-well plate (Greiner 384 *µ*Clear Black). The protein synthesis kinetics of reactions were then measured on a BioTek Synergy H1 plate reader (excitation, 579 nm; emission, 616 nm; gain 50; bottom optics; double orbital 2 sec shaking before every read), taking fluorescence readings every 2 mins for 5 hours. Relative fluorescence units were converted to physical concentration units of active fluorescent protein using a calibration curve (Figure S18, Supporting Information).

### SDS-PAGE gels

Samples for SDS-PAGE were prepared by mixing 10 *µ*l of the protein sample with 12.5 *µ*l of 4x Laemmli buffer (Bio-Rad), 5 *µ*l of 100 mM DTT, and 22.5 *µ*l of H_2_O, followed by boiling for 10 minutes. Using Bio-Rad 4-20% Mini-PROTEAN Precast Gels, the prepared samples, including 5 *µ*l of the ladder and 10 *µ*l of each sample, were loaded into wells. Electrophoresis was conducted at 180 volts for 40 minutes in a Mini-PROTEAN Tetra Vertical Electrophoresis Cell. The gel was subsequently stained with Bio-Safe Coomassie Stain (Bio-Rad) and destained using distilled water until clear. Imaging was performed using the Gel Doc XR+ System (Bio-Rad), with careful annotation of ladder increments and sample wells.

### Cloning and purification of enzymes

Genes encoding the three pathway enzymes, each with a C-terminal 6x His-tag, were obtained as gBlocks from Integrated DNA Technologies (IDT) and individually cloned into the pET21a expression vector using the Gibson Assembly^®^ Cloning Kit (NEB) according to the manufacturer’s protocol. Primers used to generate the Gibson Assembly fragments for each gene are listed in Table S8, Supporting Information. The assembled constructs were transformed into DH5*α* cells, and positive clones were selected on LB agar plates containing ampicillin. Subsequently, the assembled plasmids were miniprepped from the DH5*α* cells using the ZymoPURE Plasmid Miniprep Kit (Zymo Research), according to the manufacturer’s instructions, and transformed into BL21(DE3) cells for protein expression and purification. Protein expression and purification of the pathway enzymes were carried out according to the PURE Proteins Preparation Protocol detailed above, with several modifications. The enzyme strains were cultured overnight in LB-Amp media from their respective glycerol stocks, followed by inoculation into 500 ml of LB-Amp media. After protein purification, each enzyme was dialyzed, concentrated, and stored in 30% glycerol buffer at –80°C for future use. The Pox5 enzyme required a modified expression protocol to address protein insolubility issues; protein expression was conducted at 15°C for 15 hours.

### Enzyme activity assays

Qualitative enzyme assays were performed to assess the functionality of each purified enzyme of the pathway before supplementing them in the PURE system. **Pyruvate oxidase (Pox5)**: The activity of pyruvate oxidase (Pox5) was measured by a spectrophotometric assay [54]. The reaction mixture (300 *µ*l) contained 15 mM pyruvate, 50 mM potassium phosphate buffer (pH 6), 0.2 mM thiamine pyrophosphate (TPP), 0.01 mM flavin adenine dinucleotide (FAD), 10 mM MgSO_4_, 0.01% (w/v) 4-aminoantipyrine, 0.02% (w/v) EHSPT (N-ethyl-N-(2-hydroxy-3-sulfopropyl)-m-toluidine), 1 mM EDTA, and 5 U/ml horseradish peroxidase. The reaction was initiated by adding purified Pox5 enzyme at a final concentration of 1 *µ*M (or 3.3 mM potassium phosphate buffer in negative control) and monitoring the reaction at 37*^◦^*C for 3 minutes. The formation of quinoneimine dye, resulting from the reaction of H_2_O_2_ (produced by Pox5 activity) with 4-aminoantipyrine and EHSPT in the presence of peroxidase, was monitored at 550 nm using BioTek Synergy H1 plate reader. **Acetate kinase (AckA)**: The activity of acetate kinase was measured using the ATP Determination Kit (Invitrogen A22066) utilizing a luciferase-coupled assay. The reaction mixture (100 *µ*l) contained 90 *µ*l of standard reaction solution prepared according to the manufacturer’s protocol, and 9 *µ*l of AckA enzyme assay mix containing 300 mM acetyl phosphate, 10 mM ADP, 3 mM magnesium acetate, and 100 mM HEPES buffer (pH 7). The reaction was initiated by adding 1 *µ*l of the purified acetate kinase enzyme at a final concentration of 1.4 *µ*M (or 1 *µ*l 100 mM HEPES buffer in negative control) and incubated at 25*^◦^*C for 3 minutes. The production of ATP from ADP and acetyl phosphate by acetate kinase was coupled to the luciferase reaction. The firefly luciferase enzyme, in the presence of D-luciferin, catalyzes a reaction that emits light proportional to the ATP concentration. The luminescence of the emitted light was measured using BioTek Synergy H1 plate reader. **Catalase (KatE)**: The activity of catalase was measured at 37*^◦^*C by monitoring the decomposition of hydrogen peroxide (H_2_O_2_) at 240 nm [55]. The reaction mixture (1 ml) contained 50 mM potassium phosphate buffer (pH 7.0) and 13.1 mM H_2_O_2_. The reaction was initiated by adding the purified catalase enzyme at a final concentration of 0.01 *µ*M and the decrease in absorbance at 240 nm (A240) was recorded immediately using a spectrophotometer for 3 mins. The decrease in A240 is directly proportional to the decomposition of H_2_O_2_. Bovine serum albumin (BSA) was used at a final concentration of 0.01 *µ*M in the negative control, and potassium phosphate buffer at pH 7.0 was used as a blank.

### Design and analysis of DOE

The circumscribed central composite design (cCCD) is a symmetrical design consisting of: a center point at the design space’s center, 8 corner points forming a box around the center, representing a 2-level full factorial design, and 6 axial points set at the minimum and maximum levels for each factor, positioned *±*1.68 times the distance from the center point to each corner point [40]. 19 conditions were set up, comprising 15 from the cCCD, and 4 from a holdout test dataset identified through Latin hypercube sampling within the bounds of the design space, to validate the model [40]. Each of the reaction conditions were compiled in triplicate, except for the center point which had six replicates, and the endpoint readings were taken at 5 hours. Linear regression was used for model fitting. Design generation, modelling and plotting was performed in Python.

## Supporting Information

Supporting Information is available online.

## Contributions

NL and SY designed the experiments. SY carried out experiments and analysed the data. AP implemented the DOE optimisation and analysed the results. SY, AP, SL, and AB developed and produced the in-house PURE system. NL supervised the project. All authors contributed to writing and editing the manuscript.

## ACKNOWLEDGMENTS

NL, SL, and AB are supported by NL’s UKRI Future Leaders Fellowship (MR/V027107/1). SY is supported by a Darwin Trust of Edinburgh PhD Studentship. AP is supported by a BBSRC Eastbio CASE PhD studentship. The authors gratefully acknowledge support from the School of Biological Sciences and the Centre for Engineering Biology at the University of Edinburgh.

