## Supporting Information for "ATP Regeneration from Pyruvate in the PURE System"

### Supporting Information: ATP regeneration from Pyruvate in the PURE system

Surendra Yadav, Alexander J. P. Perkins, Sahan B. W. Liyanagedera,
Anthony Bougas, and Nadanai Laohakunakorn
*School of Biological Sciences, University of Edinburgh*

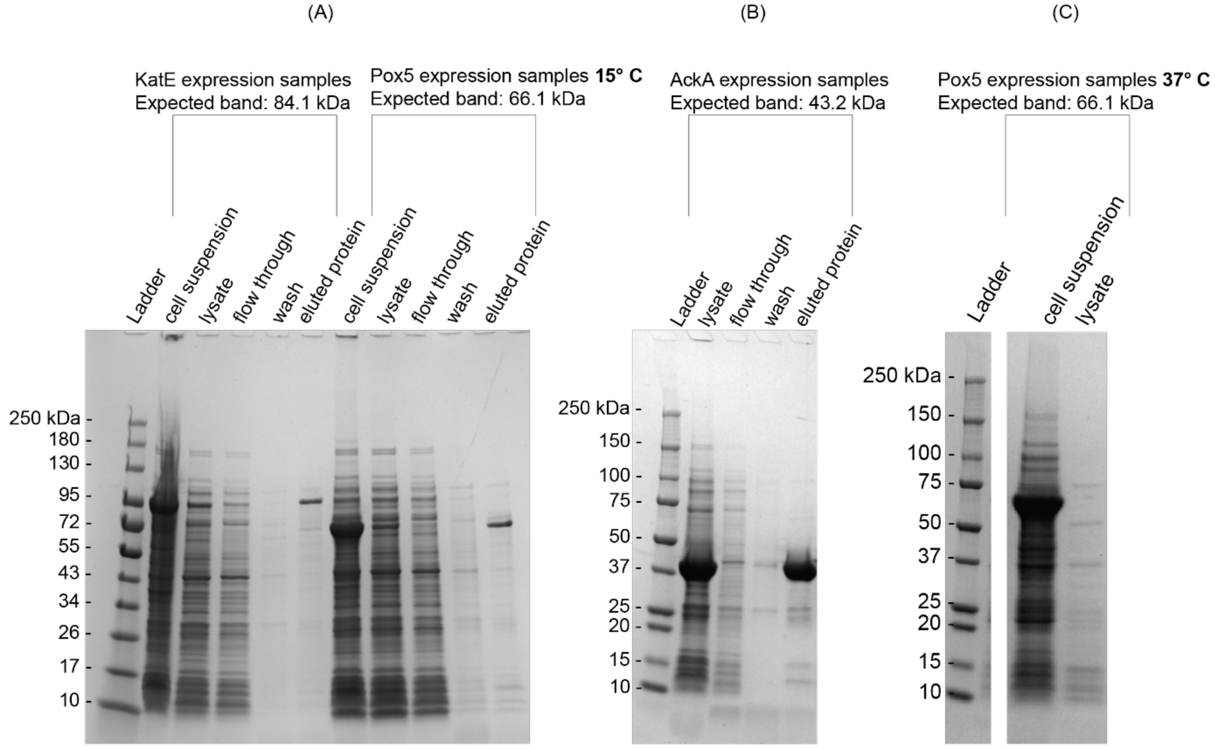

**Figure S1: SDS-PAGE imaging of Pox5, AckA and KatE protein purification samples.** Protein samples taken at different stages of protein expression and purification were run on the Mini-PROTEAN TGX Precast gels (180 V for 40 mins). KatE, Pox5 **(A)** and AckA **(B)** were overexpressed and later purified as seen in the eluted fraction. Protein expression and purification of all proteins was carried out as described in the Materials and Methods section of the article. **(C)** When Pox5 was over-expressed at 37°C, the protein was entirely sequestered in the insoluble fraction of the lysate, with no specific bands observed in the soluble fraction.

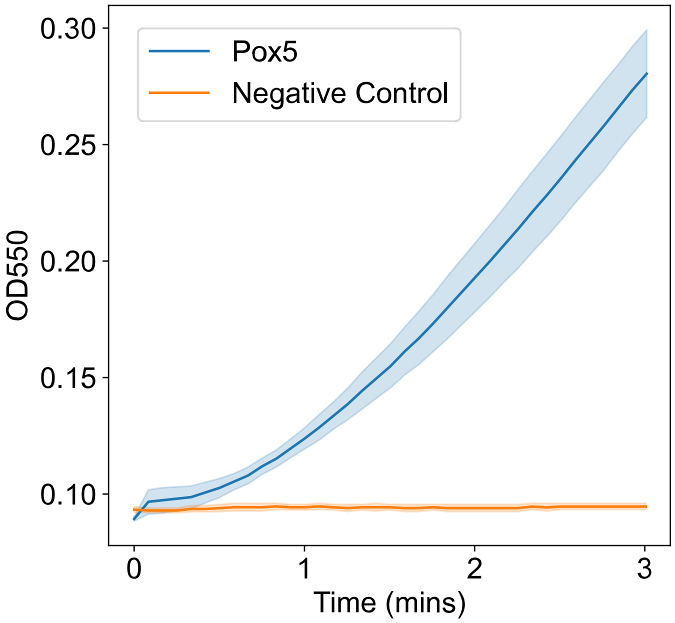

**Figure S2: Pyruvate oxidase (Pox5) enzyme activity assay.** The activity of pyruvate oxidase was measured by a spectrophotometric assay. The reaction was initiated by adding purified Pox5 enzyme at a final concentration of 1 μM (or 3.3 mM potassium phosphate buffer, pH 6 in the negative control) and monitoring the reaction at 37°C for 3 minutes. The formation of quinoneimine dye, resulting from the reaction of $H_2O_2$ with 4-aminoantipyrine and EHSPT in the presence of peroxidase, was monitored at 550 nm using a BioTek Synergy H1 plate reader. Complete details are given in the Materials and Methods section.

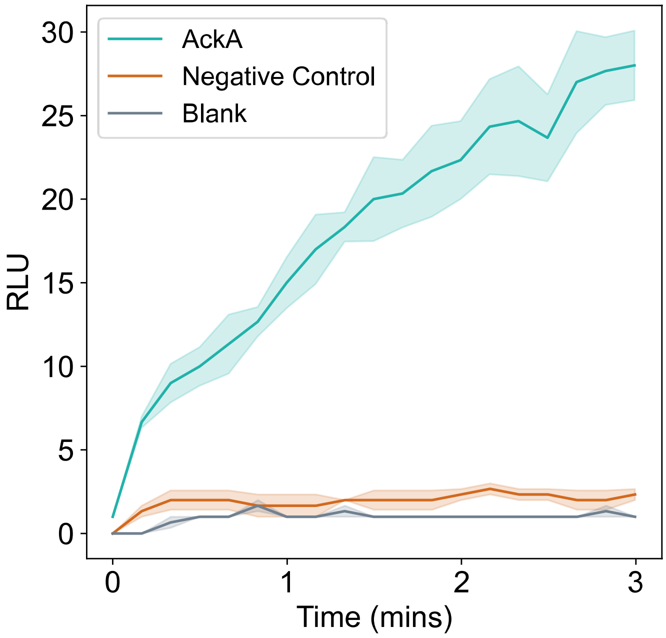

**Figure S3: Acetate kinase (AckA) enzyme activity assay.** The activity of acetate kinase was measured using the ATP Determination Kit (Invitrogen A22066) utilizing a luciferase-coupled assay. The reaction was initiated by adding 1 μL of the purified acetate kinase enzyme at a final concentration of 1.4 μM (or 1 μl 100 mM HEPES buffer in the negative control) and incubated at 25°C for 3 minutes. MilliQ® water was used as a blank. The production of ATP from ADP and acetyl phosphate by acetate kinase was coupled to the luciferase reaction. The firefly luciferase enzyme, in the presence of D-luciferin, catalyzes a reaction that emits light proportional to the ATP concentration. The luminescence of emitted light was measured using a BioTek Synergy H1 plate reader.

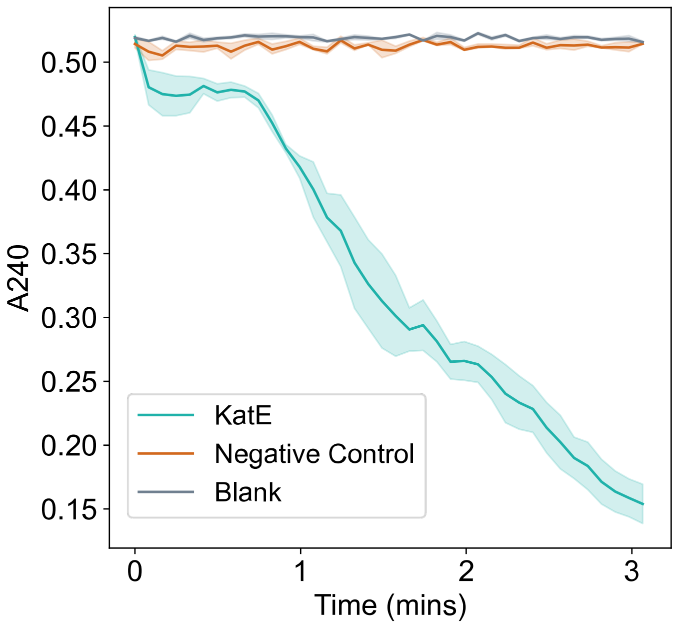

**Figure S4: Catalase (KatE) enzyme activity assay.** The activity of catalase was measured at 37°C by monitoring the decomposition of hydrogen peroxide ($H_2O_2$) at 240 nm. The reaction was initiated by adding the purified catalase enzyme at a final concentration of 0.01 µM and the decrease in absorbance at 240 nm (A240) was recorded immediately using a spectrophotometer for 3 mins. The decrease in A240 is directly proportional to the decomposition of $H_2O_2$. The negative control was 0.01 µM BSA, while the blank was potassium phosphate buffer, pH 7.

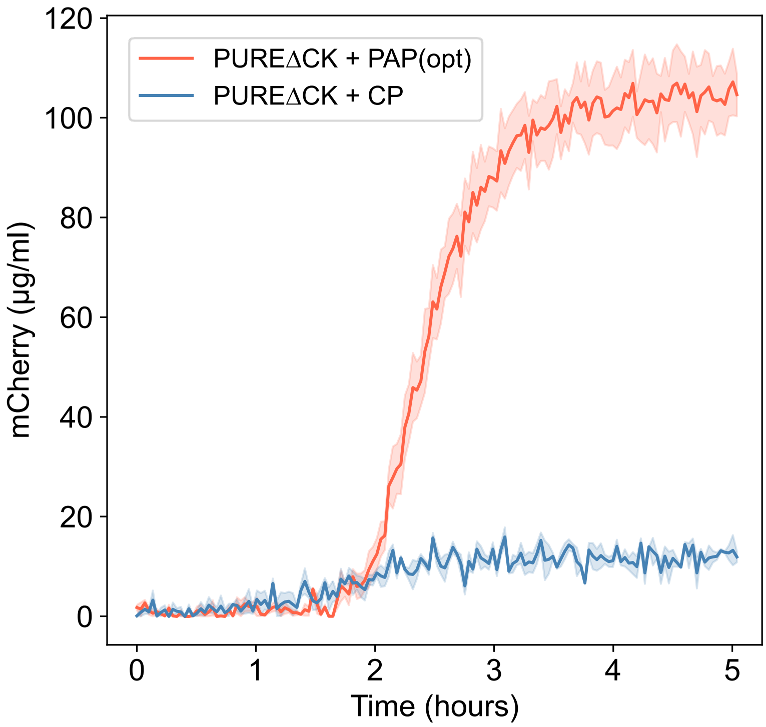

**Figure S5: PAP functions as an ATP regeneration pathway in the PUREΔCK system.** This system lacks the creatine kinase enzyme. Deletion of CK from the PURE system removes the CP/CK ATP regeneration system, resulting in low protein yield (12.0 ± 1.1 µg/mL) when supplied with creatine phosphate. However, the activity of PAP results in an increased final protein yield of 104.4 ± 4.4 µg/mL. Experiments were performed in triplicates. Data are shown as mean ± s.e. (n = 3).

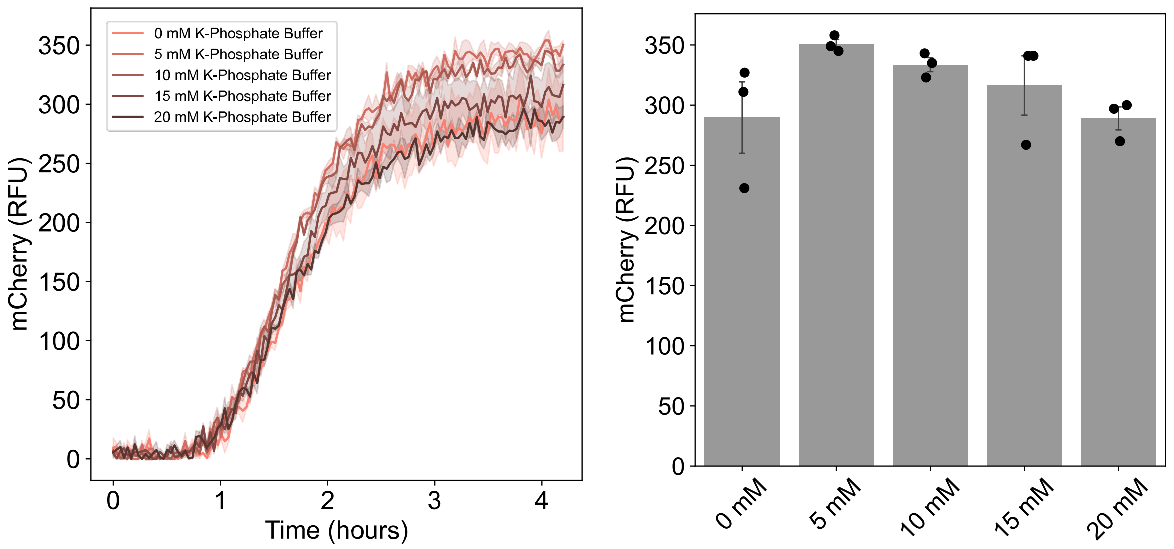

**Figure S6: Potassium phosphate buffer (pH 7) titration in the PURE system.** Phosphate titration was performed in the PURE reactions utilizing the CP/CK energy regeneration component by adding exogenous potassium phosphate buffer (pH 7) at the given final concentrations at the start of the reactions. The bar plot shows the final mCherry levels at 4h. Data are shown as mean ± s.e. (n = 3).

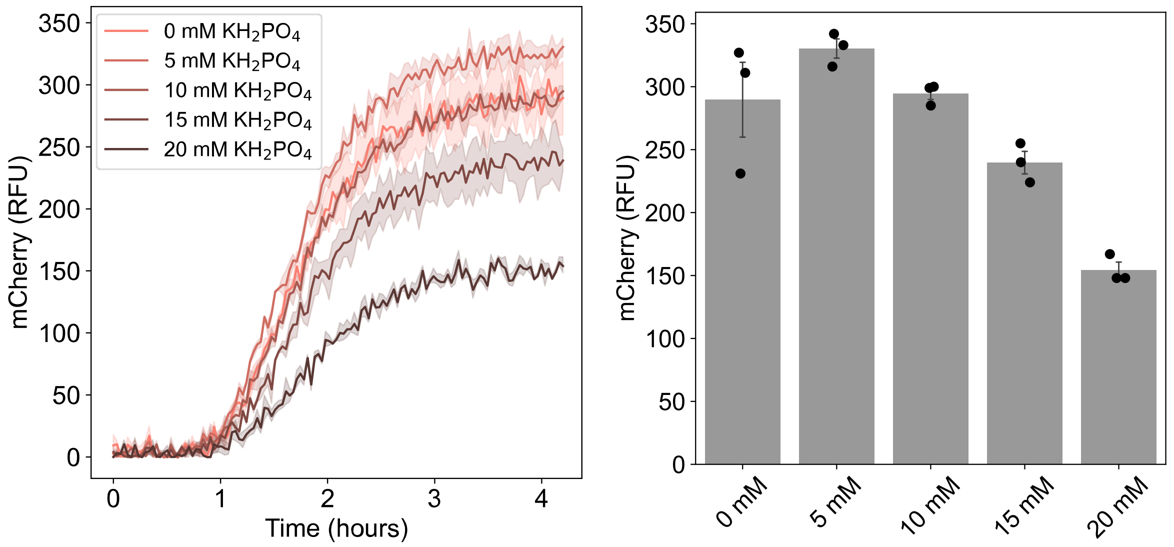

**Figure S7: Potassium phosphate monobasic (pH 4.5) titration in the PURE system.** Phosphate titration was performed in the PURE reactions utilizing the CP/CK energy regeneration component by adding exogenous $KH_2PO_4$, pH 4.5, at the given final concentrations at the start of the reactions. The bar plot shows the final mCherry levels at 4h. Data are shown as mean ± s.e. (n = 3).

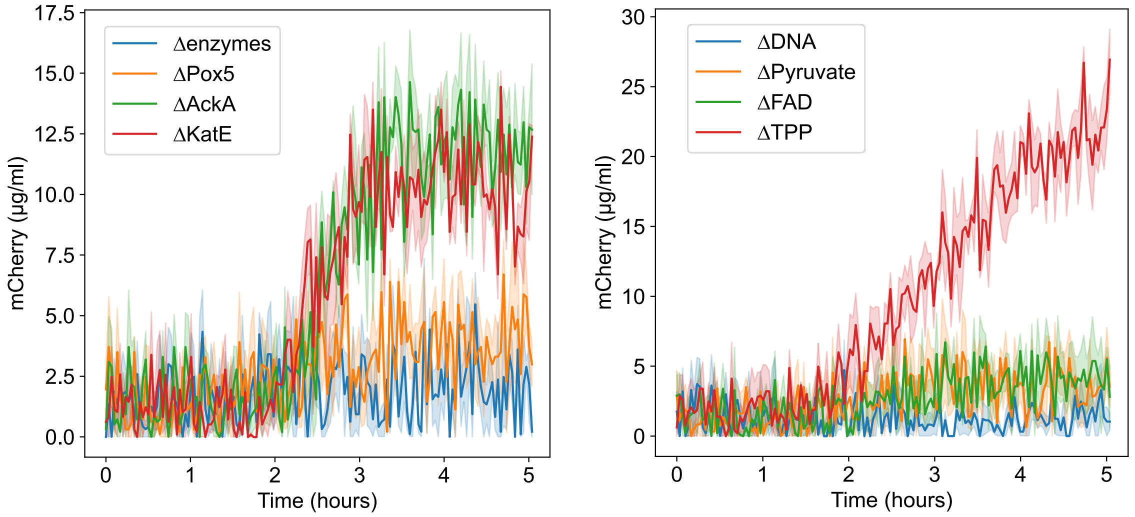

**Figure S8: Timeseries data of negative controls of the PAP**. Timeseries data of mCherry protein expression in the unoptimized pathway with excluded components (Fig 2b). The reactions were carried out with 10 mM K-phosphate buffer in the reaction. Data are shown as mean ± s.e. (n = 3).

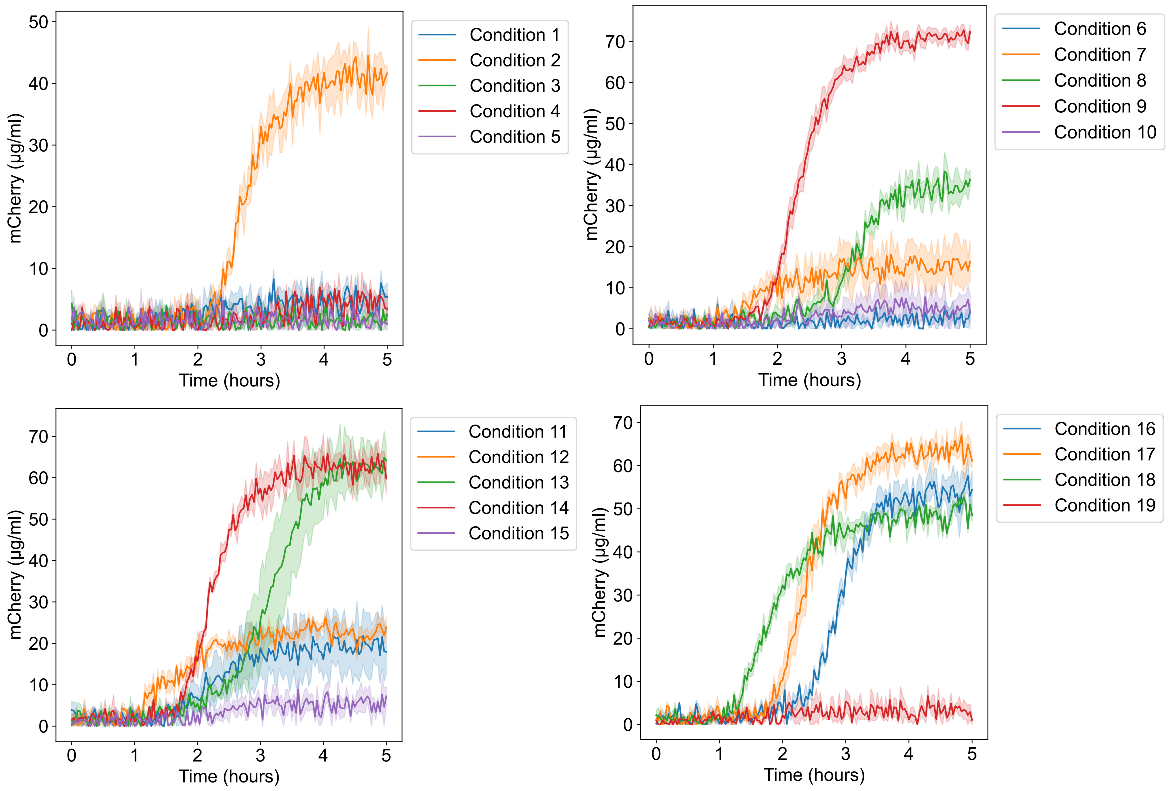

**Figure S9: Timeseries plots for all tested conditions in the DOE dataset.** Timeseries data of mCherry protein expression at different conditions of the DOE design. Data are shown as mean ± s.e. (n = 3 except Condition 9, where n=6).

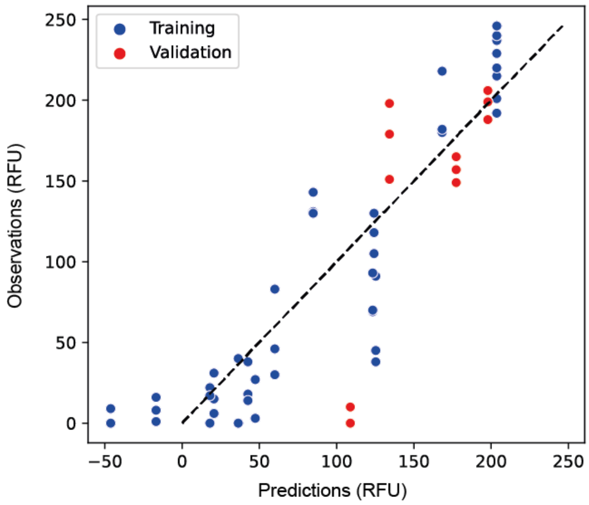

**Figure S10: Fitted model evaluation.** Plot of the regression model predictions against experimental measurements, for training data used to calibrate the model (blue points) and held-out data used for validation (red points).

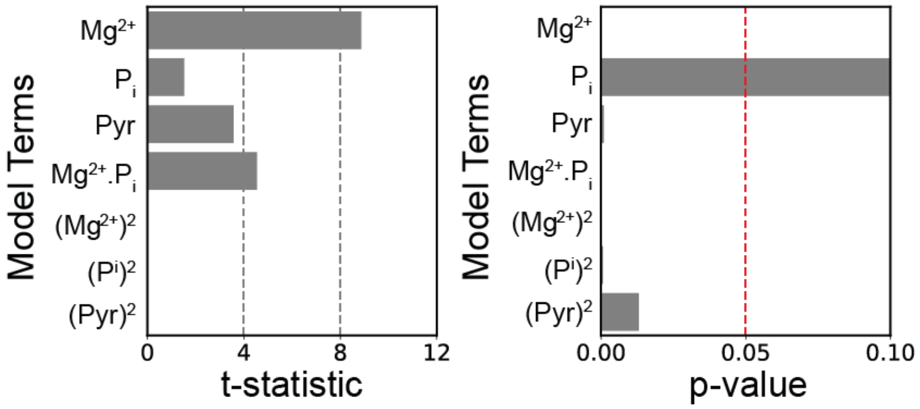

**Figure S11: Model terms analysis.** T-statistics and corresponding p-values for each of the model terms, showing that the $Mg^{2+}$ concentration is the most sensitive factor, followed by pyruvate, phosphate, and the magnesium-phosphate interaction.

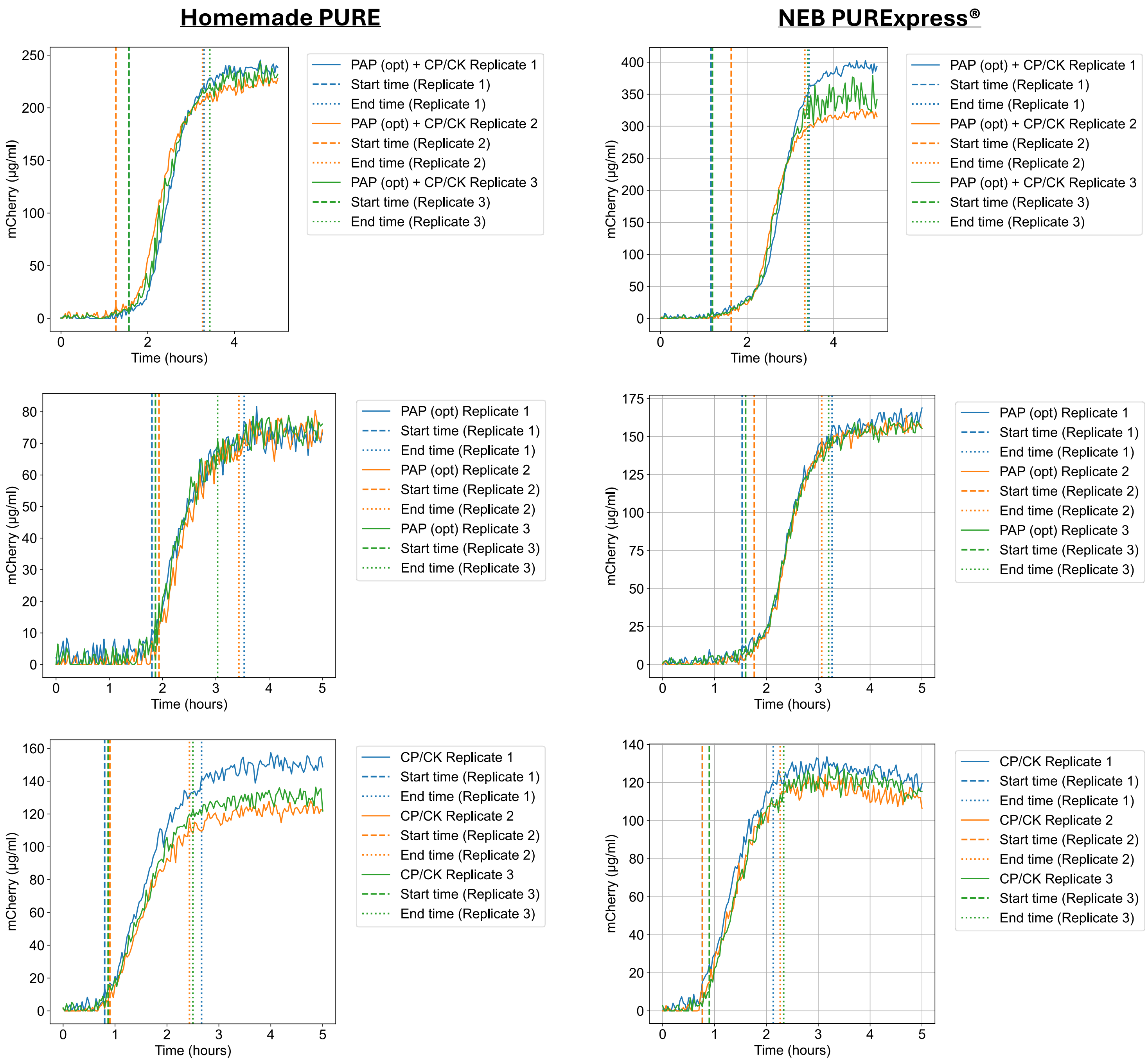

**Figure S12: Reaction lifetime and lag time analysis.** Specific start and end time points were used to calculate both the lag and reaction lifetime for different reactions in homemade and PURExpress systems. The start time was defined as the moment when the mCherry protein yield reached a concentration of 10 µg/ml, while the end time was defined as the point at which the protein yield reached 90% of its maximum value. The lag time is equal to the start time as defined here, while the reaction lifetime is calculated as the difference between the start and end times.

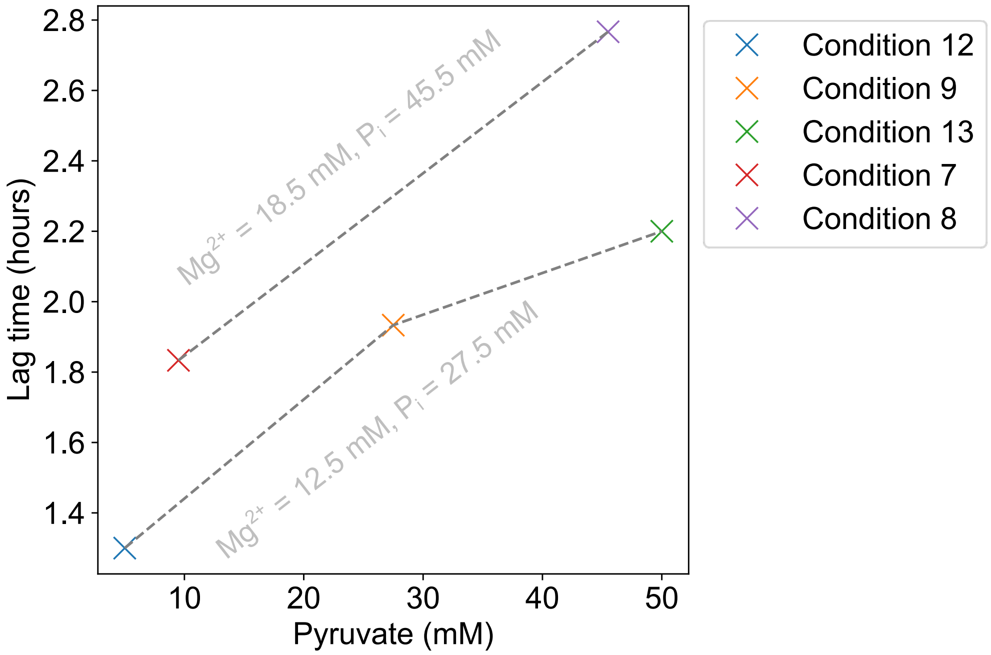

**Figure S13: Relation of initial pyruvate concentration to reaction lag time from the DOE dataset.** The lag time increases with the increasing initial concentration of pyruvate in PAP-powered reactions. Analysis of the DOE data indicated a correlation between the lag time and pyruvate concentration. Conditions 9, 12, and 13 had identical reaction compositions, differing only in pyruvate concentration. Similarly, Conditions 7 and 8 were identical except for pyruvate concentration.

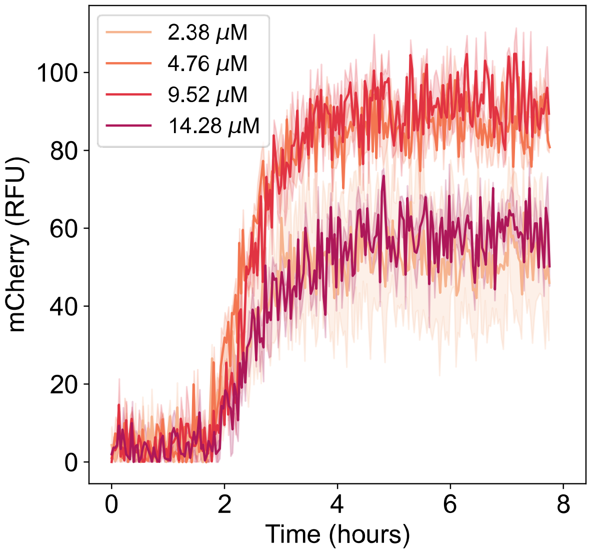

**Figure S14: Catalase (KatE) titration in the PAP powered PURE system.** Catalase titration was performed in the PURE system to investigate whether increasing catalase concentration influences the initial lag time in protein synthesis. No correlation was observed.

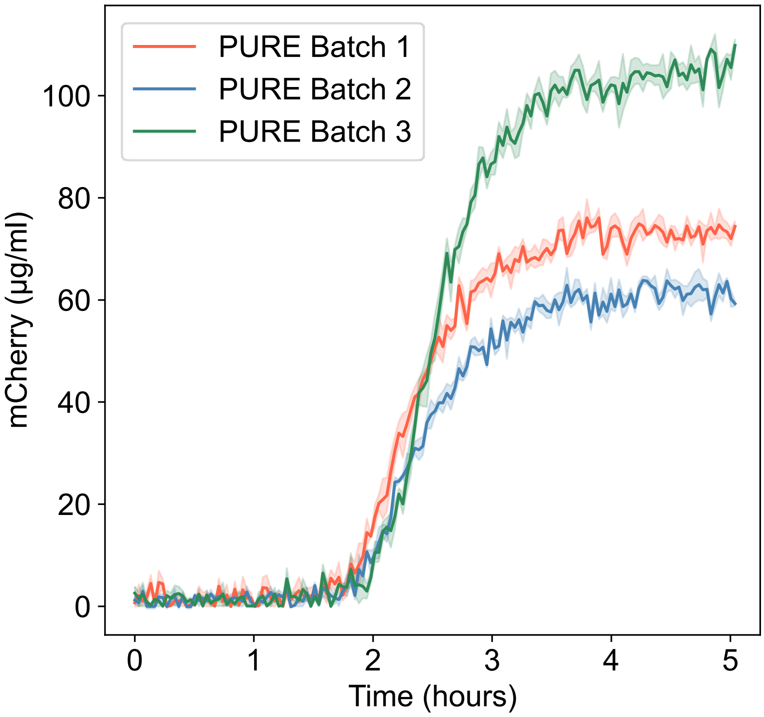

**Figure S15: PAP is active across different batches of PURE.** PAP functions as an ATP regeneration component across different batches of PURE produced in-house. PURE Batch 1 and Batch 2 were prepared alongside each other, and Batch 3 was prepared in a separate session. Experiments were performed in triplicates. Data are shown as mean ± s.e. (n = 3).

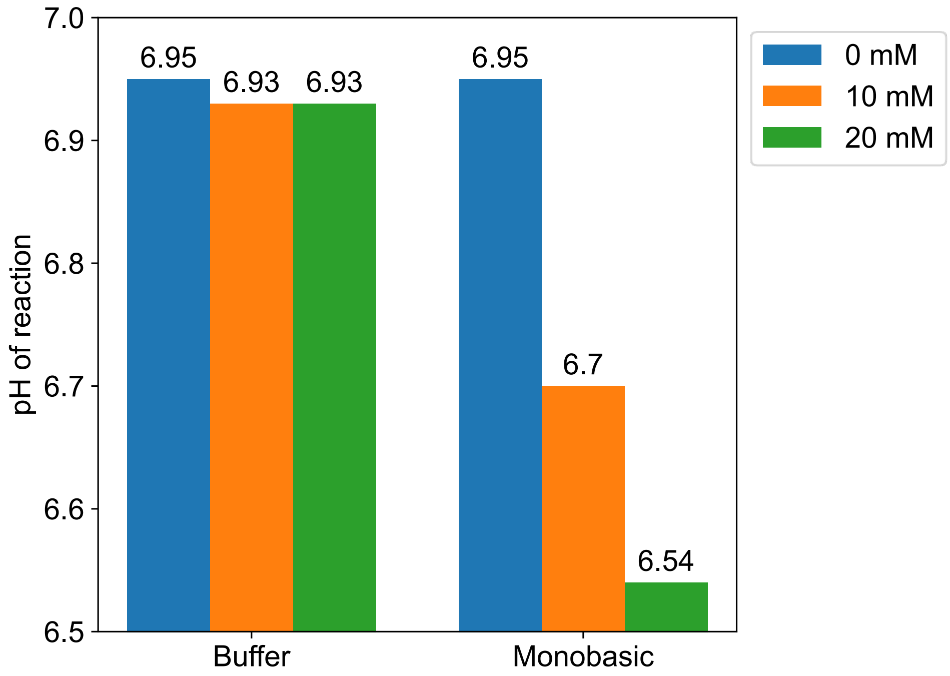

**Figure S16: Initial pH of reactions supplemented with phosphates.** Initial pH of reactions supplemented with monobasic potassium phosphate or potassium phosphate buffer (pH 7). While increasing the concentration of phosphate buffer (up to 20 mM) had negligible effects on the reaction pH, higher concentrations of monobasic phosphate significantly lowered the pH from 6.95 to 6.54. Measurements were taken with Mettler Toledo FiveEasy Plus pH meter using a Mettler Inlab Micro Electrode.

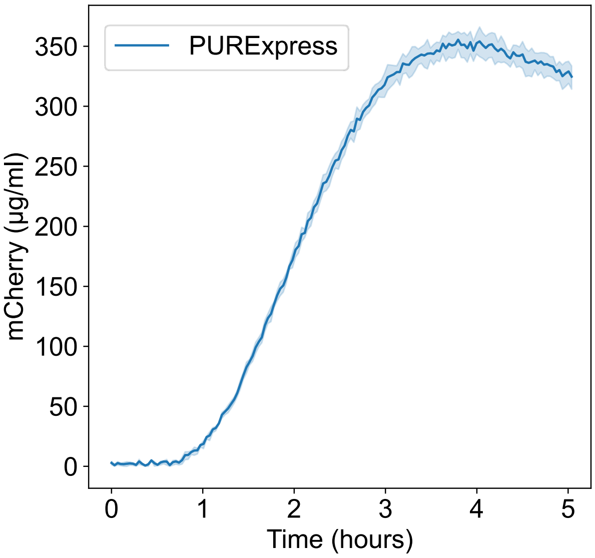

**Figure S17: mCherry expression in PURExpress.** Timeseries data of mCherry protein expression using the PURExpress kit. The maximal synthesis rates of the mCherry protein in PURExpress reactions were measured to be 4.3 ± 0.2 (µg/mL)/min. Data are shown as mean ± s.e. (n = 3).

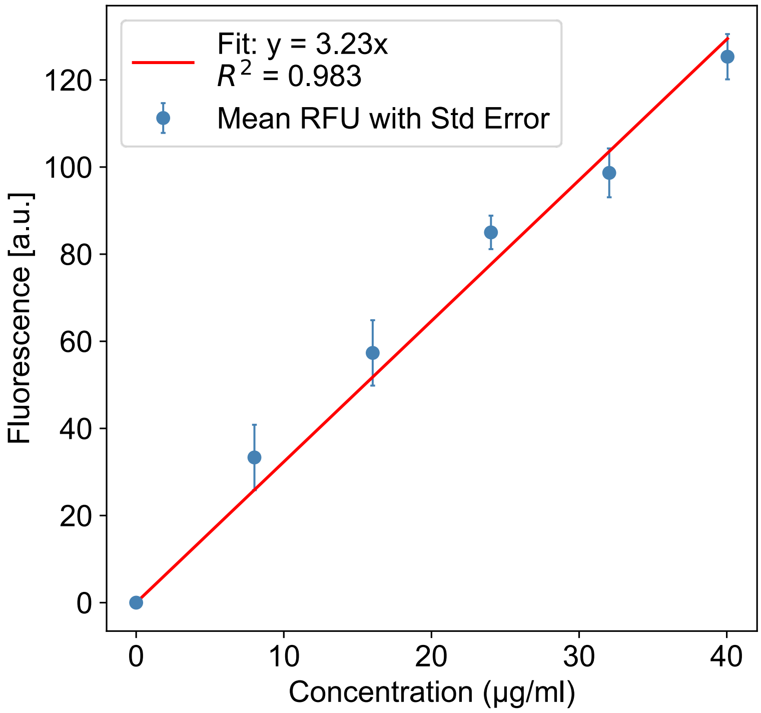

**Figure S18: Standard calibration curve for mCherry.** The standard curve was produced by measuring the fluorescence of purified mCherry at different concentrations in PBS on a plate reader with the same settings as for CFPS reactions. Excitation and emission wavelengths were 579 nm and 616 nm, respectively. Experiments were performed in triplicates. Data are shown as mean ± s.e. (n = 3).

#### Experimental Details

**PURE Production Protocol:** OnePot PURE was prepared as previously described with some modifications [Lavickova et al., ACS Synthetic Biology 2019]. Working glycerol stock plates were prepared for all 36 strains required for the PURE system at an optical density (600 nm) of 0.5 and stored at -80°C till use. To prepare starter cultures, 10 µL of each working glycerol stock (except EF-Tu) was used to inoculate 300 µL of LB media containing ampicillin in a 1 mL 96 deep well plate and sealed with breathable plate seal. For EF-Tu starter, 10 µL EF-Tu working glycerol stock was used to inoculate 3 mL of LB media containing ampicillin in a 15 mL falcon tube. The starter plate was incubated at 37°C in a plate shaker at 1000 RPM for 14 hours, whilst the EF-Tu starter was incubated at 37°C in a shaking incubator at 220 RPM.

The next day 500 mL of pre-warmed LB media containing ampicillin was inoculated with 1675 µL of EF-Tu starter and 55 µL of all other starters. The culture was incubated at 37°C in a shaking incubator at 220 RPM until an optical density (600 nm) of 0.2 - 0.3 at which point protein expression was induced by the addition of IPTG to a final concentration of 0.1 mM. Cultures were grown for a further 3 hours prior to cell harvest by centrifugation at 5000g for 15 minutes at 4°C. Cell pellets from 500 mL initial culture were re-suspended in 20 mL of ice-cold PBS to remove residual media and transferred to a 50 mL flacon tube and centrifuged at 3000g for 8 minutes at 4°C and PBS removed by decanting to obtain washed cell pellet. Washed cell pellets were centrifuged again at 3000g for 2 minutes at 4°C and residual PBS was removed via pipetting. Cell pellet mass recorded before flash freezing in liquid nitrogen and stored at -80°C.

The next day the cell pellet was thawed on ice and re-suspended in 7.5 mL Buffer A containing 1 mM TCEP by vortexing. Cells were lysed by sonication (Fisherbrand Model 120, #12337338, Fisher Scientific) at 70% amplitude, 10s on 10s off until 2000 J had been transferred in an ice water bath using a cooled sonication probe (Fisherbrand #12931181, Fisher Scientific). Following sonication, the lysate was briefly vortexed and subsequently split into 2 mL micro-centrifuge tubes in 1 mL aliquots. Lysate was clarified by centrifugation at 15923g for 20 minutes at 4°C. Pellet free supernatant was collected into a 50 mL tube on ice, combined with 2 mL of cOmplete Ni-NTA Resin (Roche) equilibrated with Buffer A and incubated with end over end mixing at 4°C for 3 hours. Subsequently, the resin and lysate mixture was added to a chromatography column and washed with 25 mL of Wash Buffer (24.75 mL Buffer A + 0.25 mL Buffer B) containing 1 mM TCEP. Then 5 mL of elution buffer (0.5 mL Buffer A + 4.5 mL Buffer B) containing 1 mM TCEP was added to the column and allowed to incubate with resin for 10 minutes following which the eluant was collected into a 5 mL tube on ice.

Eluted protein solution was then dialysed against 1L of Buffer HT (w/o TCEP) overnight in a 2 kDa MWCO dialysis cassette. The dialysed sample was combined with 10 mL of fresh Buffer HT containing 1 mM TCEP and concentrated down to 1.5 mL using a 3 kDA MWCO Amicon Ultra 15 centrifugal filter. The sample was then transferred to a 3kDa MWCO Amicon Ultra 0.5 centrifugal filter and concentrated to 25 mg/mL. Following concentration, the protein solution was transferred to a microcentrifuge tube and centrifuged at 20,000g for 10 minutes at 4°C to pellet precipitated proteins. The pellet free supernatant was extracted, volume quantified and an equal volume of Stock 60 with 1mM TCEP added to bring the final concentration of the PURE protein solution to 12.5 mg/mL. This stock was aliquoted, and flash frozen in liquid nitrogen and stored at -80°C till reaction setup.

**Crude Ribosome Purification:** A 5 mL 2x YTP mini culture was inoculated with a glycerol stock scraping of BL21 and incubated at 37°C at 220 RPM overnight. Additionally, a 5 mL 2x YTP media sample without BL21 inoculation was also incubated under the same conditions to check for media contamination. The remaining media was left to prewarm overnight.

The next day, the incubated media and prewarmed media bottles were checked for contamination before proceeding. A 200 mL 2xYTP media midi culture in a 500 mL flask was inoculated with the mini culture to achieve an initial optical density (600 nm) of 0.05. This culture was grown at 37°C at 220 RPM until it reached an optical density (600 nm) of 1. Subsequently, 750 mL 2xYTP media maxi cultures (x4) in 2.5 L baffled flasks were inoculated with the midi culture to achieve an initial optical density (600 nm) of 0.05. The maxi cultures were grown until they reached an optical density (600 nm) of 0.6. The culture density was measured every 15 minutes after the first hour to ensure optimal harvest time. During growth, centrifuge bottles were placed on ice to cool, and all subsequent steps were conducted on ice unless otherwise stated. Upon reaching the desired optical density, the flasks were placed in an ice water bath for 20 minutes to halt growth. Cells were harvested by centrifugation at 4 °C and 3220 x g for 10 minutes, with no more than 500 mL of culture per centrifuge bottle. The media was then decanted, bottles were blotted dry, and cell pellets were placed back on ice. Next 100 mL of sterile ice-cold PBS was added to each centrifuge bottle and cells thoroughly resuspended by vortexing. Cells were again harvested by centrifugation at 4 °C and 3220 x g for 10 minutes. After decanting the PBS and blotting the bottles dry, the cell pellets from the total 3L culture were again resuspended in 200 mL of PBS. Resuspended cells were split between four pre-weighed 50 mL falcon tubes, and pelleted again 3000g for 10 minutes at 4°C. The PBS was removed by decanting and pipetting, and the cell pellet mass was recorded before snap freezing with liquid nitrogen.

The next day, whilst cell pellets were thawed on ice, 200 µL of 0.5 M DTT was added to 100 mL of Ribo Buffer A to achieve a final concentration of 1 mM DTT. The cells were resuspended in Ribo Buffer A (+DTT) at a ratio of 2 mL per 1 g of cell pellet by thorough vortexing. The resuspension was divided such that each falcon tube contained 5 mL of the cell suspension, which was then lysed via sonication (#12337338, Fisher Scientific) at 30% amplitude, 20s on 20s off for 10 cycles in an ice water bath using a cooled sonication probe (#12931181, Fisher Scientific).

The lysates from the initial 3L culture were split between two pre-chilled S30 centrifugation tubes, and each tube topped up to 25 mL with Ribo Buffer A (+DTT) before centrifugation at 30,000g for 1 hour at 4 °C. The pellet free supernatant was split between two pre-chilled S100 tubes and each tube topped up to 25 mL with Ribo Buffer A (+DTT) before ultracentrifugation at 100,000g for 4 hours at 4 °C. Following ultracentrifugation, the supernatant was carefully decanted and the ribosome pellets in each tube were soaked overnight in 5 mL of Ribo Buffer B containing 1mM DTT (in each tube).

The next day a further 5 mL of Ribo Buffer B containing 1 mM DTT was added to each tube and the ribosome pellets were resuspended carefully using a 1mL pipette. The resuspended ribosome solution was split between two pre-chilled S30 centrifugation tubes, and each tube topped up to 25 mL with Ribo Buffer B (+DTT) before centrifugation at 30,000g for 1 hour at 4 °C. The pellet free supernatant was split between two pre-chilled S100 tubes and each tube topped up to 25 mL with Ribo Buffer B (+DTT) before ultracentrifugation at 100,000g for 4 hours at 4 °C. Following ultracentrifugation, the supernatant was carefully decanted and the ribosome pellets in each tube were soaked overnight in 5 mL of Ribo Buffer B containing 1mM DTT (in each tube) as previously described.

The next day, the ribosome pellets were again resuspended in Ribo Buffer B containing 1 mM DTT as previously described. The resuspended ribosome solutions were split between two new pre-chilled S100 tubes and each tube topped up to 25 mL with Ribo Buffer B (+DTT) before ultracentrifugation at 100,000g for 4 hours at 4°C. Following ultracentrifugation, the supernatant was carefully decanted and the ribosome pellets in each tube were soaked overnight in 400 µL of Ribo Buffer A containing 1mM DTT (in each tube) as previously described.

Finally, the next day, the ribosome pellet was resuspended by gentle pipetting without addition of more buffer. A 100x dilution of the solution was made and concentration tested by nanodrop at 260 nm, whereby an optical density of 10 of the diluted sample at 260 nm equates to 24 µM ribosomes in the undiluted stock [Rheinberger et al., Methods in Enzymology 1988]. Ribosome solutions were aliquoted, and flash frozen in liquid nitrogen and stored at -80°C till reaction setup.

#### Table S1: Design of experiments (DOE) dataset

| Pyruvate (mM) | Phosphate (mM) | Mg2+ (mM) | Data Point Type | Condition ID | Protein Yield (RFU) |
|---|---|---|---|---|---|
| 9.5 | 9.5 | 6.5 | Corner | 1 | 31 |
| 9.5 | 9.5 | 6.5 | Corner | 1 | 15 |
| 9.5 | 9.5 | 6.5 | Corner | 1 | 6 |
| 45.5 | 9.5 | 6.5 | Corner | 2 | 131 |
| 45.5 | 9.5 | 6.5 | Corner | 2 | 143 |
| 45.5 | 9.5 | 6.5 | Corner | 2 | 130 |
| 9.5 | 45.5 | 6.5 | Corner | 3 | 1 |
| 9.5 | 45.5 | 6.5 | Corner | 3 | 8 |
| 9.5 | 45.5 | 6.5 | Corner | 3 | 16 |
| 45.5 | 45.5 | 6.5 | Corner | 4 | 27 |
| 45.5 | 45.5 | 6.5 | Corner | 4 | 3 |
| 45.5 | 45.5 | 6.5 | Corner | 4 | 3 |
| 9.5 | 9.5 | 18.5 | Corner | 5 | 9 |
| 9.5 | 9.5 | 18.5 | Corner | 5 | 0 |
| 9.5 | 9.5 | 18.5 | Corner | 5 | 0 |
| 45.5 | 9.5 | 18.5 | Corner | 6 | 22 |
| 45.5 | 9.5 | 18.5 | Corner | 6 | 0 |
| 45.5 | 9.5 | 18.5 | Corner | 6 | 17 |
| 9.5 | 45.5 | 18.5 | Corner | 7 | 30 |
| 9.5 | 45.5 | 18.5 | Corner | 7 | 46 |
| 9.5 | 45.5 | 18.5 | Corner | 7 | 83 |
| 45.5 | 45.5 | 18.5 | Corner | 8 | 105 |
| 45.5 | 45.5 | 18.5 | Corner | 8 | 130 |
| 45.5 | 45.5 | 18.5 | Corner | 8 | 118 |
| 27.5 | 27.5 | 12.5 | Center | 9 | 215 |
| 27.5 | 27.5 | 12.5 | Center | 9 | 220 |
| 27.5 | 27.5 | 12.5 | Center | 9 | 246 |
| 27.5 | 27.5 | 12.5 | Center | 9 | 237 |
| 27.5 | 27.5 | 12.5 | Center | 9 | 240 |
| 27.5 | 27.5 | 12.5 | Center | 9 | 246 |
| 27.5 | 27.5 | 5 | Axial | 10 | 0 |
| 27.5 | 27.5 | 5 | Axial | 10 | 0 |
| 27.5 | 27.5 | 5 | Axial | 10 | 40 |
| 27.5 | 5 | 12.5 | Axial | 11 | 38 |
| 27.5 | 5 | 12.5 | Axial | 11 | 45 |
| 27.5 | 5 | 12.5 | Axial | 11 | 91 |
| 5 | 27.5 | 12.5 | Axial | 12 | 69 |
| 5 | 27.5 | 12.5 | Axial | 12 | 70 |
| 5 | 27.5 | 12.5 | Axial | 12 | 93 |
| 50 | 27.5 | 12.5 | Axial | 13 | 192 |
| 50 | 27.5 | 12.5 | Axial | 13 | 201 |
| 50 | 27.5 | 12.5 | Axial | 13 | 229 |
| 27.5 | 50 | 12.5 | Axial | 14 | 218 |
| 27.5 | 50 | 12.5 | Axial | 14 | 180 |

| | | | | | |
|---|---|---|---|---|---|
| 27.5 | 50 | 12.5 | Axial | 14 | 182 |
| 27.5 | 27.5 | 20 | Axial | 15 | 18 |
| 27.5 | 27.5 | 20 | Axial | 15 | 14 |
| 27.5 | 27.5 | 20 | Axial | 15 | 38 |
| 39.38 | 44.96 | 8.84 | Validation | 16 | 151 |
| 39.38 | 44.96 | 8.84 | Validation | 16 | 179 |
| 39.38 | 44.96 | 8.84 | Validation | 16 | 198 |
| 28.58 | 22.82 | 11.78 | Validation | 17 | 206 |
| 28.58 | 22.82 | 11.78 | Validation | 17 | 188 |
| 28.58 | 22.82 | 11.78 | Validation | 17 | 199 |
| 16.88 | 34.88 | 13.04 | Validation | 18 | 149 |
| 16.88 | 34.88 | 13.04 | Validation | 18 | 157 |
| 16.88 | 34.88 | 13.04 | Validation | 18 | 165 |
| 19.22 | 15.98 | 16.04 | Validation | 19 | 0 |
| 19.22 | 15.98 | 16.04 | Validation | 19 | 0 |
| 19.22 | 15.98 | 16.04 | Validation | 19 | 10 |

**Table S2: Fitted model parameters**

| Coefficient | Fitted value | Units |
|---|---|---|
| $\beta_0$ | $-3.63 \times 10^{2}$ | RFU |
| $\beta_1$ | $6.42 \times 10^{1}$ | RFU/mM |
| $\beta_2$ | $2.97 \times 10^{0}$ | RFU/mM |
| $\beta_3$ | $6.15 \times 10^{0}$ | RFU/mM |
| $\beta_4$ | $3.33 \times 10^{-1}$ | RFU/(mM)$^2$ |
| $\beta_5$ | $-2.92 \times 10^{0}$ | RFU/(mM)$^2$ |
| $\beta_6$ | $-1.12 \times 10^{-1}$ | RFU/(mM)$^2$ |
| $\beta_7$ | $-7.93 \times 10^{-2}$ | RFU/(mM)$^2$ |

#### Table S3: PURE reaction composition

| Component | Stock concentrations | Concentration of components in CP/CK based reactions | Concentration of components in PAP (opt) based reactions | Concentration of components in PAP (opt) + CP/CK based reactions | Units |
|---|---|---|---|---|---|
| 4x Energy Solution | 4 | 1 | 1 | 1 | X |
| PURE Protein Solution | 12.5 | 2.4 | 2.4 | 2.4 | mg/mL |
| Ribosome Solution | 24 | 2.275 | 2.275 | 2.275 | µM |
| mCherry DNA template | 337 | 10 | 10 | 10 | nM |
| Magnesium glutamate | 500 | 11.8 | 12.5* | 12.5 | mM |
| PEG 8000 | 40 | 2 | 2 | 2 | % |
| Potassium glutamate | 2000 | 100 | 100 | 100 | mM |
| Creatine phosphate | 1000 | 20 | 0 | 20 | mM |
| Potassium phosphate buffer (pH7) | 1000 | 0* | 27.5* | 27.5 | mM |
| Potassium phosphate monobasic | 1000 | 0* | 0* | 0 | mM |
| Pyruvate | 2000 | 0 | 27.5* | 27.5 | mM |
| Pox5 | 5.2 | 0 | 3.03* | 3.03 | µM |
| AckA | 24.4 | 0 | 13.89 | 13.89 | µM |
| KatE | 7.2 | 0 | 2.38* | 2.38 | µM |
| TPP | 100 | 0 | 2 | 2 | mM |
| FAD | 10 | 0 | 0.2 | 0.2 | mM |

*Concentration of the component varied according to the specific experiment's requirements, and the details of those concentrations are mentioned in the results section of the article.

PURExpress reaction compositions were the same as mentioned above except for the following modifications: 6 µl (in 50µl Master Mix) Factor Mix from PURExpress ΔRibosome Kit was used as PURE protein solution. The manufacturer's reaction setup protocol was used for PURExpress Control reaction (Fig S17) with 10 nM mCherry DNA template.

##### Table S4: Materials

| Name | Company | Catalog Number |
|---|---|---|
| 10x Tris/Glycine/SDS buffer | Bio-Rad Laboratories | 1610732 |
| 384-well µClear black plates | Greiner | 781906 |
| 4-20% Mini-PROTEAN TGX Precast Protein Gels | Bio-Rad Laboratories | 4561096 |
| 96-Well Polypropylene DeepWell plate | Nunc | 260251 |
| AMICON ULTRA 0.5 mL - 3 KDa | Merck Millipore | UFC500324 |
| AMICON ULTRA 15 mL - 3 KDa | Merck Millipore | UFC900324 |
| Amino acids | Biotech Rabbit | BR1401801 |
| Ammonium chloride | Sigma-Aldrich | 09718-1KG |
| Ammonium acetate | Sigma-Aldrich | 09689 |
| Ampicillin | Sigma-Aldrich | A8351 |
| Color Prestained Protein Standard | NEB | P7719S |
| Precision Plus Protein™ All Blue Prestained Protein Standards | Bio-Rad Laboratories | 1610373 |
| Breathe-Easy sealing membrane | Sigma-Aldrich | Z380059-1PAK |
| Ultracentrifuge tubes | Beckman Coulter | 355618 |
| Creatine phosphate | Sigma-Aldrich | 27920 |
| DNA Clean & Concentrator-25 | Zymo | ZYM-D4034-200TS |
| DTT | Roche | DTT-RO |
| Econo-Pac Chromatography Columns | Bio-Rad Laboratories | 7321010 |
| EDTA (Ethylenediaminetetraacetic acid) | Sigma-Aldrich | 03609-250G |
| Tunair™ shake flask | Sigma-Aldrich | Z710822 |
| 15 mL tubes | Sarstedt | 62.554.502 |
| 50 mL tubes | Sarstedt | 62.547.254 |
| 1.5 mL tubes | Eppendorf | 0030120086 |
| 2 mL tubes | Eppendorf | 0030120094 |
| Folinic acid | Sigma-Aldrich | PHR1541 |
| Glycerol | Sigma-Aldrich | G5516-1L |
| HEPES | Sigma-Aldrich | H3375 |
| Imidazole | Sigma-Aldrich | I2399 |
| Bio-Safe™ Coomassie Stain | Bio-Rad Laboratories | 1610786 |
| IPTG (Isopropyl-beta-D-thiogalactoside) | Thermo Scientific™ | R0392 |
| 4x Laemmli Sample Buffer | Bio-Rad Laboratories | 1610747 |
| Magnesium chloride | Sigma-Aldrich | M2670 |
| Magnesium glutamate | Sigma-Aldrich | 49605 |
| Magnesium sulphate | Sigma-Aldrich | M7506 |
| NTP | Thermo Scientific™ | R0481 |
| Phusion High-Fidelity PCR Master Mix with HF Buffer | Thermo Scientific™ | F531L |
| Potassium chloride | Sigma-Aldrich | P5405 |

| | | |
|---|---|---|
| Potassium glutamate | Sigma-Aldrich | 49601 |
| Potassium phosphate dibasic | Sigma-Aldrich | 795496 |
| Potassium phosphate monobasic | Sigma-Aldrich | P5379 |
| PURExpress Δ Ribosome Kit | NEB | E3313S |
| Quick Start Bradford 1x Dye Reagent | Bio-Rad Laboratories | 5000205 |
| Rapid-Flow Sterile Single Use Vacuum Filter Units | Thermo Scientific™ | 596-3320 |
| SealPlate film | Excel Scientific | Z369659-100EA |
| Spermidine | Sigma-Aldrich | S2626 |
| TCEP (Tris(2-carboxyethyl) phosphine-hydrochloride) | Sigma-Aldrich | 646547 |
| Tris base | Sigma-Aldrich | 93350 |
| tRNA | Roche | 10109541001 |
| PEG 8000 | Sigma-Aldrich | 89510 |
| Pyruvate | Sigma-Aldrich | P2256 |
| TPP | Sigma-Aldrich | C8754 |
| FAD | Sigma-Aldrich | F6625 |
| Acetyl phosphate | MedChem Express | HY-128730_100MG |
| Horseradish peroxidase | Thermo Scientific™ | 31490 |
| 4-aminoantipyrine | Sigma-Aldrich | A4382 |
| EHSPT (N-ethyl-N-(2-hydroxy-3-sulfopropyl)-m-toluidine) | Sigma-Aldrich | 04340 |
| ADP | Sigma-Aldrich | A2754 |
| Hydrogen peroxide | Sigma-Aldrich | H1009 |
| Bovine serum albumin (BSA) | Bio-Rad Laboratories | 5000206 |

**Table S5: List of *E. coli* strains (excluding PURE strains)**

| Strain | Application |
|---|---|
| DH5α | Cloning and plasmid maintenance of the following constructs:<br><br>a. pET21a-pox5-6xHis<br>b. pET21a-katE-6xHis<br>c. pET21a-ackA-6xHis |
| Top10 | 1. Cloning and plasmid maintenance of T7p14-mCherry-6xHis<br><br>2. Plasmid maxiprep of T7p14-mCherry-6xHis |
| BL21(DE3) | 1. Protein overexpression<br><br>a. T7p14-mCherry-6xHis<br>b. pET21a-pox5-6xHis<br>c. pET21a-katE-6xHis<br>d. pET21a-ackA-6xHis<br><br>2. Ribosome Purification |

#### Table S6: List of linear DNA fragments (gBlocks, IDT)

Green: Gene coding for protein

**6xHis tag: Purple**

| Name | DNA Sequence |
|---|---|
| pBEST-pox5-6xHis | CCAGCCAGAAAACGACCTTTCTGTGGTGAAACCGGATGCTGCAATTCAGAGCGGCAGCAAGTGGGGGACAGCAGAAGACCTGACCG<br>CCGCAGAGTGGATGTTTGACATGGTGAAGACTATCGCACCATCAGCCAGAAAACCGAATTTTGCTGGGTGGGCTAACGATATCCGCCTG<br>ATGCGTGAACGTGACGGACGTAACCACCGCGACATGTGTGTGCTGTTCCGCTGGGCATGCTGAGCTAACACCGTGCGTGTTGACAATTT<br>TACCTCTGGCGGTGATAATGGTTGCAGCTAGCAATAATTTTGTTTAACTTTAAGAAGGAGATATACCATGGTAATGAAACAGACAAAGCAAAC<br>GAATATCTTGGCGGGGGCAGCAGTGATCAAAGTACTGGAAGCCTGGGGTGTAGACCACTTGTACGGGATTCCAGGTGGGTCTATTAACTC<br>AATTATGGATGCGCTGAGCGCCGAGCGCGACCGGATTCATTATATCCAAGTCCGTCATGAGGAAGTGGGCGCAATGGCCGCGGCTGC<br>CGACGCGAAGCTCACAGGGAAGATCGGGGTGTGCTTCGGCTCTGCGGGTCCGGGTGGCACTCACTTGATGAACGGTCTCTATGACGC<br>ACGCGAGGATCACGTGCCAGTCTTGGCATTAATTGGCCAATTCGGTACAACCGGCATGAATATGGACACATTCCAGGAGATGAATGAGAA<br>TCCGATTTATGCCGACGTTGCGGATTATAATGTTACCGCAGTAAACGCAGCTACTCTCCCGCACGTCATCGACGAAGCAATCCGCCGTG<br>CATACGCCCACCAAGGCGTGGCGGTCGTGCAAATCCCTGTGGATCTCCCGTGGCAGCAAATTCCAGCGGAGGACTGGTACGCAAGTG<br>CTAACTCTTACCAAACGCCACTGTTACCTGAGCCTGACGTTCAAGCAGTTACACGTTTAACACAGACTCTCTTGGCGGCTGAGCGGCCG<br>CTGATCTACTACGGGATTGGCGCACGCAAGGCTGGGAAGGAATTAGAACAATTGTCTAAGACGCTCAAAATCCCACTCATGTCAACTTAT<br>CCAGCAAAAGGTATCGTAGCGGACCGTTATCCAGCATACCTTGGGAGCGCAAACCGCGTGGCGCAAAAACCGGCTAATGAAGCACTG<br>GCGCAAGCCGATGTAGTGCTGTTCGTTGGTAACAACTATCCTTTTGCGGAGGTGTCGAAAGCATTTAAGAACACGCGCTACTTCCTGCAA<br>ATTGATATCGACCCAGCAAAATTAGGGAAGCGGCATAAAACCGATATTGCGGTACTGGCTGATGCCCAGAAGACGCTTGCGGCCATCTTA<br>GCGCAAGTCTCCGAACGCGAAAGCACCCCATGGTGGCAGGCCAATCTGGCTAATGTTAAAAACTGGCGTGCGTATCTTGCGAGCTTAG<br>AGGATAAGCAAGAAGGGCCGCTTCAGGCGTATCAAGTGCTCCGTGCTGTGAATAAGATTGCCGAACCTGATGCAATCTATAGCATTGACG<br>TGGGCGATATCAATCTGAATGCCAATCGGCATCTGAAATTGACCCCGTCCAATCGGCATATTACAAGTAACCTGTTCGCAACTATGGGTGT<br>CGGGATCCCTGGGGCGATTGCAGCCAAACTCAACTACCCTGAGCGCCAAGTTTTAATCTTGCTGGCGACGGCGGTGCAAGTATGACA<br>ATGCAAGATTTAGCGACGCAGGTTCAATACCACTTGCCGGTCATCAACGTAGTCTTTACAAATTGCCAGTACGGGTTCATCAAGGACGAG<br>CAGGAGGACACTAACCAGAATGATTTCATTGGGGTGGAGTTTAATGACATCGACTTTTCCAAAATTGCGGACGGGGTGCATATGCAAGCC<br>TTCCGGGTAAACAAGATCGAACAACTCCCAGATGTTTTCGAACAGGCTAAAGCCATTGCCCAGCATGAACCTGTGTTGATCGATGCTGTC<br>ATTACAGGGGATCGGCCTTTGCCTGCAGAAAAATTGCGCCTTGACAGCGCCACCTCTTCCGCAGCAGACATTGAAGCATTCAAGCAAC<br>GCTATGAAGCACAAGACCTGCAACCGTTGTCGACTTACCTCAAACAGTTCGGGTTAGATGACCTTCAACACCAAATCGGCCAGGGCGG<br>CTTTCATCATCACCATCACCACTAAACAATAACTGAATAGGGGATCCCGACTGGCGAGAGCCAGGTAACGAATGGATCCCCGAGCTCGA<br>GCAAAGCCCGCCGAAAGGCGGGCTTTTCTGTGTCGACCGATGCCCTTGAGAGCCTTCAACCCAGTCAGCTCCTTCCGGTGGGCGCG<br>GGGCATGACTATCGTCGCCGCACTTATGACTGTCTTCTTTATCATGCAACTCGTAGGACAGGTGCCGGCAGCGCTCTTCCGCTTCCTCGC<br>TCACTGACTCGCTGCGCTCGGTCGTTCGGCTGCGGCGAGCGGTATCAGCTCACTCAAAGGCGGTAATACGGTTATCCACAGAATCAGG<br>GGATAACGCAGGAAAGAACATGTGAGCAAAAGG |
| pBEST-ackA-6xHis | CCAGCCAGAAAACGACCTTTCTGTGGTGAAACCGGATGCTGCAATTCAGAGCGGCAGCAAGTGGGGGACAGCAGAAGACCTGACCG<br>CCGCAGAGTGGATGTTTGACATGGTGAAGACTATCGCACCATCAGCCAGAAAACCGAATTTTGCTGGGTGGGCTAACGATATCCGCCTG<br>ATGCGTGAACGTGACGGACGTAACCACCGCGACATGTGTGTGCTGTTCCGCTGGGCATGCTGAGCTAACACCGTGCGTGTTGACAATTT<br>TACCTCTGGCGGTGATAATGGTTGCAGCTAGCAATAATTTTGTTTAACTTTAAGAAGGAGATATACCATGTCTTCCAAGCTGGTTTTAGTGCTG<br>AATTGCGGTAGCTCGAGCCTGAAATTCGCAATTATCGACGCGGTCAACGGCGAGGAGTACTTGTCTGGTCTGGCTGAGTGCTTCCATTTG<br>CCAGAAGCCCGGATTAAGTGGAAGATGGATGGCAACAAGCAGGAGGCCGCGTTGGGCGCTGGGGCTGCTCATTCCGAGGCATTGAAT<br>TTTATTGTGAATACGATTCTCGCCCAAAAACCGGAACTGAGTGCCCAACTCACCGCCATTGGTCACCGGATCGTGCATGGGGGTGAAAA<br>ATATACAAGCTCTGTAGTGATCGATGAATCAGTGATCCAGGGCATCAAAGACGCCGCCTCGTTTGCACCGCTGCACAACCCAGCGCATC<br>TGATTGGCATTGAGGAGGCGTTGAAGTCCTTTCCTCAGCTGAAAGACAAGAATGTTGCGGTATTCGATACCGCGTTCCACCAAACAATGC<br>CGGAGGAGAGCTACCTCTATGCACTCCCTTACAATCTGTACAAAGAACACGGTATCCGCCGCTATGGCGCTCATGGTACAAGCCACTTCT<br>ACGTCACGCAAGAAGCAGCGAAAATGCTCAACAAGCCTGTCGAAGAGTTAAACATTATCACGTGCCACCTTGGTAATGGGGGGTCTGTTT<br>CAGCTATCCGGAACGGCAAGTGCGTGGATACCTCGATGGGCCTGACGCCATTGGAAGGGCTGGTAATGGGCACGCGTTCCGGTGACA<br>TTGACCCGGCTATCATCTTCCATCTCCACGATACATTGGGTATGTCCGTGGACGCAATCAACAAGTTGTTGACAAAAGAGTCAGGGTTATTG<br>GGTTTGACTGAGGTTACAAGTGACTGCCGCTATGTCGAGGATAACTACGCCACGAAAGAAGATGCGAAGCGTGCTATGGACGTGTATTGT<br>CATCGCTTGGCAAAATATATCGGGGCCTACACTGCACTGATGGACGGGCGGTTGGACGCAGTCGTCTTTACCGGCGGCATCGGGGAAA<br>ACGCGGCCATGGTCCGCGAATTGAGCCTGGGTAAGCTCGGCGTTCTTGGTTTTGAAGTGGACCACGAGCGCAACTTGGCAGCTCGCTT<br>TGGGAAGTCAGGGTTCATCAATAAGGAAGGTACTCGTCCTGCGGTTGTAATCCCAACCAATGAGGAATTGGTAATTGCTCAAGATGCGTCG<br>CGTTTGACCGCTCATCATCACCATCACCACTAAACAATAACTGAATAGGGGATCCCGACTGGCGAGAGCCAGGTAACGAATGGATCCCC<br>GAGCTCGAGCAAAGCCCGCCGAAAGGCGGGCTTTTCTGTGTCGACCGATGCCCTTGAGAGCCTTCAACCCAGTCAGCTCCTTCCGGT<br>GGGCGCGGGGCATGACTATCGTCGCCGCACTTATGACTGTCTTCTTTATCATGCAACTCGTAGGACAGGTGCCGGCAGCGCTCTTCCG<br>CTTCCTCGCTCACTGACTCGCTGCGCTCGGTCGTTCGGCTGCGGCGAGCGGTATCAGCTCACTCAAAGGCGGTAATACGGTTATCCAC<br>AGAATCAGGGGATAACGCAGGAAAGAACATGTGAGCAAAAGG |
| pBEST-katE-6xHis | CCAGCCAGAAAACGACCTTTCTGTGGTGAAACCGGATGCTGCAATTCAGAGCGGCAGCAAGTGGGGGACAGCAGAAGACCTGACCG<br>CCGCAGAGTGGATGTTTGACATGGTGAAGACTATCGCACCATCAGCCAGAAAACCGAATTTTGCTGGGTGGGCTAACGATATCCGCCTG<br>ATGCGTGAACGTGACGGACGTAACCACCGCGACATGTGTGTGCTGTTCCGCTGGGCATGCTGAGCTAACACCGTGCGTGTTGACAATTT<br>TACCTCTGGCGGTGATAATGGTTGCAGCTAGCAATAATTTTGTTTAACTTTAAGAAGGAGATATACCATGAGTCAGCATAACGAAAAAAATCC<br>GCATCAACATCAGTCCCCACTCCATGACTCGTCAGAGGCGAAACCTGGTATGGACAGCTTGGCTCCAGAAGACGGCAGCCACCGTCC<br>TGCAGCGGAGCCTACACCGCCAGGCGCCCAACCAACAGCGCCTGGTTCATTAAAAGCGCCGGACACTCGTAACGAGAAATTAAATAG<br>TCTGGAGGACGTGCGCAAGGGCTCAGAGAATTATGCCTTGACTACCAACCAGGGGGTCCGTATCGCTGACGACCAGAATAGCTTACGC<br>GCGGGGAGTCGCGGGCCGACCCTCCTCGAAGACTTTATCTTACGGGAAAAGATTACACACTTTGATCATGAGCGGATTCCGGAGCGCA<br>TCGTACACGCACGGGGGAGTGCAGCCCACGGTTACTTTCAACCTTACAAAAGCTTGTCAGATATCACCAAAGCTGACTTCTTATCTGATC<br>CAAATAAAATTACACCGGTATTCGTGCGGTTCAGTACGGTCCAGGGTGGCGCAGGTTCGGCAGATACAGTACGGGATATCCGGGGTTTT<br>GCGACAAAATTTTATACGGAGGAGGGGATTTTCGATCTGGTCGGCAATAATACGCCTATTTTCTTTATCCAGGACGCGCATAAGTTCCCTGA<br>CTTTGTGCATGCAGTAAAACCAGAACCTCATTGGGCTATTCCACAGGGCCAGAGTGCTCACGATACTTTTTGGGATTACGTCTCTTTGCAG<br>CCTGAAACACTTCACAATGTGATGTGGGCAATGAGCGATCGCGGTATTCCTCGGAGCTACCGGACCATGGAGGGCTTCGGGATCCATAC<br>ATTCCGGCTGATCAATGCCGAAGGCAAGGCCACGTTTGTACGCTTTCATTGGAAACCTCTGGCTGGGAAAGCATCGTTAGTTTGGGACGA<br>GGCGCAGAAGTTAACAGGGCGGGACCCTGATTTTCACCGCCGGGAACTCTGGGAGGCTATTGAGGCTGGGGATTTCCCTGAATACGAA<br>CTCGGCTTTCAGCTTATTCCGGAGGAAGATGAATTTAAATTTGACTTCGATCTCTTAGATCCAACGAAGCTGATCCCGGAAGAATTGGTCCC<br>GGTTCAGCGCGTGGGCAAGATGGTTCTCAATCGCAACCCAGATAACTTCTTCGCGGAAAACGAACAAGCAGCTTTTCATCCTGGGCACA<br>TCGTCCCGGGGCTGGACTTTACAAATGACCCGCTTTTACAAGGGCGCTTATTTTCTTACACCGACACGCAAATCTCACGTCTCGGTGGG<br>CCAAACTTCCATGAGATCCCTATCAATCGGCCTACATGTCCGTATCACAACTTCCAACGTGACGGCATGCACCGTATGGGCATTGATACC<br>AATCCGGCAAATTATGAACCGAACTCAATCAATGACAACTGGCCGCGCGAAACCCCACCGGGCCCAAAGCGGGGGGGTTTCGAGTCA<br>TACCAAGAACGGGTTGAGGGTAATAAAGTGCGCGAGCGGAGCCCTTCGTTTGGTGAATATTATTCCCATCCGCGCCTTTTTTGGCTCAGT<br>CAAACTCCATTTGAGCAGCGGCACATCGTTGACGGGTTCAGCTTTGAGTTATCTAAAGTCGTCCGGCCGTACATTCGCGAACGTGTCGTT<br>GATCAGCTCGCTCATATCGATCTCACCTTAGCCCAGGCTGTGGCCAAGAACCTTGGTATCGAACTTACCGATGACCAATTGAACATCACT<br>CCTCCACCGGATGTGAACGGCCTGAAGAAGGATCCTAGTTTAAGCTTATACGCGATTCCTGATGGTGACGTGAAGGGCCGGGTCGTGGC<br>CATTTTGCTTAACGACGAAGTGCGGTCAGCTGACCTCCTTGCTATTTTAAAGGCACTTAAAGCTAAAGGGGTACATGCCAAGCTGTTGTACT<br>CCCGTATGGGGGAGGTGACAGCTGACGATGGTACCGTGTTGCCAATCGCAGCTACATTTGCAGGCGCACCTAGTCTTACTGTGGATGCC<br>GTTATCGTCCCTTGTGGCAACATTGCCGATATCGCTGACAACGGGGACGCGAATTATTACCTCATGGAAGCTTATAAGCACCTTAAACCTA<br>TTGCTTTAGCAGGGGACGCACGCAAATTTAAGGCGACTATCAAGATCGCCGACCAGGGTGAGGAAGGTATCGTCGAGGCAGACTCCGC<br>CGATGGGTCATTCATGGACGAATTGCTGACCCTGATGGCGGCGCACCGTGTGTGGAGCCGTATTCCGAAGATCGACAAAATTCCTGCTC<br>ATCATCACCATCACCACTAAACAATAACTGAATAGGGGATCCCGACTGGCGAGAGCCAGGTAACGAATGGATCCCCGAGCTCGAGCAA<br>AGCCCGCCGAAAGGCGGGCTTTTCTGTGTCGACCGATGCCCTTGAGAGCCTTCAACCCAGTCAGCTCCTTCCGGTGGGCGCGGGGC |

| | |
|---|---|
| MTF-6xHis | ATGACTATCGTCGCCGCACTTATGACTGTCTTCTTTATCATGCAACTCGTAGGACAGGTGCCGGCAGCGCTCTTCCGCTTCCTCGCTCAC<br>TGACTCGCTGCGCTCGGTCGTTCGGCTGCGGCGAGCGGTATCAGCTCACTCAAAGGCGGTAATACGGTTATCCACAGAATCAGGGGAT<br>AACGCAGGAAAGAACATGTGAGCAAAAGG<br>TAATACGACTCACTATAGGGGAATTGTGAGCGGATAACAATTCCCCTCTAGAAATAATTTTGTTTAACTTTAAGAAGGAGATATACATATGTCA<br>GAATCACTACGTATTATTTTTGCGGGTACACCTGACTTTGCAGCGCGTCATCTCGACGCGCTGTTGTCTTCTGGTCATAACGTCGTTGGCGT<br>GTTCACCCAGCCAGACCGACCGGCAGGACGCGGTAAAAAACTGATGCCCAGCCCGGTTAAAGTTCTGGCTGAGGAAAAAGGTCTGCC<br>CGTTTTTCAACCTGTTTCCCTGCGTCCACAAGAAAACCAGCAACTGGTCGCCGAACTGCAGGCTGATGTTATGGTCGTCGTCGCCTATGG<br>TTTAATTCTGCCGAAAGCAGTGCTGGAGATGCCGCGTCTTGGCTGTATCAACGTTCATGGTTCACTGCTGCCACGCTGGCGCGGTGCTG<br>CACCAATCCAACGCTCACTATGGGCGGGTGATGCAGAAACTGGTGTGACCATTATGCAAATGGATGTCGGTTTAGACACCGGTGATATGC<br>TCTATAAGCTCTCCTGCCCGATTACTGCAGAAGATACCAGTGGTACGCTGTACGACAAGCTGGCAGAGCTTGGCCCACAAGGGCTTATC<br>ACCACGTTGAAACAACTGGCAGACGGCACGGCGAAACCAGAAGTTCAGGACGAAACTCTTGTCACTTACGCCGAGAAGTTGAGTAAAG<br>AAGAAGCGCGTATTGACTGGTCACTTTCGGCAGCACAGCTTGAACGCTGCATTCGCGCTTTCAATCCATGGCCAATGAGCTGGCTGGAA<br>ATTGAAGGACAGCCGGTTAAAGTCTGGAAAGCATCGGTCATTGATACGGCAACCAACGCTGCACCAGGAACGATCCTTGAAGCCAACA<br>AACAAGGCATTCAGGTTGCGACTGGTGATGGCATCCTGAACCTGCTCTCGTTACAACCTGCGGGTAAGAAAGCGATGAGCGCGCAAGA<br>CCTCCTGAACTCTCGTCGGGAATGGTTTGTTCCGGGCAACCGTCTGGTCCTCGAGCACCACCACCACCACCACTGAGATCCGGCTGCT<br>AACAAAGCCCGAAAGGAAGCTGAGTTGGCTGCTGCCACCGCTGAGCAATAACTAGCATAACCCCTTGGGGCCTCTAAACGGGTCTTGA<br>GGGGTTTTTTG |

### Table S7: List of *E. coli* strains used to produce OnePot PURE

| Number | Protein | Protein name | Vector | Strain |
|---|---|---|---|---|
| **1** | **AlaRS** | Alanyl-tRNA synthetase | pQE30 | M15 |
| **2** | **ArgRS** | Arginyl-tRNA synthetase | pET16b | BL21(DE3) |
| **3** | **AsnRS** | Asparaginyl-tRNA synthetase | pQE30 | M15 |
| **4** | **AspRS** | Aspartate-tRNA synthetase | pET21a | BL21(DE3) |
| **5** | **CysRS** | Cysteinyl-tRNA synthetase | pET21a | BL21(DE3) |
| **6** | **GlnRS** | Glutaminyl-tRNA synthetase | pET21a | BL21(DE3) |
| **7** | **GluRS** | Glutamyl-tRNA synthetase | pET21a | BL21(DE3) |
| **8** | **GlyRS** | Glycyl-tRNA synthetase | pET21a | BL21(DE3) |
| **9** | **HisRS** | Histidyl-tRNA synthetase | pET21a | BL21(DE3) |
| **10** | **IleRS** | Isoleucyl-tRNA synthetase | pET21a | BL21(DE3) |
| **11** | **LeuRS** | Leucyl-tRNA synthetase | pET21a | BL21(DE3) |
| **12** | **LysRS** | Lysyl-tRNA synthetase | pET21a | BL21(DE3) |
| **13** | **MetRS** | Methionine--tRNA ligase | pET21a | BL21(DE3) |
| **14** | **PheRS** | Phenylalanyl-tRNA synthetase | pQE30 | M15 |
| **15** | **ProRS** | Prolyl-tRNA synthetase | pET21a | BL21(DE3) |
| **16** | **SerRS** | Seryl-tRNA synthetase | pET21a | BL21(DE3) |
| **17** | **ThrRS** | Threonyl-tRNA synthetase | pQE30 | M15 |
| **18** | **TrpRS** | Tryptophanyl-tRNA synthetase | pET21a | BL21(DE3) |
| **19** | **TyrRS** | Tyrosyl-tRNA synthetase | pET21a | BL21(DE3) |
| **20** | **ValRS** | Valyl-tRNA synthetase | pET21a | BL21(DE3) |
| **21** | **IF1** | Initiation factor 1 | pQE30 | M15 |
| **22** | **IF2** | Initiation factor 2 | pQE30 | M15 |
| **23** | **IF3** | Initiation factor 3 | pQE30 | M15 |
| **24** | **EF-G** | Elongation factor G | pQE60 | M15 |
| **25** | **EF-Tu** | Elongation factor Tu | pQE60 | M15 |
| **26** | **EF-Ts** | Elongation factor Ts | pQE60 | M15 |
| **27** | **RF1** | Release factor 1 | pQE30 | M15 |
| **28** | **RF2** | Release factor 2 | pET15b | BL21(DE3) |
| **29** | **RF3** | Release factor 3 | pQE30 | M15 |
| **30** | **RRF** | Ribosome recycling factor | pQE60 | M15 |
| **31** | **MTF** | Methionyl-tRNA formyltransferase | pET21a | BL21(DE3) |
| **32** | **CK** | Creatine kinase | pQE30 | M15 |
| **33** | **MK** | Adenylate kinase (Myokinase) | pET21a | BL21(DE3) |
| **34** | **NDK** | Nucleotide diphosphate kinase | pQE30 | M15 |
| **35** | **PPiase** | Inorganic pyrophosphatase | pET21a | BL21(DE3) |
| **36** | **T7 RNAP** | T7 RNA polymerase | pQE30 | M15 |

#### Table S8: List of primers

| Name | Sequence (5' -----> 3') | Details |
|---|---|---|
| P_01_FWD | CTTTAAGAAGGAGATATACATATGGTAATGAA ACAGACAAAGC | Gibson assembly primer for lifting the pox5-6xHis gene from pBEST-pox5-6xHis with overhangs complementary to pET21a backbone |
| P_01_REV | GCTTTGTTAGCAGCCGGATCTTAGTGGTGAT GGTGATGAT | Gibson assembly primer for lifting the pox5-6xHis gene from pBEST-pox5-6xHis with overhangs complementary to pET21a backbone |
| P_02_FWD | ATCATCACCATCACCACTAAGATCCGGCTG CTAACAAAGC | Gibson assembly primer for lifting the pET21a backbone from pET21a-AspRS-6xHis (PURE plasmid) with overhangs complementary to pox5-6xHis gene region |
| P_02_REV | TTTGTCTGTTTCATTACCATATGTATATCTCCTT CTTAAAGTTAAACAA | Gibson assembly primer for lifting the pET21a backbone from pET21a-AspRS-6xHis (PURE plasmid) with overhangs complementary to pox5-6xHis gene region |
| P_03_FWD | TTTAAGAAGGAGATATACATATGTCTTCCAAG CTGGTTTT | Gibson assembly primer for lifting the ackA-6xHis gene from pBEST-ackA-6xHis with overhangs complementary to pET21a backbone |
| P_03_REV | GCTTTGTTAGCAGCCGGATCTTAGTGGTGAT GGTGATGAT | Gibson assembly primer for lifting the ackA-6xHis gene from pBEST-ackA-6xHis with overhangs complementary to pET21a backbone |
| P_04_FWD | ATCATCACCATCACCACTAAGATCCGGCTG CTAACAAAGC | Gibson assembly primer for lifting the pET21a backbone from pET21a-AspRS-6xHis (PURE plasmid) with overhangs complementary to ackA-6xHis gene region |
| P_04_REV | AAAACCAGCTTGGAAGACATATGTATATCTC CTTCTTAAAGTTAAACAA | Gibson assembly primer for lifting the pET21a backbone from pET21a-AspRS-6xHis (PURE plasmid) with overhangs complementary to ackA-6xHis gene region |
| P_05_FWD | ACTTTAAGAAGGAGATATACATATGAGTCAG CATAACGAAAAA | Gibson assembly primer for lifting the katE-6xHis gene from pBEST-katE-6xHis with overhangs complementary to pET21a backbone |
| P_05_REV | GCTTTGTTAGCAGCCGGATCTTAGTGGTGAT GGTGATGAT | Gibson assembly primer for lifting the katE-6xHis gene from pBEST-katE-6xHis with overhangs complementary to pET21a backbone |
| P_06_FWD | ATCATCACCATCACCACTAAGATCCGGCTG CTAACAAAGC | Gibson assembly primer for lifting the pET21a backbone from pET21a-AspRS-6xHis (PURE plasmid) with overhangs complementary to katE-6xHis gene region |
| P_06_REV | TTTTCGTTATGCTGACTCATATGTATATCTCCT TCTTAAAGTTAAACAA | Gibson assembly primer for lifting the pET21a backbone from pET21a-AspRS-6xHis (PURE plasmid) with overhangs complementary to katE-6xHis gene region |
| P_07_FWD | TAATACGACTCACTATAGGGG | Primer to amplify linear MTF-6xHis DNA fragment |

| | | |
|---|---|---|
| P_07_REV | CAAAAAACCCCTCAAGACCC | Primer to amplify linear MTF-6xHis DNA fragment |
| P_08_FWD | ACCCCTTGGGGCCTCTAAA | Primer to lift pET21a backbone from pET21a-ackA-6xHis plasmid for gibson assembly with MTF-6xHis |
| P_08_REV | TTGTTATCCGCTCACAATTCCCC | Primer to lift pET21a backbone from pET21a-ackA-6xHis plasmid for gibson assembly with MTF-6xHis |
| P_09_FWD | GCATGGACGAGCTGTACAAGCACCACCACCACCATCACTAAGATCCGGCTGCTAACAA | Gibson assembly primer for lifting the T7p14 backbone from T7p14-deGFP plasmid with overhangs complementary to mCherry-6xHis gene region |
| P_09_REV | TCTTCGCCCTTGCTCACCATGGTATATCTCCTTCTTAAAGTTAAACA | Gibson assembly primer for lifting the T7p14 backbone from T7p14-deGFP plasmid with overhangs complementary to mCherry gene region |
| P_10_FWD | CTTTAAGAAGGAGATATACCATGGTGAGCAAGGGCGAAGA | Gibson assembly primer for lifting the mCherry gene from pLtetO-mCherry with overhangs complementary to T7p14 backbone |
| P_10_REV | TTGTTAGCAGCCGGATCTTAGTGATGGTGGTGGTGGTGCTTGTACAGCTCGTCCATGC | Gibson assembly primer for lifting the mCherry gene from pLtetO-mCherry and inserting C-terminal 6xHis with overhangs complementary to T7p14 backbone |

#### Table S9: List of plasmids (excluding PURE plasmids)

Blue: T7 Promoter    Red: RBS    Green: Gene coding for protein    **Bold: T7 terminator**    **6xHis tag: Purple**

| Plasmid | DNA Sequence |
|---|---|
| T7p14-mCherry-6xHis | TTAGATTTCATACACGGTGCCTGACTGCGTTAGCAATTTAACTGTGATAAACTACCGCATTAAAGCTTATCGATGATAAGCTGTCAAACATG<br>AGAATTCGTAATCATGTCATAGCTGTTTCCTGTGTGAAATTGTTATCCGCTCACAATTCCACACAACATACGAGCCGGAAGCATAAAGTGT<br>AAAGCCTGGGGTGCCTAATGAGTGAGCTAACTCACATTAATTGCGTTGCGCTCACTGCCCGCTTTCCAGTCGGGAAACCTGTCGTGC<br>CAGCTGCATTAATGAATCGGCCAACGCGCGGGGAGAGGCGGTTTGCGTATTGGGCGCTCTTCCGCTTCCTCGCTCACTGACTCGCT<br>GCGCTCGGTCGTTCGGCTGCGGCGAGCGGTATCAGCTCACTCAAAGGCGGTAATACGGTTATCCACAGAATCAGGGGATAACGCAG<br>GAAAGAACATGTGAGCAAAAGGCCAGCAAAAGGCCAGGAACCGTAAAAAGGCCGCGTTGCTGGCGTTTTTCCATAGGCTCCGCCC<br>CCCTGACGAGCATCACAAAAATCGACGCTCAAGTCAGAGGTGGCGAAACCCGACAGGACTATAAAGATACCAGGCGTTTCCCCCTG<br>GAAGCTCCCTCGTGCGCTCTCCTGTTCCGACCCTGCCGCTTACCGGATACCTGTCCGCCTTTCTCCCTTCGGGAAGCGTGGCGCTT<br>TCTCATAGCTCACGCTGTAGGTATCTCAGTTCGGTGTAGGTCGTTCGCTCCAAGCTGGGCTGTGTGCACGAACCCCCCGTTCAGCCC<br>GACCGCTGCGCCTTATCCGGTAACTATCGTCTTGAGTCCAACCCGGTAAGACACGACTTATCGCCACTGGCAGCAGCCACTGGTAAC<br>AGGATTAGCAGAGCGAGGTATGTAGGCGGTGCTACAGAGTTCTTGAAGTGGTGGCCTAACTACGGCTACACTAGAAGGACAGTATTTG<br>GTATCTGCGCTCTGCTGAAGCCAGTTACCTTCGGAAAAAGAGTTGGTAGCTCTTGATCCGGCAAACAAACCACCGCTGGTAGCGGTG<br>GTTTTTTTGTTTGCAAGCAGCAGATTACGCGCAGAAAAAAAGGATCTCAAGAAGATCCTTTGATCTTTTCTACGGGGTCTGACGCTCAGT<br>GGAACGAAAACTCACGTTAAGGGATTTTGGTCATGAGATTATCAAAAAGGATCTTCACCTAGATCCTTTTAAATTAAAAATGAAGTTTTAAA<br>TCAATCTAAAGTATATATGAGTAAACTTGGTCTGACAGTTACCAATGCTTAATCAGTGAGGCACCTATCTCAGCGATCTGTCTATTTCGTTC<br>ATCCATAGTTGCCTGACTCCCCGTCGTGTAGATAACTACGATACGGGAGGGCTTACCATCTGGCCCCAGTGCTGCAATGATACCGCG<br>AGACCCACGCTCACCGGCTCCAGATTTATCAGCAATAAACCAGCCAGCCGGAAGGGCCGAGCGCAGAAGTGGTCCTGCAACTTTA<br>TCCGCCTCCATCCAGTCTATTAATTGTTGCCGGGAAGCTAGAGTAAGTAGTTCGCCAGTTAATAGTTTGCGCAACGTTGTTGCCATTGCT<br>ACAGGCATCGTGGTGTCACGCTCGTCGTTTGGTATGGCTTCATTCAGCTCCGGTTCCCAACGATCAAGGCGAGTTACATGATCCCCCA<br>TGTTGTGCAAAAAAGCGGTTAGCTCCTTCGGTCCTCCGATCGTTGTCAGAAGTAAGTTGGCCGCAGTGTTATCACTCATGGTTATGGCA<br>GCACTGCATAATTCTCTTACTGTCATGCCATCCGTAAGATGCTTTTCTGTGACTGGTGAGTACTCAACCAAGTCATTCTGAGAATAGTGTAT<br>GCGGCGACCGAGTTGCTCTTGCCCGGCGTCAATACGGGATAATACCGCGCCACATAGCAGAACTTTAAAAGTGCTCATCATTGGAAA<br>ACGTTCTTCGGGGCGAAAACTCTCAAGGATCTTACCGCTGTTGAGATCCAGTTCGATGTAACCCACTCGTGCACCCAACTGATCTTCA<br>GCATCTTTTACTTTCACCAGCGTTTCTGGGTGAGCAAAAACAGGAAGGCAAAATGCCGCAAAAAAGGGAATAAGGGCGACACGGAAA<br>TGTTGAATACTCATACTCTTCCTTTTTCAATATTATTGAAGCATTTATCAGGGTTATTGTCTCATGAGCGGATACATATTTGAATGTATTTAGAA<br>AAATAAACAAATAGGGGTTCCGCGCACATTTCCCCGAAAAGTGCCACCTGACGTCTAAGAAACCATTATTATCATGACATTAACCTATAA<br>AAATAGGCGTATCACGAGGCCCTTTCGTCTCGCGCGTTTCGGTGATGACGGTGAAAACCTCTGACACATGCAGCTCCCGGAGACGG<br>TCACAGCTTGTCTGTAAGCGGATGCCGGGAGCAGACAAGCCCGTCAGGGCGCGTCAGCGGGTGTTGGCGGGTGTCGGGGCTGGC<br>TTAACTATGCGGCATCAGAGCAGATTGTACTGAGAGTGCACCATATATGCGGTGTGAAATACCGCACAGATGCGTAAGGAGAAAATACC<br>GCATCAGGCGCCATTCGCCATTCAGGCTGCGCAACTGTTGGGAAGGGCGATCGGTGCGGGCCTCTTCGCTATTACGCCAGCTGGC<br>GAAAGGGGGATGTGCTGCAAGGCGATTAAGTTGGGTAACGCCAGGGTTTTCCCAGTCACGACGTTGTAAAACGACGGCCAGTGCCA<br>AGCTTGCATGCAAGGAGATGGCGCCCAACAGTCCCCCGGCCACGGGGCCTGCCACCATACCCACGCCGAAACAAGCGCTCATG<br>AGCCCGAAGTGGCGAGCCCGATCTTCCCCATCGGTGATGTCGGCGATATAGGCGCCAGCAACCGCACCTGTGGCGCCGGTGATG<br>CCGGCCACGATGCGTCCGGCGTAGAGGATCGAGATCTCGATCCCGCGAAATTAATACGACTCACTATAGGGAGACCACAACGGTTT<br>CCCTCTAGAAATAATTTTGTTTAACTTTAAGAAGGAGATATACCATGGTGAGCAAGGGCGAAGAAGATAACATGGCCATCATCAAGGAGT<br>TCATGCGCTTCAAGGTGCACATGGAGGGCTCCGTGAACGGCCACGAGTTCGAGATCGAGGGCGAGGGCGAGGGCCGCCCCTAC<br>GAGGGCACCCAGACCGCCAAGCTGAAGGTGACCAAGGGTGGCCCCCTGCCCTTCGCCTGGGACATCCTGTCCCCTCAGTTCATG<br>TACGGCTCCAAGGCCTACGTGAAGCACCCCGCCGACATCCCCGACTACTTGAAGCTGTCCTTCCCCGAGGGCTTCAAGTGGGAGC<br>GCGTGATGAACTTCGAGGACGGCGGCGTGGTGACCGTGACCCAGGACTCCTCCCTGCAGGACGGCGAGTTCATCTACAAGGTGAA<br>GCTGCGCGGCACCAACTTCCCCTCCGACGGCCCCGTAATGCAGAAGAAGACCATGGGCTGGGAGGCCTCCTCCGAGCGGATGTA<br>CCCCGAGGACGGCGCCCTGAAGGGCGAGATCAAGCAGAGGCTGAAGCTGAAGGACGGCGGCCACTACGACGCTGAGGTCAAGA<br>CCACCTACAAGGCCAAGAAGCCCGTGCAGCTGCCCGGCGCCTACAACGTCAACATCAAGTTGGACATCACCTCCCACAACGAGG<br>ACTACACCATCGTGGAACAGTACGAACGCGCCGAGGGCCGCCACTCCACCGGCGGCATGGACGAGCTGTACAAGCACCACCAC<br>CACCATCACTAAGATCCGGCTGCTAACAAAGCCCGAAAGGAAGCTGAGTTGGCTGCTGCCACCGCTGAGCAATAAC**TAGCATAACC**<br>**CCTTGGGGCCTCTAAACGGGTCTTGAGGGGTTTTTTG**CTGAAAGGAGGAACTATATCCGGATATCCACAGGACGGGTGTGGTCGCC<br>ATGATCGCGTAGTCGATAGTGGCTCCAAGTAGCGAAGCGAGCAGGACTGGGCGGCGGCCAAAGCGGTCGGACAGTGCTCCGAGA<br>ACGGGTGCGCATAGAAATTGCATCAACGCATATAGCGCTAGCAGCACGCCATAGTGACTGGCGATGCTGTCGGAATGGACGATATCC<br>CGCAAGAGGCCCGGCAGTACCGGCATAACCAAGCCTATGCCTACAGCATCCAGGGTGACGGTGCCGAGGATGACGATGAGCGCA<br>TTG |
| pET21a-pox5-6xHis | GAGTCCACGTTCTTTAATAGTGGACTCTTGTTCCAAACTGGAACAACACTCAACCCTATCTCGGTCTATTCTTTTGATTTATAAGGGATTTT<br>GCCGATTTCGGCCTATTGGTTAAAAAATGAGCTGATTTAACAAAAATTTAACGCGAATTTTAACAAAATATTAACGCTTACAATTTAGGTGG<br>CACTTTTCGGGGAAATGTGCGCGGAACCCCTATTTGTTTATTTTTCTAAATACATTCAAATATGTATCCGCTCATGAGACAATAACCCTGAT<br>AAATGCTTCAATAATATTGAAAAAGGAAGAGTATGAGTATTCAACATTTCCGTGTCGCCCTTATTCCCTTTTTTGCGGCATTTTGCCTTCCT<br>GTTTTTGCTCACCCAGAAACGCTGGTGAAAGTAAAAGATGCTGAAGATCAGTTGGGTGCACGAGTGGGTTACATCGAACTGGATCTCA<br>ACAGCGGTAAGATCCTTGAGAGTTTTCGCCCCGAAGAACGTTTTCCAATGATGAGCACTTTTAAAGTTCTGCTATGTGGCGCGGTATTAT<br>CCCGTATTGACGCCGGGCAAGAGCAACTCGGTCGCCGCATACACTATTCTCAGAATGACTTGGTTGAGTACTCACCAGTCACAGAAA<br>AGCATCTTACGGATGGCATGACAGTAAGAGAATTATGCAGTGCTGCCATAACCATGAGTGATAACACTGCGGCCAACTTACTTCTGACA<br>ACGATCGGAGGACCGAAGGAGCTAACCGCTTTTTTGCACAACATGGGGGATCATGTAACTCGCCTTGATCGTTGGGAACCGGAGCTG<br>AATGAAGCCATACCAAACGACGAGCGTGACACCACGATGCCTGCAGCAATGGCAACAACGTTGCGCAAACTATTAACTGGCGAACT<br>ACTTACTCTAGCTTCCCGGCAACAATTAATAGACTGGATGGAGGCGGATAAAGTTGCAGGACCACTTCTGCGCTCGGCCCTTCCGGC<br>TGGCTGGTTTATTGCTGATAAATCTGGAGCCGGTGAGCGTGGGTCTCGCGGTATCATTGCAGCACTGGGGCCAGATGGTAAGCCCTC<br>CCGTATCGTAGTTATCTACACGACGGGGAGTCAGGCAACTATGGATGAACGAAATAGACAGATCGCTGAGATAGGTGCCTCACTGATTA<br>AGCATTGGTAACTGTCAGACCAAGTTTACTCATATATACTTTAGATTGATTTAAAACTTCATTTTTAATTTAAAAGGATCTAGGTGAAGATCCTT<br>TTTGATAATCTCATGACCAAAATCCCTTAACGTGAGTTTTCGTTCCACTGAGCGTCAGACCCCGTAGAAAAGATCAAAGGATCTTCTTGA<br>GATCCTTTTTTTCTGCGCGTAATCTGCTGCTTGCAAACAAAAAAACCACCGCTACCAGCGGTGGTTTGTTTGCCGGATCAAGAGCTAC<br>CAACTCTTTTTCCGAAGGTAACTGGCTTCAGCAGAGCGCAGATACCAAATACTGTCCTTCTAGTGTAGCCGTAGTTAGGCCACCACTTC<br>AAGAACTCTGTAGCACCGCCTACATACCTCGCTCTGCTAATCCTGTTACCAGTGGCTGCTGCCAGTGGCGATAAGTCGTGTCTTACCG<br>GGTTGGACTCAAGACGATAGTTACCGGATAAGGCGCAGCGGTCGGGCTGAACGGGGGGTTCGTGCACACAGCCCAGCTTGGAGC<br>GAACGACCTACACCGAACTGAGATACCTACAGCGTGAGCTATGAGAAAGCGCCACGCTTCCCGAAGGGAGAAAGGCGGACAGGTA<br>TCCGGTAAGCGGCAGGGTCGGAACAGGAGAGCGCACGAGGGAGCTTCCAGGGGGAAACGCCTGGTATCTTTATAGTCCTGTCGGG<br>TTTCGCCACCTCTGACTTGAGCGTCGATTTTTGTGATGCTCGTCAGGGGGGCGGAGCCTATGGAAAAACGCCAGCAACGCGGCCTTT<br>TTACGGTTCCTGGCCTTTTGCTGGCCTTTTGCTCACATGTTCTTTCCTGCGTTATCCCCTGATTCTGTGGATAACCGTATTACCGCCTTTG<br>AGTGAGCTGATACCGCTCGCCGCAGCCGAACGACCGAGCGCAGCGAGTCAGTGAGCGAGGAAGCGGAAGAGCGCCTGATGCGG<br>TATTTTCTCCTTACGCATCTGTGCGGTATTTCACACCGCAATGGTGCACTCTCAGTACAATCTGCTCTGATGCCGCATAGTTAAGCCAGT<br>ATACACTCCGCTATCGCTACGTGACTGGGTCATGGCTGCGCCCCGACACCCGCCAACACCCGCTGACGCGCCCTGACGGGCTTG<br>TCTGCTCCCGGCATCCGCTTACAGACAAGCTGTGACCGTCTCCGGGAGCTGCATGTGTCAGAGGTTTTCACCGTCATCACCGAAACG<br>CGCGAGGCAGCTGCGGTAAAGCTCATCAGCGTGGTCGTGAAGCGATTCACAGATGTCTGCCTGTTCATCCGCGTCCAGCTCGTTGA<br>GTTTCTCCAGAAGCGTTAATGTCTGGCTTCTGATAAAGCGGGCCATGTTAAGGGCGGTTTTTTCCTGTTTGGTCACTGATGCCTCCGTGT<br>AAGGGGGATTTCTGTTCATGGGGGTAATGATACCGATGAAACGAGAGAGGATGCTCACGATACGGGTTACTGATGATGAACATGCCCG<br>GTTACTGGAACGTTGTGAGGGTAAACAACTGGCGGTATGGATGCGGCGGGACCAGAGAAAAATCACTCAGGGTCAATGCCAGCGCT<br>TCGTTAATACAGATGTAGGTGTTCCACAGGGTAGCCAGCAGCATCCTGCGATGCAGATCCGGAACATAATGGTGCAGGGCGCTGACT<br>TCCGCGTTTCCAGACTTTACGAAACACGGAAACCGAAGACCATTCATGTTGTTGCTCAGGTCGCAGACGTTTTGCAGCAGCAGTCGCT |

pET21a-ackA-6xHis

TCACGTTCGCTCGCGTATCGGTGATTCATTCTGCTAACCAGTAAGGCAACCCCGCCAGCCTAGCCGGGTCCTCAACGACAGGAGCA
CGATCATGCGCACCCGTGGGGCCGCCATGCCGGCGATAATGGCCTGCTTCTCGCCGAAACGTTTGGTGGCGGGACCAGTGACGA
AGGCTTGAGCGAGGGCGTGCAAGATTCCGAATACCGCAAGCGACAGGCCGATCATCGTCGCGCTCCAGCGAAAGCGGTCCTCGC
CGAAAATGACCCAGAGCGCTGCCGGCACCTGTCCTACGAGTTGCATGATAAAGAAGACAGTCATAAGTGCGGCGACGATAGTCATG
CCCCGCGCCCACCGGAAGGAGCTGACTGGGTTGAAGGCTCTCAAGGGCATCGGTCGAGATCCCGGTGCCTAATGAGTGAGCTAAC
TTACATTAATTGCGTTGCGCTCACTGCCCGCTTTCCAGTCGGGAAACCTGTCGTGCCAGCTGCATTAATGAATCGGCCAACGCGCGG
GGAGAGGCGGTTTGCGTATTGGGCGCCAGGGTGGTTTTTCTTTTCACCAGTGAGACGGGCAACAGCTGATTGCCCTTCACCGCCTG
GCCCTGAGAGAGTTGCAGCAAGCGGTCCACGCTGGTTTGCCCCAGCAGGCGAAAATCCTGTTTGATGGTGGTTAACGGCGGGATAT
AACATGAGCTGTCTTCGGTATCGTCGTATCCCACTACCGAGATATCCGCACCAACGCGCAGCCCGGACTCGGTAATGGCGCGCATT
GCGCCCAGCGCCATCTGATCGTTGGCAACCAGCATCGCAGTGGGAACGATGCCCTCATTCAGCATTTGCATGGTTTGTTGAAAACCG
GACATGGCACTCCAGTCGCCTTCCCGTTCCGCTATCGGCTGAATTTGATTGCGAGTGAGATATTTATGCCAGCCAGCCAGACGCAGA
CGCGCCGAGACAGAACTTAATGGGCCCGCTAACAGCGCGATTTGCTGGTGACCCAATGCGACCAGATGCTCCACGCCCAGTCGC
GTACCGTCTTCATGGGAGAAAATAATACTGTTGATGGGTGTCTGGTCAGAGACATCAAGAAATAACGCCGGAACATTAGTGCAGGCAG
CTTCCACAGCAATGGCATCCTGGTCATCCAGCGGATAGTTAATGATCAGCCCACTGACGCGTTGCGCGAGAAGATTGTGCACCGCC
GCTTTACAGGCTTCGACGCCGCTTCGTTCTACCATCGACACCACCACGCTGGCACCCAGTTGATCGGCGCGAGATTTAATCGCCGC
GACAATTTGCGACGGCGCGTGCAGGGCCAGACTGGAGGTGGCAACGCCAATCAGCAACGACTGTTTGCCCGCCAGTTGTTGTGCC
ACGCGGTTGGGAATGTAATTCAGCTCCGCCATCGCCGCTTCCACTTTTTCCCGCGTTTTCGCAGAAACGTGGCTGGCCTGGTTCACC
ACGCGGGAAACGGTCTGATAAGAGACACCGGCATACTCTGCGACATCGTATAACGTTACTGGTTTCACATTCACCACCCTGAATTGAC
TCTCTTCCGGGCGCTATCATGCCATACCGCGAAAGGTTTTGCGCCATTCGATGGTGTCCGGGATCTCGACGCTCTCCCTTATGCGACT
CCTGCATTAGGAAGCAGCCCAGTAGTAGGTTGAGGCCGTTGAGCACCGCCGCCGCAAGGAATGGTGCATGCAAGGAGATGGCGC
CCAACAGTCCCCCGGCCACGGGGCCTGCCACCATACCCACGCCGAAACAAGCGCTCATGAGCCCGAAGTGGCGAGCCCGATCT
TCCCCATCGGTGATGTCGGCGATATAGGCGCCAGCAACCGCACCTGTGGCGCCGGTGATGCCGGCCACGATGCGTCCGGCGTAG
AGGATCGAGATCTCGATCCCGCGAAATTAATACGACTCACTATAGGGGAATTGTGAGCGGATAACAATTCCCCTCTAGAAATAATTTTGT
TTAACTTTAAGAAGGAGATATACATATGGTAATGAAACAGACAAAGCAAACGAATATCTTGGCGGGGGCAGCAGTGATCAAAGTACTGG
AAGCCTGGGGTGTAGACCACTTGTACGGGATTCCAGGTGGGTCTATTAACTCAATTATGGATGCGCTGAGCGCCGAGCGCGACCGG
ATTCATTATATCCAAGTCCGTCATGAGGAAGTGGGCGCAATGGCCGCGGCTGCCGACGCGAAGCTCACAGGGAAGATCGGGGTGTG
CTTCGGCTCTGCGGGTCCGGGTGGCACTCACTTGATGAACGGTCTCTATGACGCACGCGAGGATCACGTGCCAGTCTTGGCATTAAT
TGGCCAATTCGGTACAACCGGCATGAATATGGACACATTCCAGGAGATGAATGAGAATCCGATTTATGCCGACGTTGCGGATTATAATG
TTACCGCAGTAAACGCAGCTACTCTCCCGCACGTCATCGACGAAGCAATCCGCCGTGCATACGCCCACCAAGGCGTGGCGGTCGT
GCAAATCCCTGTGGATCTCCCGTGGCAGCAAATTCCAGCGGAGGACTGGTACGCAAGTGCTAACTCTTACCAAACGCCACTGTTACC
TGAGCCTGACGTTCAAGCAGTTACACGTTTAACACAGACTCTCTTGGCGGCTGAGCGGCCGCTGATCTACTACGGGATTGGCGCACG
CAAGGCTGGGAAGGAATTAGAACAATTGTCTAAGACGCTCAAAATCCCACTCATGTCAACTTATCCAGCAAAAGGTATCGTAGCGGAC
CGTTATCCAGCATACCTTGGGAGCGCAAACCGCGTGGCGCAAAAACCGGCTAATGAAGCACTGGCGCAAGCCGATGTAGTGCTGTT
CGTTGGTAACAACTATCCTTTTGCGGAGGTGTCGAAAGCATTTAAGAACACGCGCTACTTCCTGCAAATTGATATCGACCCAGCAAAAT
TAGGGAAGCGGCATAAAACCGATATTGCGGTACTGGCTGATGCCCAGAAGACGCTTGCGGCCATCTTAGCGCAAGTCTCCGAACGC
GAAAGCACCCCATGGTGGCAGGCCAATCTGGCTAATGTTAAAAACTGGCGTGCGTATCTTGCGAGCTTAGAGGATAAGCAAGAAGGG
CCGCTTCAGGCGTATCAAGTGCTCCGTGCTGTGAATAAGATTGCCGAACCTGATGCAATCTATAGCATTGACGTGGGCGATATCAATCT
GAATGCCAATCGGCATCTGAAATTGACCCCGTCCAATCGGCATATTACAAGTAACCTGTTCGCAACTATGGGTGTCGGGATCCCTGGG
GCGATTGCAGCCAAACTCAACTACCCTGAGCGCCAAGTTTTTAATCTTGCTGGCGACGGCGGTGCAAGTATGACAATGCAAGATTTAG
CGACGCAGGTTCAATACCACTTGCCGGTCATCAACGTAGTCTTTACAAATTGCCAGTACGGGTTCATCAAGGACGAGCAGGAGGACA
CTAACCAGAATGATTTCATTGGGGTGGAGTTTAATGACATCGACTTTTCCAAAATTGCGGACGGGGTGCATATGCAAGCCTTCCGGGTA
AACAAGATCGAACAACTCCCAGATGTTTTCGAACAGGCTAAAGCCATTGCCCAGCATGAACCTGTGTTGATCGATGCTGTCATTACAGG
GGATCGGCCTTTGCCTGCAGAAAAATTGCGCCTTGACAGCGCCACCTCTTCCGCAGCAGACATTGAAGCATTCAAGCAACGCTATGA
AGCACAAGACCTGCAACCGTTGTCGACTTACCTCAAACAGTTCGGGTTAGATGACCTTCAACACCAAATCGGCCAGGGCGGCTTTCA
TCATCACCATCACCACTAAGATCCGGCTGCTAACAAAGCCCGAAAGGAAGCTGAGTTGGCTGCTGCCACCGCTGAGCAATAAC**TAG**
**CATAACCCCTTGGGGCCTCTAAACGGGTCTTGAGGGGTTTTTTG**CTGAAAGGAGGAACTATATCCGGATTGGCGAATGGGACGCG
CCCTGTAGCGGCGCATTAAGCGCGGCGGGTGTGGTGGTTACGCGCAGCGTGACCGCTACACTTGCCAGCGCCCTAGCGCCCGCT
CCTTTCGCTTTCTTCCCTTCCTTTCTCGCCACGTTCGCCGGCTTTCCCCGTCAAGCTCTAAATCGGGGGCTCCCTTTAGGGTTCCGAT
TTAGTGCTTTACGGCACCTCGACCCCAAAAAACTTGATTAGGGTGATGGTTCACGTAGTGGGCCATCGCCCTGATAGACGGTTTTTCG
CCCTTTGACGTTG
GAGTCCACGTTCTTTAATAGTGGACTCTTGTTCCAAACTGGAACAACACTCAACCCTATCTCGGTCTATTCTTTTGATTTATAAGGGATTTT
GCCGATTTCGGCCTATTGGTTAAAAAATGAGCTGATTTAACAAAAATTTAACGCGAATTTTAACAAAATATTAACGCTTACAATTTAGGTGG
CACTTTTCGGGGAAATGTGCGCGGAACCCCTATTTGTTTATTTTTCTAAATACATTCAAATATGTATCCGCTCATGAGACAATAACCCTGAT
AAATGCTTCAATAATATTGAAAAAGGAAGAGTATGAGTATTCAACATTTCCGTGTCGCCCTTATTCCCTTTTTTGCGGCATTTTGCCTTCCT
GTTTTTGCTCACCCAGAAACGCTGGTGAAAGTAAAAGATGCTGAAGATCAGTTGGGTGCACGAGTGGGTTACATCGAACTGGATCTCA
ACAGCGGTAAGATCCTTGAGAGTTTTCGCCCCGAAGAACGTTTTCCAATGATGAGCACTTTTAAAGTTCTGCTATGTGGCGCGGTATTAT
CCCGTATTGACGCCGGGCAAGAGCAACTCGGTCGCCGCATACACTATTCTCAGAATGACTTGGTTGAGTACTCACCAGTCACAGAAA
AGCATCTTACGGATGGCATGACAGTAAGAGAATTATGCAGTGCTGCCATAACCATGAGTGATAACACTGCGGCCAACTTACTTCTGACA
ACGATCGGAGGACCGAAGGAGCTAACCGCTTTTTTGCACAACATGGGGGATCATGTAACTCGCCTTGATCGTTGGGAACCGGAGCTG
AATGAAGCCATACCAAACGACGAGCGTGACACCACGATGCCTGCAGCAATGGCAACAACGTTGCGCAAACTATTAACTGGCGAACT
ACTTACTCTAGCTTCCCGGCAACAATTAATAGACTGGATGGAGGCGGATAAAGTTGCAGGACCACTTCTGCGCTCGGCCCTTCCGGC
TGGCTGGTTTATTGCTGATAAATCTGGAGCCGGTGAGCGTGGGTCTCGCGGTATCATTGCAGCACTGGGGCCAGATGGTAAGCCCTC
CCGTATCGTAGTTATCTACACGACGGGGAGTCAGGCAACTATGGATGAACGAAATAGACAGATCGCTGAGATAGGTGCCTCACTGATTA
AGCATTGGTAACTGTCAGACCAAGTTTACTCATATATACTTTAGATTGATTTAAAACTTCATTTTTAATTTAAAAGGATCTAGGTGAAGATCCTT
TTTGATAATCTCATGACCAAAATCCCTTAACGTGAGTTTTCGTTCCACTGAGCGTCAGACCCCGTAGAAAAGATCAAAGGATCTTCTTGA
GATCCTTTTTTTCTGCGCGTAATCTGCTGCTTGCAAACAAAAAAACCACCGCTACCAGCGGTGGTTTGTTTGCCGGATCAAGAGCTAC
CAACTCTTTTTCCGAAGGTAACTGGCTTCAGCAGAGCGCAGATACCAAATACTGTCCTTCTAGTGTAGCCGTAGTTAGGCCACCACTTC
AAGAACTCTGTAGCACCGCCTACATACCTCGCTCTGCTAATCCTGTTACCAGTGGCTGCTGCCAGTGGCGATAAGTCGTGTCTTACCG
GGTTGGACTCAAGACGATAGTTACCGGATAAGGCGCAGCGGTCGGGCTGAACGGGGGGTTCGTGCACACAGCCCAGCTTGGAGC
GAACGACCTACACCGAACTGAGATACCTACAGCGTGAGCTATGAGAAAGCGCCACGCTTCCCGAAGGGAGAAAGGCGGACAGGTA
TCCGGTAAGCGGCAGGGTCGGAACAGGAGAGCGCACGAGGGAGCTTCCAGGGGGAAACGCCTGGTATCTTTATAGTCCTGTCGGG
TTTCGCCACCTCTGACTTGAGCGTCGATTTTTGTGATGCTCGTCAGGGGGGCGGAGCCTATGGAAAAACGCCAGCAACGCGGCCTTT
TTACGGTTCCTGGCCTTTTGCTGGCCTTTTGCTCACATGTTCTTTCCTGCGTTATCCCCTGATTCTGTGGATAACCGTATTACCGCCTTTG
AGTGAGCTGATACCGCTCGCCGCAGCCGAACGACCGAGCGCAGCGAGTCAGTGAGCGAGGAAGCGGAAGAGCGCCTGATGCGG
TATTTTCTCCTTACGCATCTGTGCGGTATTTCACACCGCAATGGTGCACTCTCAGTACAATCTGCTCTGATGCCGCATAGTTAAGCCAGT
ATACACTCCGCTATCGCTACGTGACTGGGTCATGGCTGCGCCCCGACACCCGCCAACACCCGCTGACGCGCCCTGACGGGCTTG
TCTGCTCCCGGCATCCGCTTACAGACAAGCTGTGACCGTCTCCGGGAGCTGCATGTGTCAGAGGTTTTCACCGTCATCACCGAAACG
CGCGAGGCAGCTGCGGTAAAGCTCATCAGCGTGGTCGTGAAGCGATTCACAGATGTCTGCCTGTTCATCCGCGTCCAGCTCGTTGA
GTTTCTCCAGAAGCGTTAATGTCTGGCTTCTGATAAAGCGGGCCATGTTAAGGGCGGTTTTTTCCTGTTTGGTCACTGATGCCTCCGTGT
AAGGGGGATTTCTGTTCATGGGGGTAATGATACCGATGAAACGAGAGAGGATGCTCACGATACGGGTTACTGATGATGAACATGCCCG
GTTACTGGAACGTTGTGAGGGTAAACAACTGGCGGTATGGATGCGGCGGGACCAGAGAAAAATCACTCAGGGTCAATGCCAGCGCT
TCGTTAATACAGATGTAGGTGTTCCACAGGGTAGCCAGCAGCATCCTGCGATGCAGATCCGGAACATAATGGTGCAGGGCGCTGACT
TCCGCGTTTCCAGACTTTACGAAACACGGAAACCGAAGACCATTCATGTTGTTGCTCAGGTCGCAGACGTTTTGCAGCAGCAGTCGCT
TCACGTTCGCTCGCGTATCGGTGATTCATTCTGCTAACCAGTAAGGCAACCCCGCCAGCCTAGCCGGGTCCTCAACGACAGGAGCA
CGATCATGCGCACCCGTGGGGCCGCCATGCCGGCGATAATGGCCTGCTTCTCGCCGAAACGTTTGGTGGCGGGACCAGTGACGA
AGGCTTGAGCGAGGGCGTGCAAGATTCCGAATACCGCAAGCGACAGGCCGATCATCGTCGCGCTCCAGCGAAAGCGGTCCTCGC
CGAAAATGACCCAGAGCGCTGCCGGCACCTGTCCTACGAGTTGCATGATAAAGAAGACAGTCATAAGTGCGGCGACGATAGTCATG
CCCCGCGCCCACCGGAAGGAGCTGACTGGGTTGAAGGCTCTCAAGGGCATCGGTCGAGATCCCGGTGCCTAATGAGTGAGCTAAC
TTACATTAATTGCGTTGCGCTCACTGCCCGCTTTCCAGTCGGGAAACCTGTCGTGCCAGCTGCATTAATGAATCGGCCAACGCGCGG
GGAGAGGCGGTTTGCGTATTGGGCGCCAGGGTGGTTTTTCTTTTCACCAGTGAGACGGGCAACAGCTGATTGCCCTTCACCGCCTG
GCCCTGAGAGAGTTGCAGCAAGCGGTCCACGCTGGTTTGCCCCAGCAGGCGAAAATCCTGTTTGATGGTGGTTAACGGCGGGATAT
AACATGAGCTGTCTTCGGTATCGTCGTATCCCACTACCGAGATATCCGCACCAACGCGCAGCCCGGACTCGGTAATGGCGCGCATT
GCGCCCAGCGCCATCTGATCGTTGGCAACCAGCATCGCAGTGGGAACGATGCCCTCATTCAGCATTTGCATGGTTTGTTGAAAACCG
GACATGGCACTCCAGTCGCCTTCCCGTTCCGCTATCGGCTGAATTTGATTGCGAGTGAGATATTTATGCCAGCCAGCCAGACGCAGA
CGCGCCGAGACAGAACTTAATGGGCCCGCTAACAGCGCGATTTGCTGGTGACCCAATGCGACCAGATGCTCCACGCCCAGTCGC

pET21a-katE-6xHis

GTACCGTCTTCATGGGAGAAAATAATACTGTTGATGGGTGTCTGGTCAGAGACATCAAGAAATAACGCCGGAACATTAGTGCAGGCAG
CTTCCACAGCAATGGCATCCTGGTCATCCAGCGGATAGTTAATGATCAGCCCACTGACGCGTTGCGCGAGAAGATTGTGCACCGCC
GCTTTACAGGCTTCGACGCCGCTTCGTTCTACCATCGACACCACCACGCTGGCACCCAGTTGATCGGCGCGAGATTTAATCGCCGC
GACAATTTGCGACGGCGCGTGCAGGGCCAGACTGGAGGTGGCAACGCCAATCAGCAACGACTGTTTGCCCGCCAGTTGTTGTGCC
ACGCGGTTGGGAATGTAATTCAGCTCCGCCATCGCCGCTTCCACTTTTTCCCGCGTTTTCGCAGAAACGTGGCTGGCCTGGTTCACC
ACGCGGGAAACGGTCTGATAAGAGACACCGGCATACTCTGCGACATCGTATAACGTTACTGGTTTCACATTCACCACCCTGAATTGAC
TCTCTTCCGGGCGCTATCATGCCATACCGCGAAAGGTTTTGCGCCATTCGATGGTGTCCGGGATCTCGACGCTCTCCCTTATGCGACT
CCTGCATTAGGAAGCAGCCCAGTAGTAGGTTGAGGCCGTTGAGCACCGCCGCCGCAAGGAATGGTGCATGCAAGGAGATGGCGC
CCAACAGTCCCCCGGCCACGGGGCCTGCCACCATACCCACGCCGAAACAAGCGCTCATGAGCCCGAAGTGGCGAGCCCGATCT
TCCCCATCGGTGATGTCGGCGATATAGGCGCCAGCAACCGCACCTGTGGCGCCGGTGATGCCGGCCACGATGCGTCCGGCGTAG
AGGATCGAGATCTCGATCCCGCGAAATTAATACGACTCACTATAGGGGAATTGTGAGCGGATAACAATTCCCCTCTAGAAATAATTTTGT
TTAACTTTAAGAAGGAGATATACATATGTCTTCCAAGCTGGTTTTAGTGCTGAATTGCGGTAGCTCGAGCCTGAAATTCGCAATTATCGAC
GCGGTCAACGGCGAGGAGTACTTGTCTGGTCTGGCTGAGTGCTTCCATTTGCCAGAAGCCCGGATTAAGTGGAAGATGGATGGCAAC
AAGCAGGAGGCCGCGTTGGGCGCTGGGGCTGCTCATTCCGAGGCATTGAATTTTATTGTGAATACGATTCTCGCCCAAAAACCGGAA
CTGAGTGCCCAACTCACCGCCATTGGTCACCGGATCGTGCATGGGGGTGAAAAATATACAAGCTCTGTAGTGATCGATGAATCAGTGA
TCCAGGGCATCAAAGACGCCGCCTCGTTTGCACCGCTGCACAACCCAGCGCATCTGATTGGCATTGAGGAGGCGTTGAAGTCCTTT
CCTCAGCTGAAAGACAAGAATGTTGCGGTATTCGATACCGCGTTCCACCAAACAATGCCGGAGGAGAGCTACCTCTATGCACTCCCT
TACAATCTGTACAAAGAACACGGTATCCGCCGCTATGGCGCTCATGGTACAAGCCACTTCTACGTCACGCAAGAAGCAGCGAAAATG
CTCAACAAGCCTGTCGAAGAGTTAAACATTATCACGTGCCACCTTGGTAATGGGGGTCTGTTTCAGCTATCCGGAACGGCAAGTGCG
TGGATACCTCGATGGGCCTGACGCCATTGGAAGGGCTGGTAATGGGCACGCGTTCCGGTGACATTGACCCGGCTATCATCTTCCATC
TCCACGATACATTGGGTATGTCCGTGGACGCAATCAACAAGTTGTTGACAAAAGAGTCAGGGTTATTGGGTTTGACTGAGGTTACAAGT
GACTGCCGCTATGTCGAGGATAACTACGCCACGAAAGAAGATGCGAAGCGTGCTATGGACGTGTATTGTCATCGCTTGGCAAAATATAT
CGGGGCCTACACTGCACTGATGGACGGGCGGTTGGACGCAGTCGTCTTTACCGGCGGCATCGGGGAAAACGCGGCCATGGTCCG
CGAATTGAGCCTGGGTAAGCTCGGCGTTCTTGGTTTTGAAGTGGACCACGAGCGCAACTTGGCAGCTCGCTTTGGGAAGTCAGGGTT
CATCAATAAGGAAGGTACTCGTCCTGCGGTTGTAATCCCAACCAATGAGGAATTGGTAATTGCTCAAGATGCGTCGCGTTTGACCGCTC
ATCATCACCATCACCACTAAGATCCGGCTGCTAACAAAGCCCGAAAGGAAGCTGAGTTGGCTGCTGCCACCGCTGAGCAATAAC**TA**
**GCATAACCCCTTGGGGCCTCTAAACGGGTCTTGAGGGGTTTTTTG**CTGAAAGGAGGAACTATATCCGGATTGGCGAATGGGACGC
GCCCTGTAGCGGCGCATTAAGCGCGGCGGGTGTGGTGGTTACGCGCAGCGTGACCGCTACACTTGCCAGCGCCCTAGCGCCCG
CTCCTTTCGCTTTCTTCCCTTCCTTTCTCGCCACGTTCGCCGGCTTTCCCCGTCAAGCTCTAAATCGGGGGCTCCCTTTAGGGTTCCG
ATTTAGTGCTTTACGGCACCTCGACCCCAAAAAACTTGATTAGGGTGATGGTTCACGTAGTGGGCCATCGCCCTGATAGACGGTTTTTC
GCCCTTTGACGTTG
GAGTCCACGTTCTTTAATAGTGGACTCTTGTTCCAAACTGGAACAACACTCAACCCTATCTCGGTCTATTCTTTTGATTTATAAGGGATTTT
GCCGATTTCGGCCTATTGGTTAAAAAATGAGCTGATTTAACAAAAATTTAACGCGAATTTTAACAAAATATTAACGCTTACAATTTAGGTGG
CACTTTTCGGGGAAATGTGCGCGGAACCCCTATTTGTTTATTTTTCTAAATACATTCAAATATGTATCCGCTCATGAGACAATAACCCTGAT
AAATGCTTCAATAATATTGAAAAAGGAAGAGTATGAGTATTCAACATTTCCGTGTCGCCCTTATTCCCTTTTTTGCGGCATTTTGCCTTCCT
GTTTTTGCTCACCCAGAAACGCTGGTGAAAGTAAAAGATGCTGAAGATCAGTTGGGTGCACGAGTGGGTTACATCGAACTGGATCTCA
ACAGCGGTAAGATCCTTGAGAGTTTTCGCCCCGAAGAACGTTTTCCAATGATGAGCACTTTTAAAGTTCTGCTATGTGGCGCGGTATTAT
CCCGTATTGACGCCGGGCAAGAGCAACTCGGTCGCCGCATACACTATTCTCAGAATGACTTGGTTGAGTACTCACCAGTCACAGAAA
AGCATCTTACGGATGGCATGACAGTAAGAGAATTATGCAGTGCTGCCATAACCATGAGTGATAACACTGCGGCCAACTTACTTCTGACA
ACGATCGGAGGACCGAAGGAGCTAACCGCTTTTTTGCACAACATGGGGGATCATGTAACTCGCCTTGATCGTTGGGAACCGGAGCTG
AATGAAGCCATACCAAACGACGAGCGTGACACCACGATGCCTGCAGCAATGGCAACAACGTTGCGCAAACTATTAACTGGCGAACT
ACTTACTCTAGCTTCCCGGCAACAATTAATAGACTGGATGGAGGCGGATAAAGTTGCAGGACCACTTCTGCGCTCGGCCCTTCCGGC
TGGCTGGTTTATTGCTGATAAATCTGGAGCCGGTGAGCGTGGGTCTCGCGGTATCATTGCAGCACTGGGGCCAGATGGTAAGCCCTC
CCGTATCGTAGTTATCTACACGACGGGGAGTCAGGCAACTATGGATGAACGAAATAGACAGATCGCTGAGATAGGTGCCTCACTGATTA
AGCATTGGTAACTGTCAGACCAAGTTTACTCATATATACTTTAGATTGATTTAAAACTTCATTTTTAATTTAAAAGGATCTAGGTGAAGATCCTT
TTTGATAATCTCATGACCAAAATCCCTTAACGTGAGTTTTCGTTCCACTGAGCGTCAGACCCCGTAGAAAAGATCAAAGGATCTTCTTGA
GATCCTTTTTTTCTGCGCGTAATCTGCTGCTTGCAAACAAAAAAACCACCGCTACCAGCGGTGGTTTGTTTGCCGGATCAAGAGCTAC
CAACTCTTTTTCCGAAGGTAACTGGCTTCAGCAGAGCGCAGATACCAAATACTGTCCTTCTAGTGTAGCCGTAGTTAGGCCACCACTTC
AAGAACTCTGTAGCACCGCCTACATACCTCGCTCTGCTAATCCTGTTACCAGTGGCTGCTGCCAGTGGCGATAAGTCGTGTCTTACCG
GGTTGGACTCAAGACGATAGTTACCGGATAAGGCGCAGCGGTCGGGCTGAACGGGGGGTTCGTGCACACAGCCCAGCTTGGAGC
GAACGACCTACACCGAACTGAGATACCTACAGCGTGAGCTATGAGAAAGCGCCACGCTTCCCGAAGGGAGAAAGGCGGACAGGTA
TCCGGTAAGCGGCAGGGTCGGAACAGGAGAGCGCACGAGGGAGCTTCCAGGGGGAAACGCCTGGTATCTTTATAGTCCTGTCGGG
TTTCGCCACCTCTGACTTGAGCGTCGATTTTTGTGATGCTCGTCAGGGGGGCGGAGCCTATGGAAAAACGCCAGCAACGCGGCCTTT
TTACGGTTCCTGGCCTTTTGCTGGCCTTTTGCTCACATGTTCTTTCCTGCGTTATCCCCTGATTCTGTGGATAACCGTATTACCGCCTTTG
AGTGAGCTGATACCGCTCGCCGCAGCCGAACGACCGAGCGCAGCGAGTCAGTGAGCGAGGAAGCGGAAGAGCGCCTGATGCGG
TATTTTCTCCTTACGCATCTGTGCGGTATTTCACACCGCAATGGTGCACTCTCAGTACAATCTGCTCTGATGCCGCATAGTTAAGCCAGT
ATACACTCCGCTATCGCTACGTGACTGGGTCATGGCTGCGCCCCGACACCCGCCAACACCCGCTGACGCGCCCTGACGGGCTTG
TCTGCTCCCGGCATCCGCTTACAGACAAGCTGTGACCGTCTCCGGGAGCTGCATGTGTCAGAGGTTTTCACCGTCATCACCGAAACG
CGCGAGGCAGCTGCGGTAAAGCTCATCAGCGTGGTCGTGAAGCGATTCACAGATGTCTGCCTGTTCATCCGCGTCCAGCTCGTTGA
GTTTCTCCAGAAGCGTTAATGTCTGGCTTCTGATAAAGCGGGCCATGTTAAGGGCGGTTTTTTCCTGTTTGGTCACTGATGCCTCCGTGT
AAGGGGGATTTCTGTTCATGGGGGTAATGATACCGATGAAACGAGAGAGGATGCTCACGATACGGGTTACTGATGATGAACATGCCCG
GTTACTGGAACGTTGTGAGGGTAAACAACTGGCGGTATGGATGCGGCGGGACCAGAGAAAAATCACTCAGGGTCAATGCCAGCGCT
TCGTTAATACAGATGTAGGTGTTCCACAGGGTAGCCAGCAGCATCCTGCGATGCAGATCCGGAACATAATGGTGCAGGGCGCTGACT
TCCGCGTTTCCAGACTTTACGAAACACGGAAACCGAAGACCATTCATGTTGTTGCTCAGGTCGCAGACGTTTTGCAGCAGCAGTCGCT
TCACGTTCGCTCGCGTATCGGTGATTCATTCTGCTAACCAGTAAGGCAACCCCGCCAGCCTAGCCGGGTCCTCAACGACAGGAGCA
CGATCATGCGCACCCGTGGGGCCGCCATGCCGGCGATAATGGCCTGCTTCTCGCCGAAACGTTTGGTGGCGGGACCAGTGACGA
AGGCTTGAGCGAGGGCGTGCAAGATTCCGAATACCGCAAGCGACAGGCCGATCATCGTCGCGCTCCAGCGAAAGCGGTCCTCGC
CGAAAATGACCCAGAGCGCTGCCGGCACCTGTCCTACGAGTTGCATGATAAAGAAGACAGTCATAAGTGCGGCGACGATAGTCATG
CCCCGCGCCCACCGGAAGGAGCTGACTGGGTTGAAGGCTCTCAAGGGCATCGGTCGAGATCCCGGTGCCTAATGAGTGAGCTAAC
TTACATTAATTGCGTTGCGCTCACTGCCCGCTTTCCAGTCGGGAAACCTGTCGTGCCAGCTGCATTAATGAATCGGCCAACGCGCGG
GGAGAGGCGGTTTGCGTATTGGGCGCCAGGGTGGTTTTTCTTTTCACCAGTGAGACGGGCAACAGCTGATTGCCCTTCACCGCCTG
GCCCTGAGAGAGTTGCAGCAAGCGGTCCACGCTGGTTTGCCCCAGCAGGCGAAAATCCTGTTTGATGGTGGTTAACGGCGGGATAT
AACATGAGCTGTCTTCGGTATCGTCGTATCCCACTACCGAGATATCCGCACCAACGCGCAGCCCGGACTCGGTAATGGCGCGCATT
GCGCCCAGCGCCATCTGATCGTTGGCAACCAGCATCGCAGTGGGAACGATGCCCTCATTCAGCATTTGCATGGTTTGTTGAAAACCG
GACATGGCACTCCAGTCGCCTTCCCGTTCCGCTATCGGCTGAATTTGATTGCGAGTGAGATATTTATGCCAGCCAGCCAGACGCAGA
CGCGCCGAGACAGAACTTAATGGGCCCGCTAACAGCGCGATTTGCTGGTGACCCAATGCGACCAGATGCTCCACGCCCAGTCGC
GTACCGTCTTCATGGGAGAAAATAATACTGTTGATGGGTGTCTGGTCAGAGACATCAAGAAATAACGCCGGAACATTAGTGCAGGCAG
CTTCCACAGCAATGGCATCCTGGTCATCCAGCGGATAGTTAATGATCAGCCCACTGACGCGTTGCGCGAGAAGATTGTGCACCGCC
GCTTTACAGGCTTCGACGCCGCTTCGTTCTACCATCGACACCACCACGCTGGCACCCAGTTGATCGGCGCGAGATTTAATCGCCGC
GACAATTTGCGACGGCGCGTGCAGGGCCAGACTGGAGGTGGCAACGCCAATCAGCAACGACTGTTTGCCCGCCAGTTGTTGTGCC
ACGCGGTTGGGAATGTAATTCAGCTCCGCCATCGCCGCTTCCACTTTTTCCCGCGTTTTCGCAGAAACGTGGCTGGCCTGGTTCACC
ACGCGGGAAACGGTCTGATAAGAGACACCGGCATACTCTGCGACATCGTATAACGTTACTGGTTTCACATTCACCACCCTGAATTGAC
TCTCTTCCGGGCGCTATCATGCCATACCGCGAAAGGTTTTGCGCCATTCGATGGTGTCCGGGATCTCGACGCTCTCCCTTATGCGACT
CCTGCATTAGGAAGCAGCCCAGTAGTAGGTTGAGGCCGTTGAGCACCGCCGCCGCAAGGAATGGTGCATGCAAGGAGATGGCGC
CCAACAGTCCCCCGGCCACGGGGCCTGCCACCATACCCACGCCGAAACAAGCGCTCATGAGCCCGAAGTGGCGAGCCCGATCT
TCCCCATCGGTGATGTCGGCGATATAGGCGCCAGCAACCGCACCTGTGGCGCCGGTGATGCCGGCCACGATGCGTCCGGCGTAG
AGGATCGAGATCTCGATCCCGCGAAATTAATACGACTCACTATAGGGGAATTGTGAGCGGATAACAATTCCCCTCTAGAAATAATTTTGT
TTAACTTTAAGAAGGAGATATACATATGAGTCAGCATAACGAAAAAAATCCGCATCAACATCAGTCCCCACTCCATGACTCGTCAGAGG
CGAAACCTGGTATGGACAGCTTGGCTCCAGAAGACGGCAGCCACCGTCCTGCAGCGGAGCCTACACCGCCAGGCGCCCAACCA
ACAGCGCCTGGTTCATTAAAAGCGCCGGACACTCGTAACGAGAAATTAAATAGTCTGGAGGACGTGCGCAAGGGCTCAGAGAATTAT
GCCTTGACTACCAACCAGGGGGTCCGTATCGCTGACGACCAGAATAGCTTACGCGCGGGGAGTCGCGGGCCGACCCTCCTCGAA
GACTTTATCTTACGGGAAAAGATTACACACTTTGATCATGAGCGGATTCCGGAGCGCATCGTACACGCACGGGGGAGTGCAGCCCAC
GGTTACTTTCAACCTTACAAAAGCTTGTCAGATATCACCAAAGCTGACTTCTTATCTGATCCAAATAAAATTACACCGGTATTCGTGCGGT
TCAGTACGGTCCAGGGTGGCGCAGGTTCGGCAGATACAGTACGGGATATCCGGGGTTTTGCGACAAAATTTTATACGGAGGAGGGGA
TTTTCGATCTGGTCGGCAATAATACGCCTATTTTCTTTATCCAGGACGCGCATAAGTTCCCTGACTTTGTGCATGCAGTAAAACCAGAAC

CTCATTGGGCTATTCCACAGGGCCAGAGTGCTCACGATACTTTTTGGGATTACGTCTCTTTGCAGCCTGAAACACTTCACAATGTGATGT
GGGCAATGAGCGATCGCGGTATTCCTCGGAGCTACCGGACCATGGAGGGCTTCGGGATCCATACATTCCGGCTGATCAATGCCGAA
GGCAAGGCCACGTTTGTACGCTTTCATTGGAAACCTCTGGCTGGGAAAGCATCGTTAGTTTGGGACGAGGCGCAGAAGTTAACAGGG
CGGGACCCTGATTTTCACCGCCGGGAACTCTGGGAGGCTATTGAGGCTGGGGATTTCCCTGAATACGAACTCGGCTTTCAGCTTATT
CCGGAGGAAGATGAATTTAAATTTGACTTCGATCTCTTAGATCCAACGAAGCTGATCCCGGAAGAATTGGTCCCGGTTCAGCGCGTGG
GCAAGATGGTTCTCAATCGCAACCCAGATAACTTCTTCGCGGAAAACGAACAAGCAGCTTTTCATCCTGGGCACATCGTCCCGGGGC
TGGACTTTACAAATGACCCGCTTTTACAAGGGCGCTTATTTCTTACACCGACACGCAAATCTCACGTCTCGGTGGGCCAAACTTCCAT
GAGATCCCTATCAATCGGCCTACATGTCCGTATCACAACTTCCAACGTGACGGCATGCACCGTATGGGCATTGATACCAATCCGGCAA
ATTATGAACCGAACTCAATCAATGACAACTGGCCGCGCGAAACCCCACCGGGCCCAAAGCGGGGGGGTTTCGAGTCATACCAAGA
ACGGGTTGAGGGTAATAAAGTGCGCGAGCGGAGCCCTTCGTTTGGTGAATATTATTCCCATCCGCGCCTTTTTTGGCTCAGTCAAACT
CCATTTGAGCAGCGGCACATCGTTGACGGGTTCAGCTTTGAGTTATCTAAAGTCGTCCGGCCGTACATTCGCGAACGTGTCGTTGATC
AGCTCGCTCATATCGATCTCACCTTAGCCCAGGCTGTGGCCAAGAACCTTGGTATCGAACTTACCGATGACCAATTGAACATCACTCC
TCCACCGGATGTGAACGGCCTGAAGAAGGATCCTAGTTTAAGCTTATACGCGATTCCTGATGGTGACGTGAAGGGCCGGGTCGTGGC
CATTTTGCTTAACGACGAAGTGCGGTCAGCTGACCTCCTTGCTATTTTAAAGGCACTTAAAGCTAAAGGGGTACATGCCAAGCTGTTGT
ACTCCCGTATGGGGGAGGTGACAGCTGACGATGGTACCGTGTTGCCAATCGCAGCTACATTTGCAGGCGCACCTAGTCTTACTGTGG
ATGCCGTTATCGTCCCTTGTGGCAACATTGCCGATATCGCTGACAACGGGGACGCGAATTATTACCTCATGGAAGCTTATAAGCACCTT
AAACCTATTGCTTTAGCAGGGGACGCACGCAAATTTAAGGCGACTATCAAGATCGCCGACCAGGGTGAGGAAGGTATCGTCGAGGC
AGACTCCGCCGATGGGTCATTCATGGACGAATTGCTGACCCTGATGGCGGCGCACCGTGTGTGGAGCCGTATTCCGAAGATCGACA
AAATTCCTGCTCATCATCACCATCACCACTAAGATCCGGCTGCTAACAAAGCCCGAAAGGAAGCTGAGTTGGCTGCTGCCACCGCTG
AGCAATAAC**TAGCATAACCCCTTGGGGCCTCTAAACGGGTCTTGAGGGGTTTTTTG**CTGAAAGGAGGAACTATATCCGGATTGGCG
AATGGGACGCGCCCTGTAGCGGCGCATTAAGCGCGGCGGGTGTGGTGGTTACGCGCAGCGTGACCGCTACACTTGCCAGCGCC
CTAGCGCCCGCTCCTTTCGCTTTCTTCCCTTCCTTTCTCGCCACGTTCGCCGGCTTTCCCCGTCAAGCTCTAAATCGGGGGCTCCCT
TTAGGGTTCCGATTTAGTGCTTTACGGCACCTCGACCCCAAAAAACTTGATTAGGGTGATGGTTCACGTAGTGGGCCATCGCCCTGATA
GACGGTTTTTCGCCCTTTGACGTTG

#### Table S10: Amino acid sequences of proteins (excluding PURE proteins)

| Protein | Amino Acid Sequence | Molecular Weight |
|---|---|---|
| mCherry | MVSKGEEDNMAIIKEFMRFKVHMEGSVNGHEFEIEGEGEGRPYEGTQTAKLKVTK<br>GGPLPFAWDILSPQFMYGSKAYVKHPADIPDYLKLSFPEGFKWERVMNFEDGGVV<br>TVTQDSSLQDGEFIYKVKLRGTNFPSDGPVMQKKTMGWEASSERMYPEDGALKG<br>EIKQRLKLKDGGHYDAEVKTTYKAKKPVQLPGAYNVNIKLDITSHNEDYTIVEQYER<br>AEGRHSTGGMDELYK | 26.7 kDa |
| Pyruvate oxidase (Pox5) | MVMKQTKQTNILAGAAVIKVLEAWGVDHLYGIPGGSINSIMDALSAERDRIHYIQVR<br>HEEVGAMAAAADAKLTGKIGVCFGSAGPGGTHLMNGLYDAREDHVPVLALIGQF<br>GTTGMNMDTFQEMNENPIYADVADYNVTAVNAATLPHVIDEAIRRAYAHQGVAVV<br>QIPVDLPWQQIPAEDWYASANSYQTPLLPEPDVQAVTRLTQTLLAAERPLIYYGIGA<br>RKAGKELEQLSKTLKIPLMSTYPAKGIVADRYPAYLGSANRVAQKPANEALAQADVV<br>LFVGNNYPFAEVSKAFKNTRYFLQIDIDPAKLGKRHKTDIAVLADAQKTLAAILAQVS<br>ERESTPWWQANLANVKNWRAYLASLEDKQEGPLQAYQVLRAVNKIAEPDAIYSID<br>VGDINLNANRHLKLTPSNRHITSNLFATMGVGIPGAIAAKLNYPERQVFNLAGDGG<br>ASMTMQDLATQVQYHLPVINVVFTNCQYGFIKDEQEDTNQNDFIGVEFNDIDFSKI<br>ADGVHMQAFRVNKIEQLPDVFEQAKAIAQHEPVLIDAVITGDRPLPAEKLRLDSATS<br>SAADIEAFKQRYEAQDLQPLSTYLKQFGLDDLQHQIGQGGF | 66.1 kDa |
| Acetate kinase (AckA) | MSSKLVLVLNCGSSSLKFAIIDAVNGEEYLSGLAECFHLPEARIKWKMDGNKQEAA<br>LGAGAAHSEALNFIVNTILAQKPELSAQLTAIGHRIVHGGEKYTSSVVIDESVIQGIK<br>DAASFAPLHNPAHLIGIEEALKSFPQLKDKNVAVFDTAFHQTMPEESYLYALPYNLY<br>KEHGIRRYGAHGTSHFYVTQEAAKMLNKPVEELNIITCHLGNGGSVSAIRNGKCVD<br>TSMGLTPLEGLVMGTRSGDIDPAIIFHLHDTLGMSVDAINKLLTKESGLLGLTEVTSD<br>CRYVEDNYATKEDAKRAMDVYCHRLAKYIGAYTALMDGRLDAVVFTGGIGENAAM<br>VRELSLGKLGVLGFEVDHERNLAARFGKSGFINKEGTRPAVVIPTNEELVIAQDASR<br>LTA | 43.2 kDa |
| Catalase (KatE) | MSQHNEKNPHQHQSPLHDSSEAKPGMDSLAPEDGSHRPAAEPTPPGAQPTAPG<br>SLKAPDTRNEKLNSLEDVRKGSENYALTTNQGVRIADDQNSLRAGSRGPTLLEDFI<br>LREKITHFDHERIPERIVHARGSAAHGYFQPYKSLSDITKADFLSDPNKITPVFVRFS<br>TVQGGAGSADTVRDIRGFATKFYTEEGIFDLVGNNTPIFFIQDAHKFPDFVHAVKPE<br>PHWAIPQGQSAHDTFWDYVSLQPETLHNVMWAMSDRGIPRSYRTMEGFGIHTFR<br>LINAEGKATFVRFHWKPLAGKASLVWDEAQKLTGRDPDFHRRELWEAIEAGDFPE<br>YELGFQLIPEEDEFKFDFDLLDPTKLIPEELVPVQRVGKMVLNRNPDNFFAENEQA<br>AFHPGHIVPGLDFTNDPLLQGRLFSYTDTQISRLGGPNFHEIPINRPTCPYHNFQR<br>DGMHRMGIDTNPANYEPNSINDNWPRETPPGPKRGGFESYQERVEGNKVRERS<br>PSFGEYYSHPRLFWLSQTPFEQRHIVDGFSFELSKVVRPYIRERVVDQLAHIDLTLA<br>QAVAKNLGIELTDDQLNITPPPDVNGLKKDPSLSLYAIPDGDVKGRVVAILLNDEVR<br>SADLLAILKALKAKGVHAKLLYSRMGEVTADDGTVLPIAATFAGAPSLTVDAVIVPCG<br>NIADIADNGDANYYLMEAYKHLKPIALAGDARKFKATIKIADQGEEGIVEADSADGS<br>FMDELLTLMAAHRVWSRIPKIDKIPA | 84.1 kDa |

#### Table S11: Buffers for protein purification

| Compound | Stock Solution (mM) | Buffer A (mM) | Buffer B (mM) | Buffer HT (mM) | Stock 60 (mM) | Stock 30 (mM) |
|---|---|---|---|---|---|---|
| HEPES | 1000 | 50 | 50 | 50 | 50 | 50 |
| Magnesium chloride | 1000 | 10 | 10 | 10 | 10 | 10 |
| Potassium chloride | 2000 | 0 | 100 | 100 | 100 | 100 |
| Ammonium chloride | - | 1000 | 0 | 0 | 0 | 0 |
| Imidazole | - | 0 | 500 | 0 | 0 | 0 |
| Glycerol | 100 | 0 | 0 | 0 | 60% | 30% |
| TCEP* | 500 | 1 | 1 | 1 | 1 | 1 |
| **Notes** | | **pH- 7.6, KOH** | **pH- 7.6, HCl** | | | |

*TCEP was added to the buffers just before use.

#### Table S12: Buffers for ribosome purification

| Compound | Stock Solution (mM) | Ribosome Buffer A (mM) | Ribosome Buffer B (mM) |
|---|---|---|---|
| HEPES | 1000 | 20 | 20 |
| Magnesium glutamate | 1000 | 6 | 6 |
| Ammonium acetate | 1000 | 30 | 400 |
| DTT* | 1000 | 1 | 1 |
| **Notes** | | **pH- 7.6, KOH** | **pH- 7.6, KOH** |

*DTT was added to the buffers just before use.

#### Table S13: Energy solution composition

| Component | Stock concentration [mM] | Concentration of components in reaction [mM] | Concentration in 4x Energy solution [mM] |
|---|---|---|---|
| **HEPES** | 1000 | 50 | **200** |
| **ATP** | 100 | 2 | **8** |
| **GTP** | 100 | 2 | **8** |
| **CTP** | 100 | 1 | **4** |
| **UTP** | 100 | 1 | **4** |
| **tRNA [mg/mL]** | 215 | 3.5 | **14** |
| **TCEP** | 500 | 1 | **4** |
| **Folinic acid** | 34 | 0.02 | **0.08** |
| **Spermidine** | 500 | 2 | **8** |
| **Amino Acid solution** | 6 | 0.3 | **1.2** |
